# RNA inhibits dMi-2/CHD4 chromatin binding and nucleosome remodelling

**DOI:** 10.1101/2021.06.03.446896

**Authors:** Ikram Ullah, Clemens Thoelken, Yichen Zhong, Mara John, Oliver Rossbach, Jonathan Lenz, Markus Großringer, Andrea Nist, Thorsten Stiewe, Roland Hartmann, Olalla Vazquez, Ho-Ryung Chung, Joel P Mackay, Alexander Brehm

**Affiliations:** Institute of Molecular Biology and Tumor Research, Biomedical Research Center, Philipps-University, Marburg, Germany; Institute for Medical Bioinformatics and Biostatistic, Philipps-University, Marburg, Germany; School of Life and Environmental Sciences, University of Sydney, NSW 2006, Australia; Institute of Biochemistry, Department of Biology and Chemistry, Justus Liebig University Giessen, Giessen, Germany; Institute of Pharmaceutical Chemistry, Philipps-University, Marburg, Germany; Genomics Core Facility, Institute of Molecular Oncology, Member of the German Center for Lung Research (DZL), Philipps-University, Marburg, Germany; Faculty of Chemistry, Philipps-University, Hans-Meerwein-Strasse 4, 35043 Marburg, Germany

**Keywords:** Chromatin, ATP-dependent chromatin remodelling, RNA, NuRD, iCLIP, gene regulation

## Abstract

The ATP-dependent nucleosome remodeller Mi-2/CHD4 broadly modulates epigenetic landscapes to repress transcription and to maintain genome integrity. Here we use individual nucleotide resolution crosslinking and immunoprecipitation (iCLIP) to show that *Drosophila* Mi-2 associates with thousands of mRNA molecules *in vivo*. Biochemical data reveal that recombinant dMi-2 preferentially binds to G-rich RNA molecules using two intrinsically disordered regions of previously undefined function. Pharmacological inhibition of transcription and RNase digestion approaches establish that RNA inhibits the association of dMi-2 with chromatin. We also show that RNA inhibits dMi-2-mediated nucleosome mobilization by competing with the nucleosome substrate. Importantly, this activity is shared by CHD4, the human homolog of dMi-2, strongly suggesting that RNA-mediated regulation of remodeller activity is an evolutionary conserved mechanism. Our data support a model in which RNA serves to protect actively transcribed regions of the genome from dMi-2/CHD4- mediated establishment of repressive chromatin structures.

## Introduction

Although a growing number of chromatin regulating proteins have been demonstrated to bind RNA, the functional implications of these interactions are largely not understood (Castello et al., 2016; He et al., 2016; Hendrickson et al., 2016). In only a few cases has the functional relationship between such proteins and their cognate RNA molecules been studied in detail. One example is the Polycomb Repressive Complex 2 (PRC2), which catalyses the repressive trimethylation of lysine 27 of histone H3 (H3K27me3). PRC2 binds RNA in a promiscuous manner while displaying a preference for G-quadruplex-forming and G-tract containing RNAs (Wang et al., 2017). Interaction with RNA can affect several PRC2 functions, sometimes with seemingly opposite outcomes. For example, RNA binding is important genome wide for association of PRC2 with PRC2-repressed targets, implicating RNA in PRC2 recruitment (Long et al., 2020). At the same time, nascent RNA evicts PRC2 from active genes (Beltran et al., 2019; Beltran et al., 2016; Long et al., 2020). Moreover, RNA binding to an allosteric regulatory site of the methyltransferase subunit Enhancer of zeste homolog 2 (EZH2) inhibits PRC2 enzymatic activity (Zhang et al., 2019). Thus, RNA binding can lead to multifaceted effects on PRC2 function.

RNA-mediated regulation of PRC2 impacts H3K27me3 levels within PRC2-repressed regions which are essential for stable transcriptional repression of lineage-inappropriate genes and proper differentiation of ES cells (Long et al., 2020; Shan et al., 2017). H3K27 methylation is facilitated by the Nucleosome Remodelling and Deacetylation (NuRD) complex (Bracken et al., 2019; Reynolds et al., 2012b). In ES cells, NuRD removes the acetyl group from acetylated lysine 27 of histone H3 (H3K27ac), a prerequisite for PRC2 chromatin association and H3K27 methylation. Moreover, NuRD antagonizes activating SWI/SNF remodelling complexes and increases nucleosome density (Kreher et al., 2017; Liang et al., 2017; Moshkin et al., 2012). These NuRD activities are believed to further assist PRC-mediated chromatin compaction. Like PRC2, NuRD is important for ES cell differentiation (Bornelov et al., 2018; Bracken et al., 2019; Burgold et al., 2019; Ragheb et al., 2020). It is not known if NuRD function, like PRC2 function, is modulated by RNA and whether RNA can coordinately regulate both chromatin regulatory complexes.

CHD4, the ATPase subunit of NuRD, has been identified as an RNA binding protein in several large-scale screens but the consequences of RNA binding on CHD4 function have not been systematically analysed (Castello et al., 2016; He et al., 2016; Hendrickson et al., 2016). CHD4 binds PAPAS, a long non-coding RNA derived from rRNA promoters and this interaction has been suggested to aid association of NuRD with rRNA genes (Zhao et al., 2018). Our own work on the recruitment of the *Drosophila* CHD4 homologue, dMi-2, to active heat shock (HS) genes indicates that dMi-2 binding to nascent HS gene transcripts might contribute to its association with HS gene bodies (Mathieu et al., 2012; Murawska et al., 2011). However, a comprehensive view of CHD4/dMi-2 RNA binding and its functional consequences is lacking.

Here, we use iCLIP and fRIP to identify dMi-2 associated RNAs *in vivo*. We show that dMi-2 binds to thousands of mRNAs and exhibits a preference for their 3’ regions. Using a combination of bioinformatic analyses, biochemical approaches and chromatin immunoprecipitation following pharmacological inhibition of RNA polymerase II, we reveal that RNA restricts dMi-2 association with chromatin. We establish that dMi-2 displays broad RNA binding activity with a preference for G-rich RNA sequences *in vitro*. We map RNA binding activity to two intrinsically disordered regions (IDRs) that are conserved between fly and human. Finally, we establish that G-rich RNA inhibits nucleosome remodelling by both *Drosophila* Mi-2 and human CHD4 and uncover the molecular mechanism of this inhibition. We propose that RNA antagonizes both dMi-2/CHD4 and PRC2 in a concerted manner to safeguard actively transcribed genes from the establishment of repressive chromatin structures.

## Results

### dMi-2 associates with thousands of mRNAs in vivo

To investigate the *in vivo* association of dMi-2 with RNA in an unbiased manner we used individual nucleotide resolution <u>c</u>rosslinking and immuno<u>p</u>recipitation (iCLIP2). We first tested if dMi-2 can be photo-crosslinked to RNA *in vivo*. S2 cells expressing GFP-tagged dMi-2 (dMi-2-GFP) from the endogenous dMi-2 locus (Lenz et al., 2021) were irradiated with UV (254 nm) to photo-crosslink RNA and proteins. Extracts from these cells were partially digested with RNase I to shorten crosslinked RNA molecules and immunoprecipitated with GFP-Trap beads. Crosslinked RNAs were then labelled with ^32^P at their 5’ ends and protein/[^32^P]-RNA photo-adducts were visualized by autoradiography following SDS-PAGE and transfer to a membrane **(Figure 1A)**. No significant radioactive signal was detected in samples derived from UV-treated S2 control cells which did not express GFP-tagged dMi-2 (upper panel: lanes 2 to 4) or in samples derived from S2 cells expressing dMi-2-GFP that had not been exposed to UV (lanes 8 to 10). In contrast, robust signals were obtained in samples derived from UV-crosslinked, dMi-2-GFP expressing cells treated with low or intermediate RNase concentrations (lanes 6 and 7). These signals largely disappeared when the sample was treated with the highest RNase concentration (lane 5). The radioactive signal was concentrated in the apparent molecular weight range of 250-300 kDa. This corresponds well to the apparent molecular weight of dMi-2-GFP as determined by Western blot (lower panel). These results demonstrate that dMi-2-GFP can be efficiently photo-crosslinked to RNA and, therefore, strongly imply that dMi-2 is closely associated with RNA *in vivo*.

**Figure 1:**
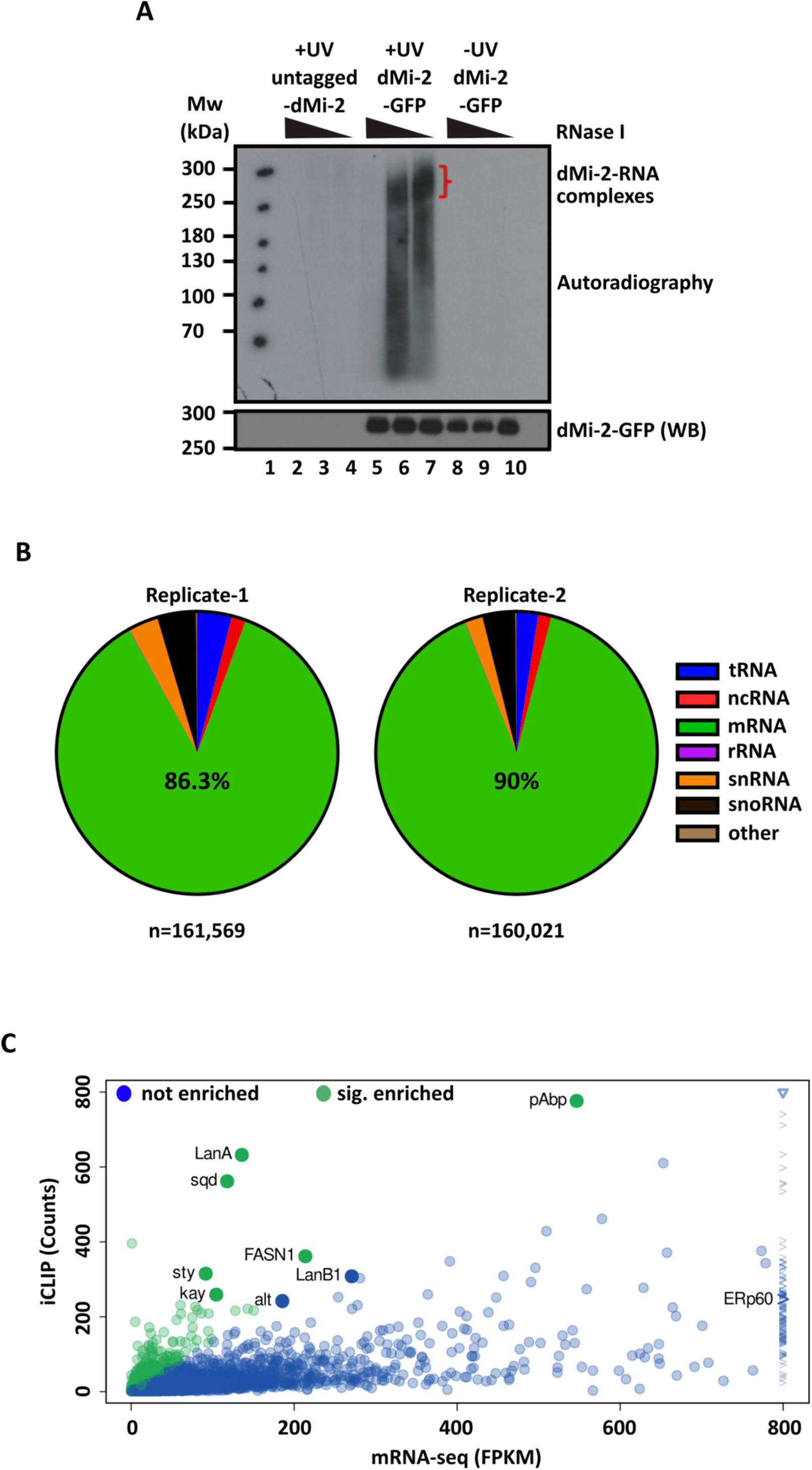
dMi-2 associates with thousands of mRNAs *in vivo*. **(A)** dMi-2 binds RNA *in vivo*. Autoradiograph (upper panel) and dMi-2 Western blot (WB) (lower panel). S2 cells expressing dMi-2-GFP (dMi-2-GFP) or control S2 cells (untagged-dMi-2) were UV-irradiated (+UV) or left untreated (-UV) as indicated. Samples were treated with decreasing concentrations of RNase I as shown on top. Red closing bracket indicates dMi-2-RNA complexes. Protein molecular weights are indicated on the left. **(B)** The distribution of different types of RNAs in replicates-1 and -2 are represented as pie charts, the total number of reads (n) in each replicate is shown below. **(C)** Correlation of the number of crosslinks per gene (iCLIP2 counts, y-axis) with RNA-Seq abundances (RNA-seq, FPKM, x-axis). We used DESeq2 to determine robustly and significantly enriched mRNAs (green dots) across the two biological iCLIP replicates and three RNA-Seq replicates (fold difference > 4 and p < 0.05). RNAs chosen for validation by fRIP-qPCR are indicated.

We then repeated the above procedure with dMi-2-GFP expressing cells and control cells not expressing dMi-2-GFP. We excized membrane slices to recover protein/[^32^P]-RNA photo-adducts in the 250-300 kDa range for further processing **(Supplementary Figure 1A)**. Equivalent membrane regions were excized for samples derived from UV-crosslinked cells expressing dMi-2-GFP (two biological replicates) and samples derived from UV-crosslinked control cells not expressing dMi-2-GFP (control; two biological replicates). dMi-2 exists in two multi-subunit protein complexes: dNuRD and dMec (Kunert et al., 2009). Given that dMi-2 is by far the largest subunit in both complexes, restricting the analysis to photo-adducts in the 250-300 kDa range ensured that RNAs identified by iCLIP were most likely directly bound to dMi-2 as opposed to other dNuRD or dMec subunits (Lenz et al., 2021).

We processed the four membrane slices according to the iCLIP2 library preparation protocol (Buchbender et al., 2020) and DNA deep sequencing resulted in a combined total of 5.03 million reads. Of these 41.8% were derived from dMi-2-GFP replicate-1, 31.9% from dMi-2- GFP replicate-2 and less than 2% from each of the two control replicates **(Supplementary Figure 1B)**. The remaining reads were missing barcode sequences and could not be assigned to any replicate. 161,569 reads (dMi-2-GFP replicate-1) and 160,021 reads (dMi-2-GFP replicate-2), respectively, uniquely mapped to RNAs encoded by the *Drosophila melanogaster* genome (*dm6*). These RNAs included tRNAs, rRNAs, snRNAs, snoRNAs and other noncoding RNAs **(Figure 1B)**. However, the majority mapped to mRNAs (86.3% dMi-2-GFP replicate-1, 90.0% dMi-2-GFP replicate-2). Furthermore, the data sets of the two replicates were highly correlated (squared Pearson correlation coefficient (R^2^) = 0.97; **(Supplementary Figure 1C)**.

We defined dMi-2 associated RNAs as RNAs exhibiting five or more crosslinks in each replicate. 5471 RNAs fulfilled this criterion. Given dMi-2’s established role in the regulation of protein coding genes, we focused our analysis on protein coding RNAs. We asked if the propensity of dMi-2 to crosslink to a given mRNA was a function of the expression level of that mRNA. We plotted the number of iCLIP2 crosslinks to a given mRNA against the expression level of that mRNA as determined by RNA-seq (Lenz et al., 2021) (**Figure 1C**, see Methods for details). This demonstrated that there was no strict linear correlation between RNA expression level and the number of dMi-2 iCLIP2 crosslinks. We identified 580 transcripts displaying an enrichment of dMi-2 crosslinks (see Methods for details; **Figure 1C**, green dots, **Table 1**).

**Table 1:**
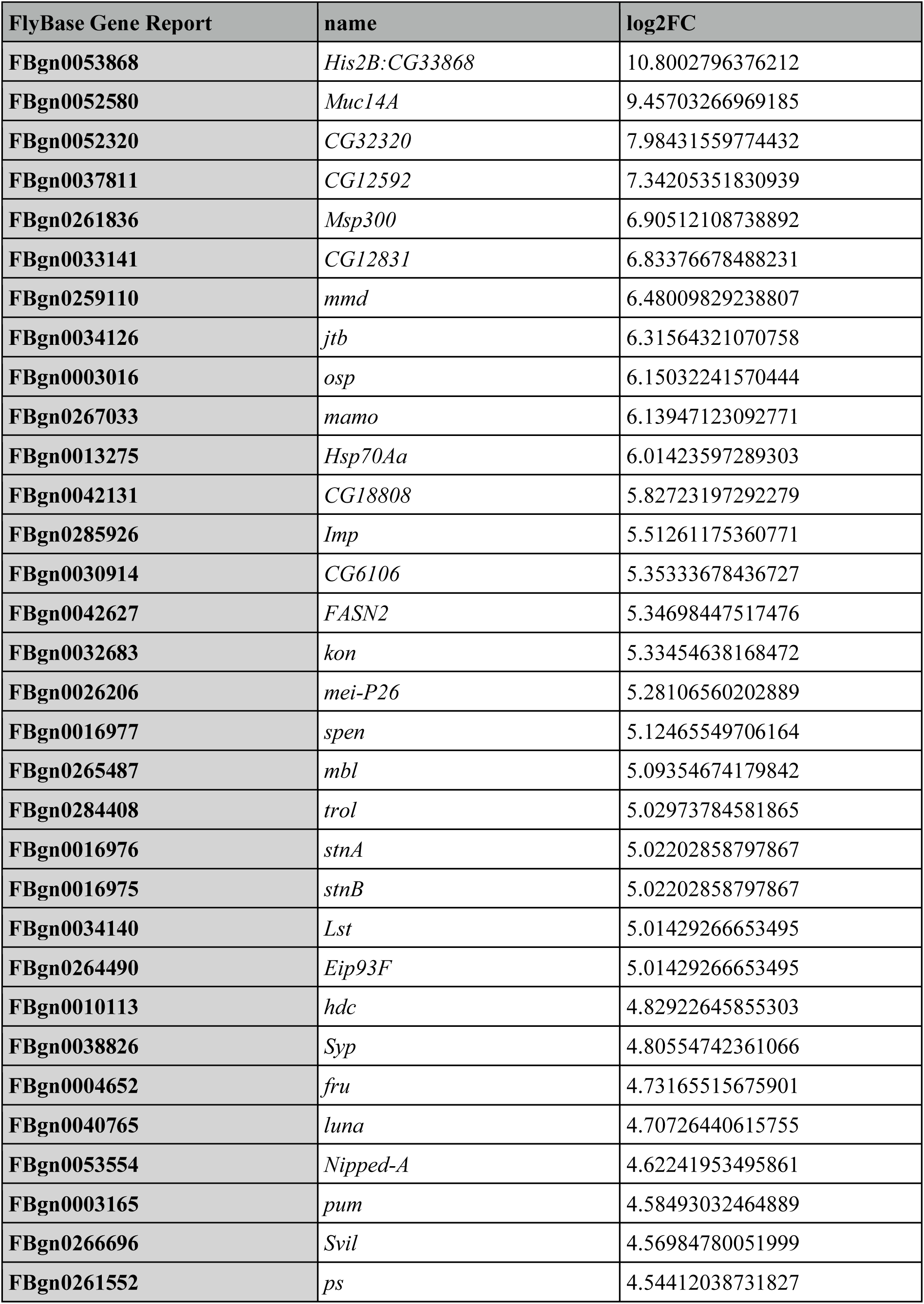
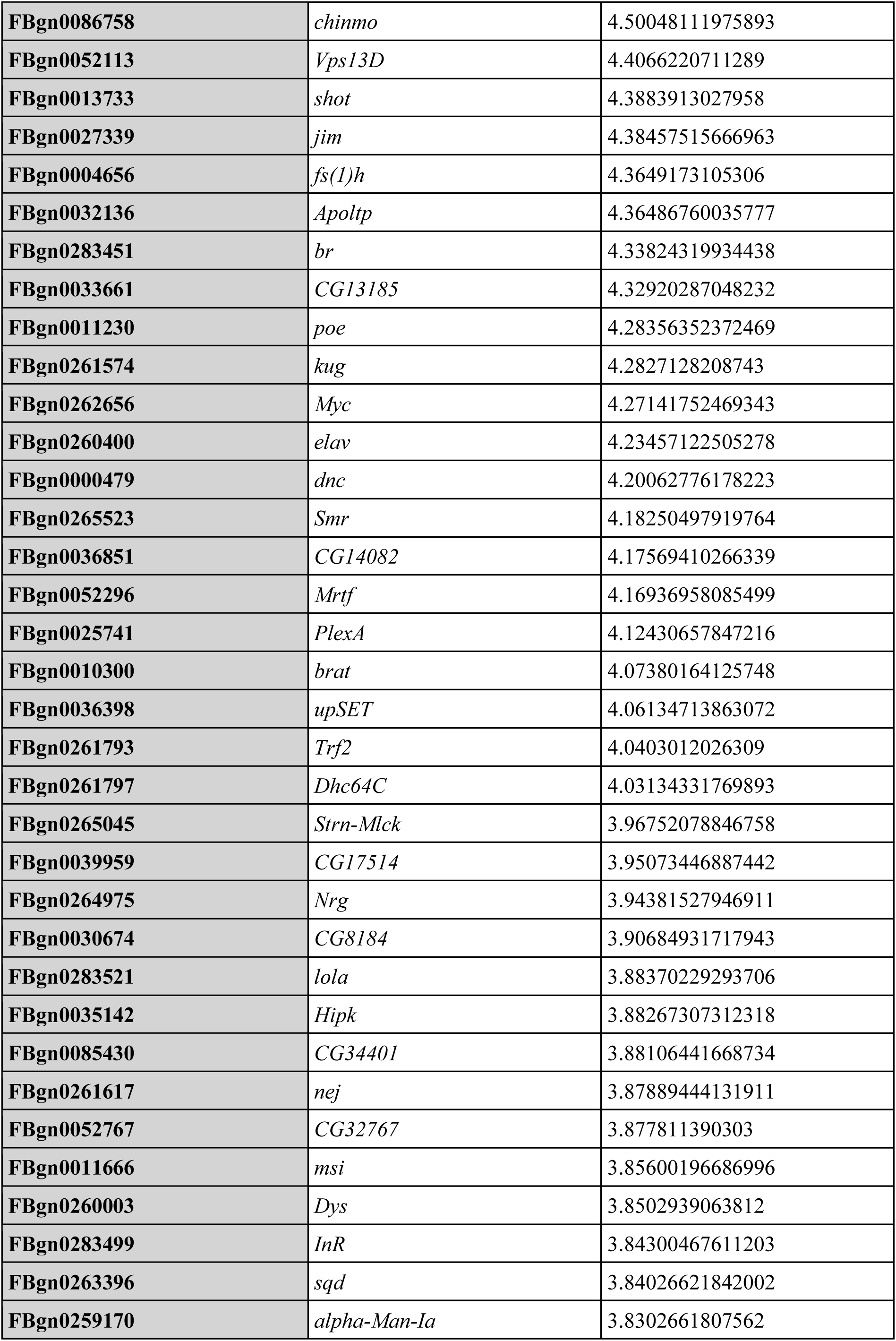
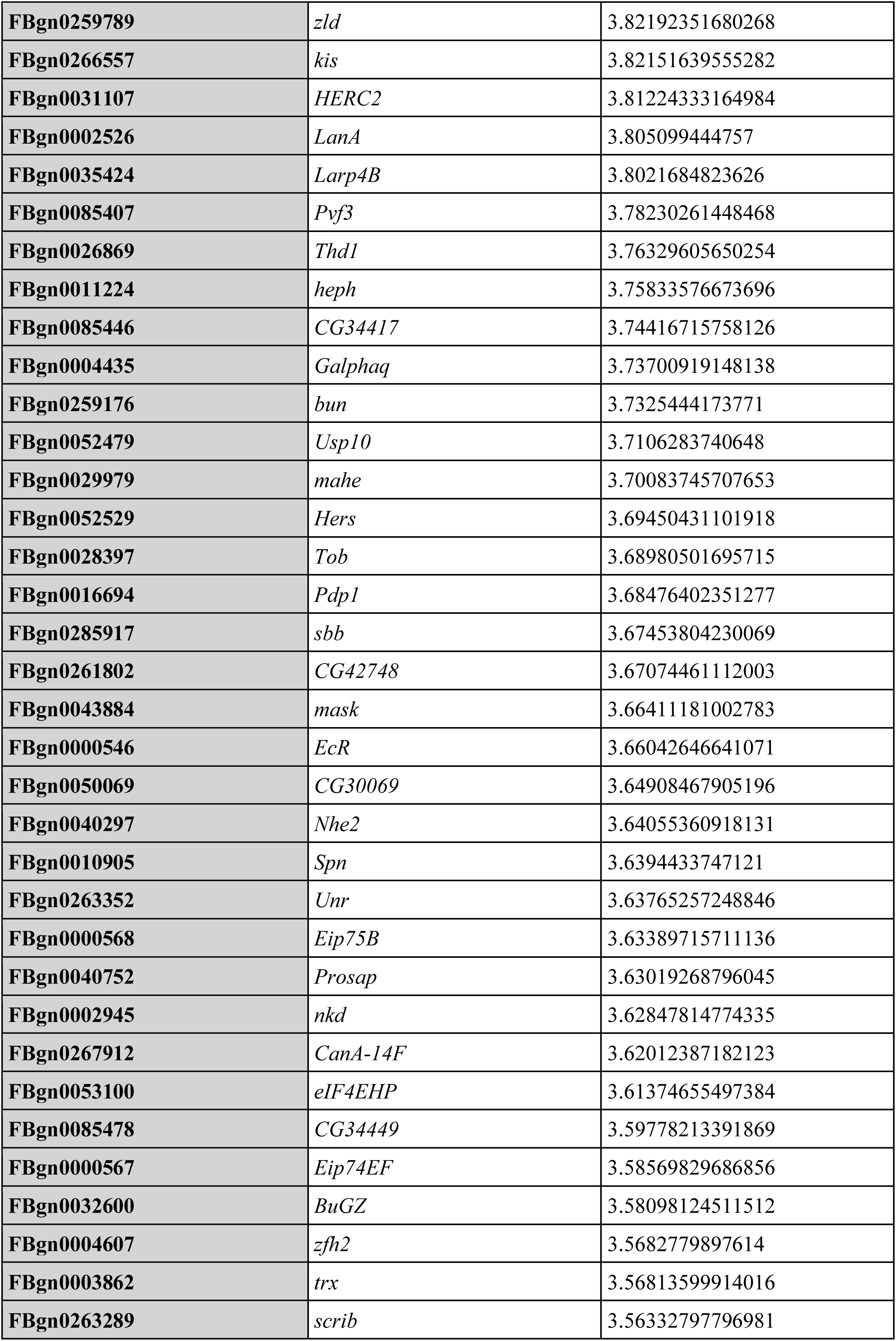
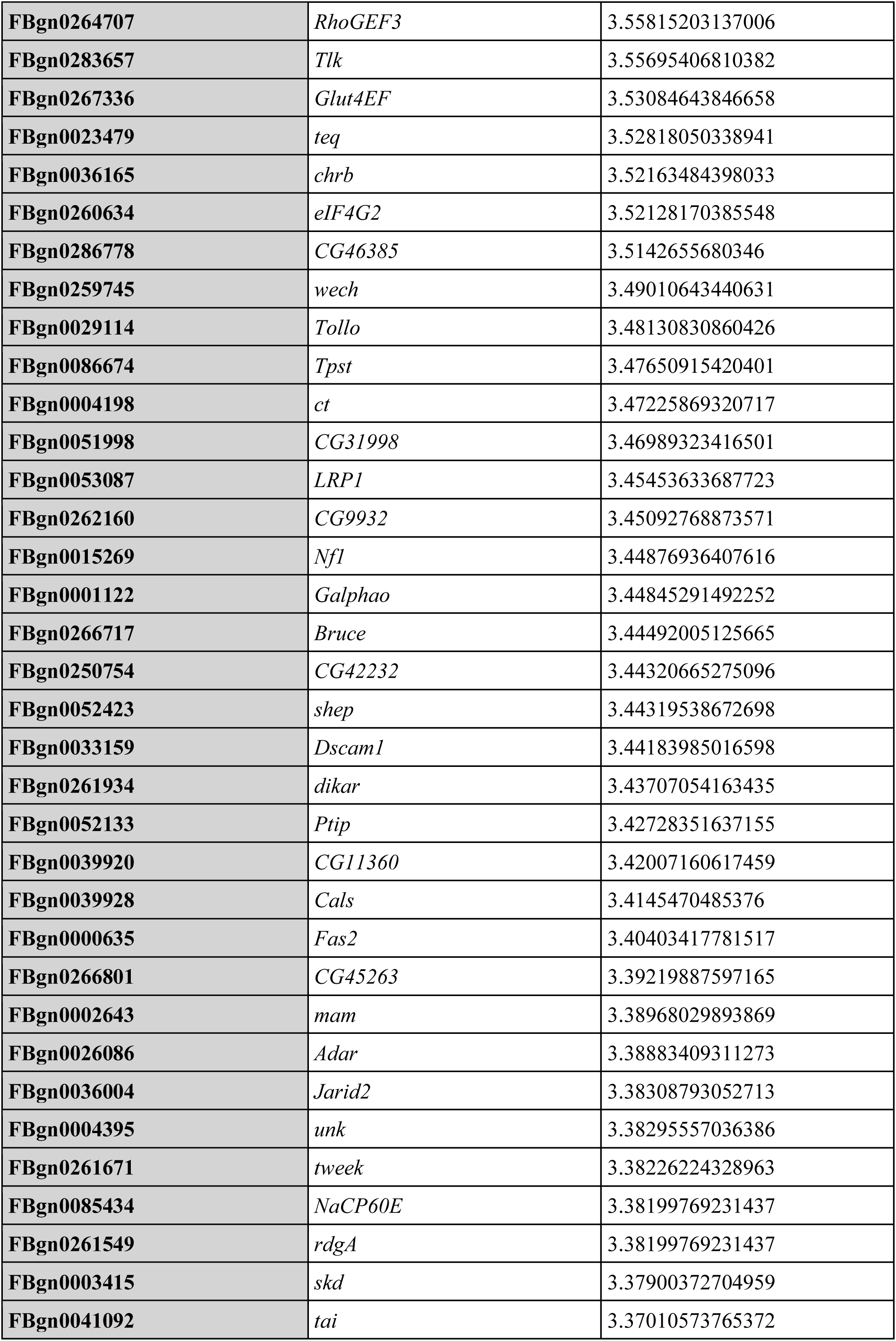
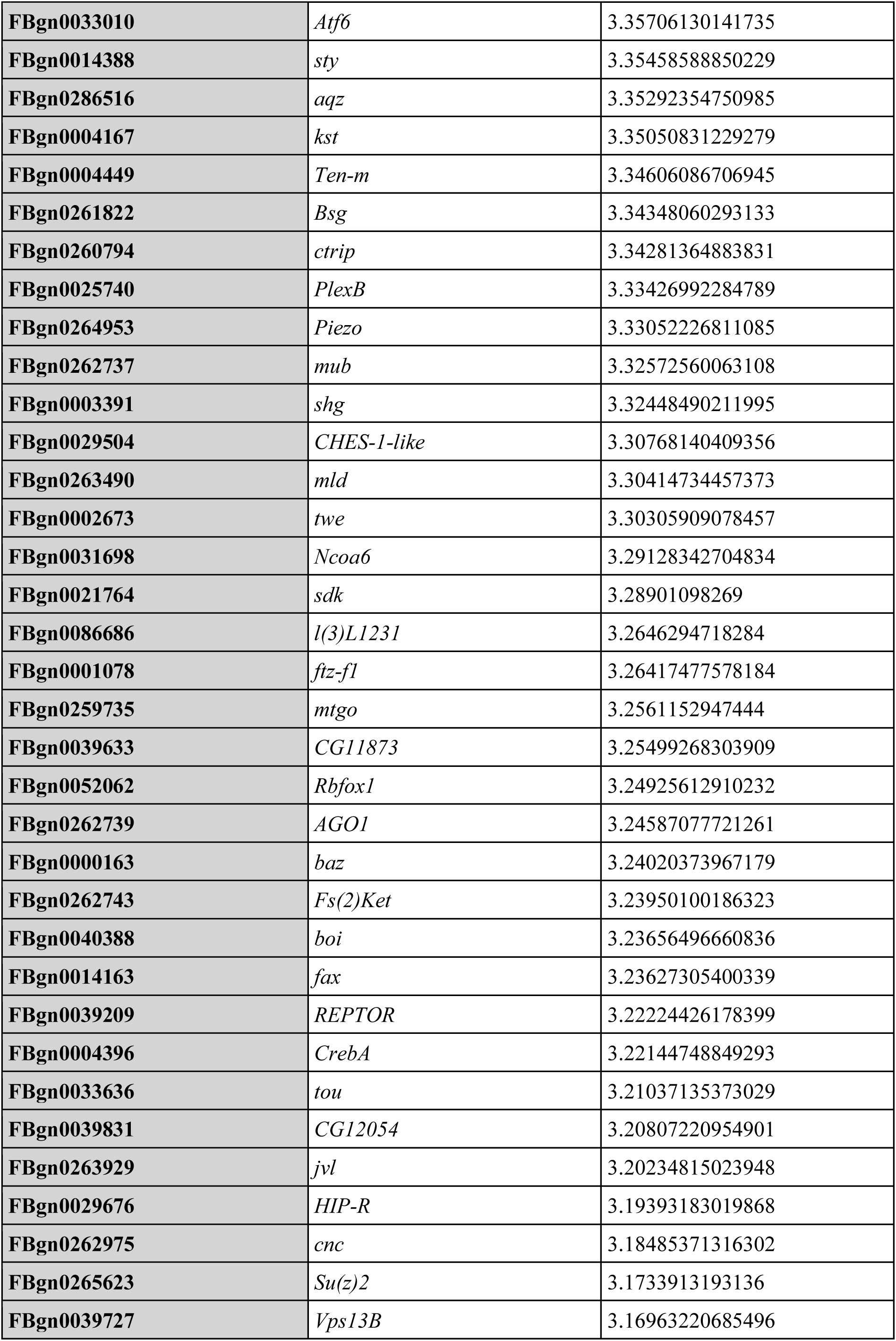
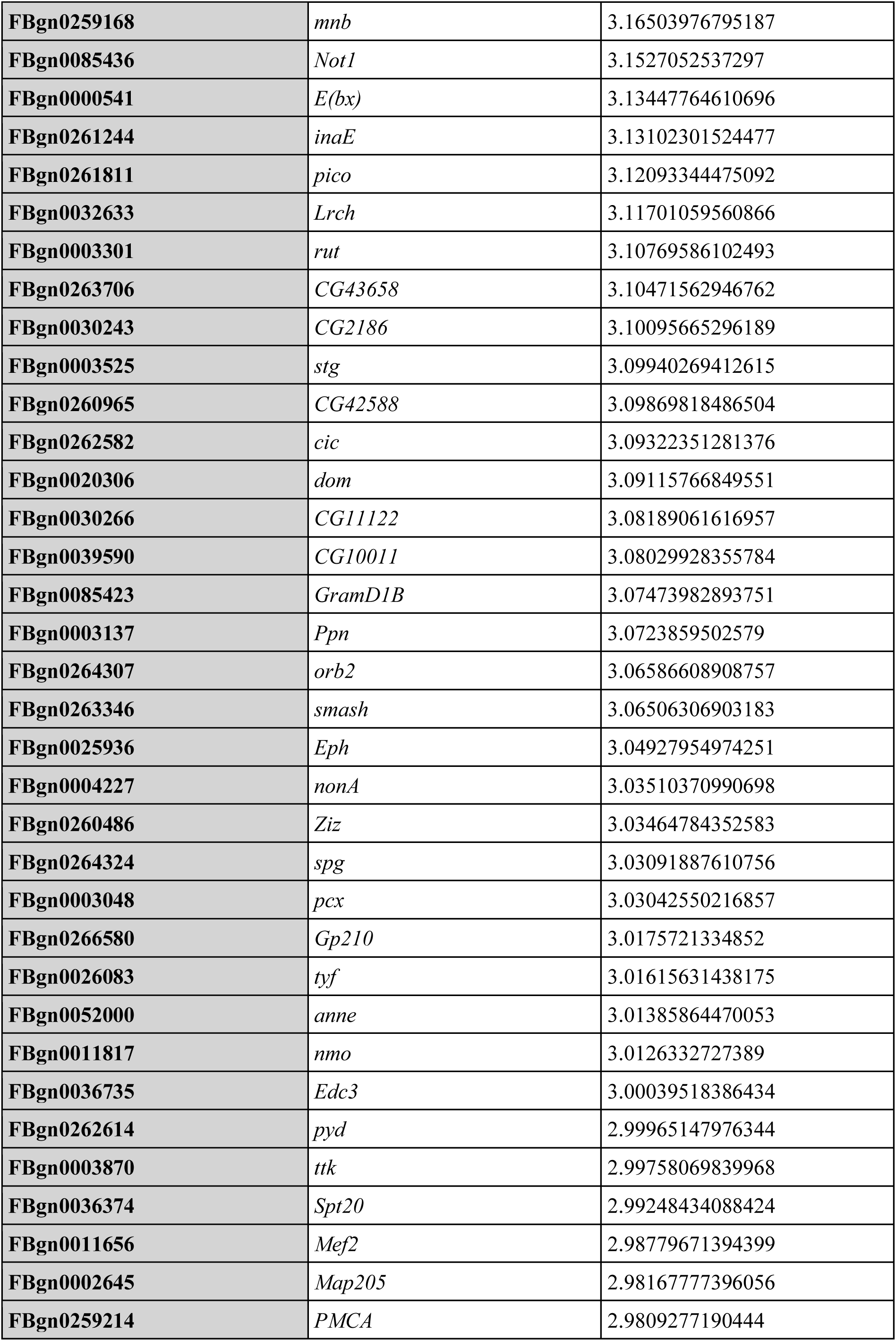
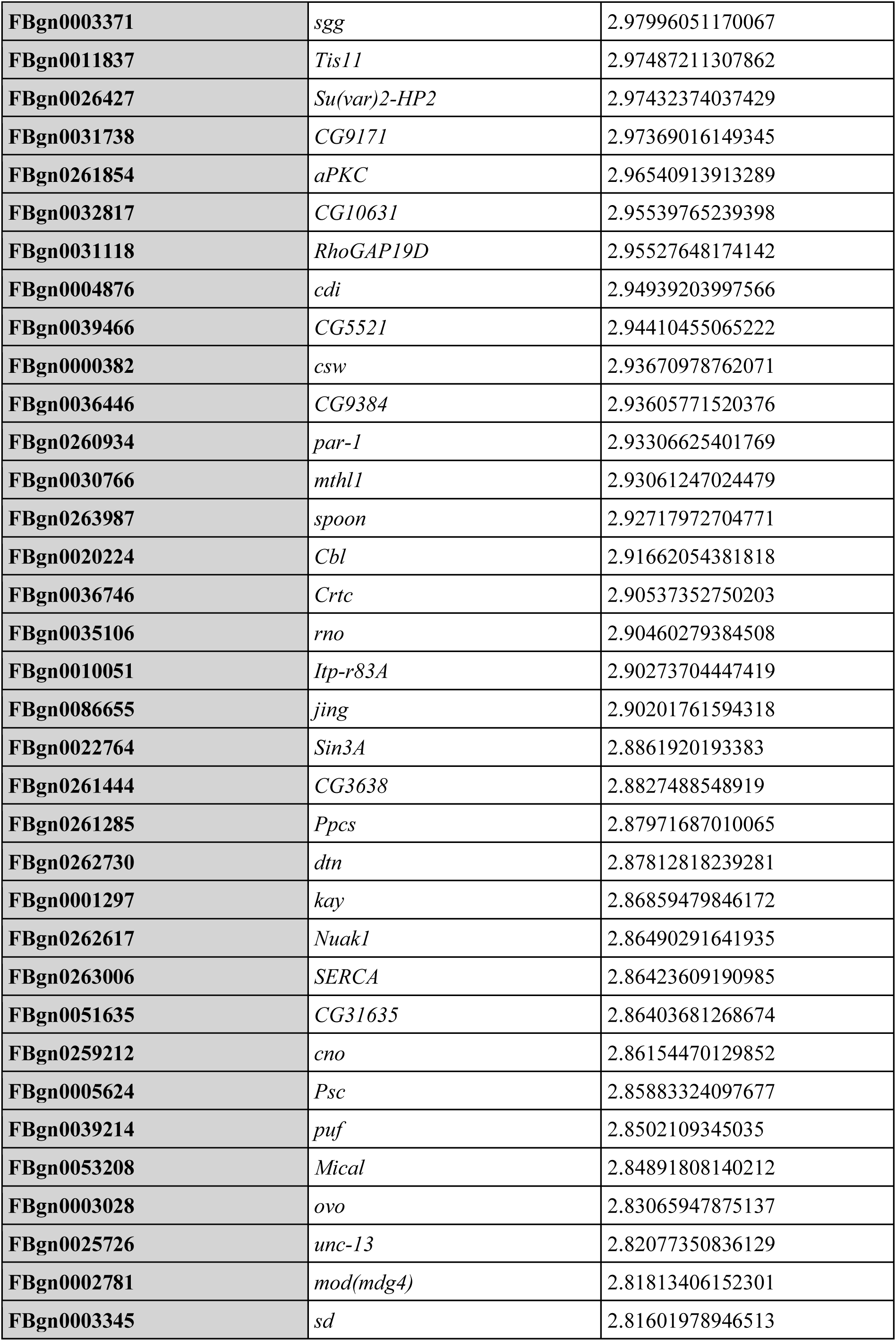
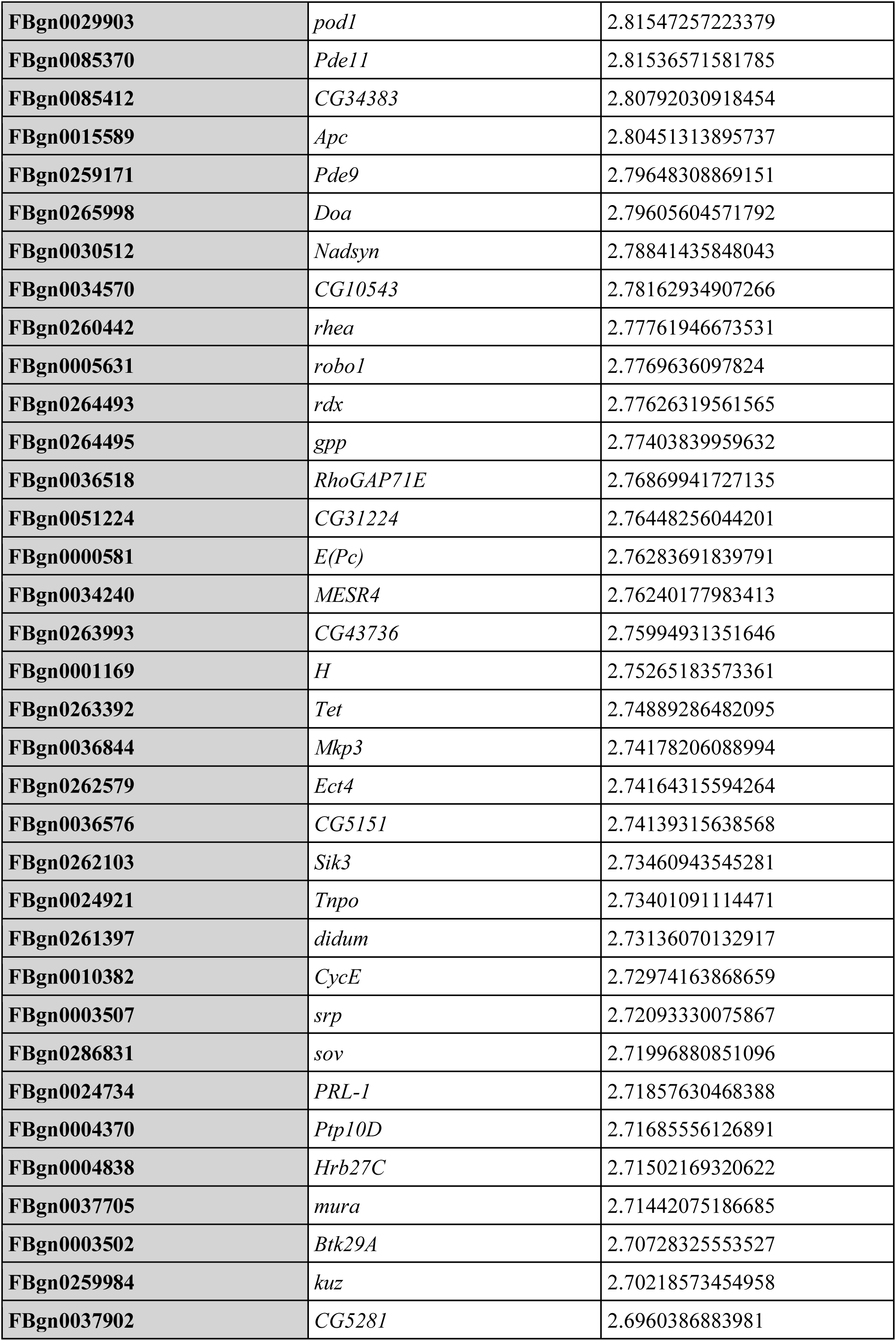
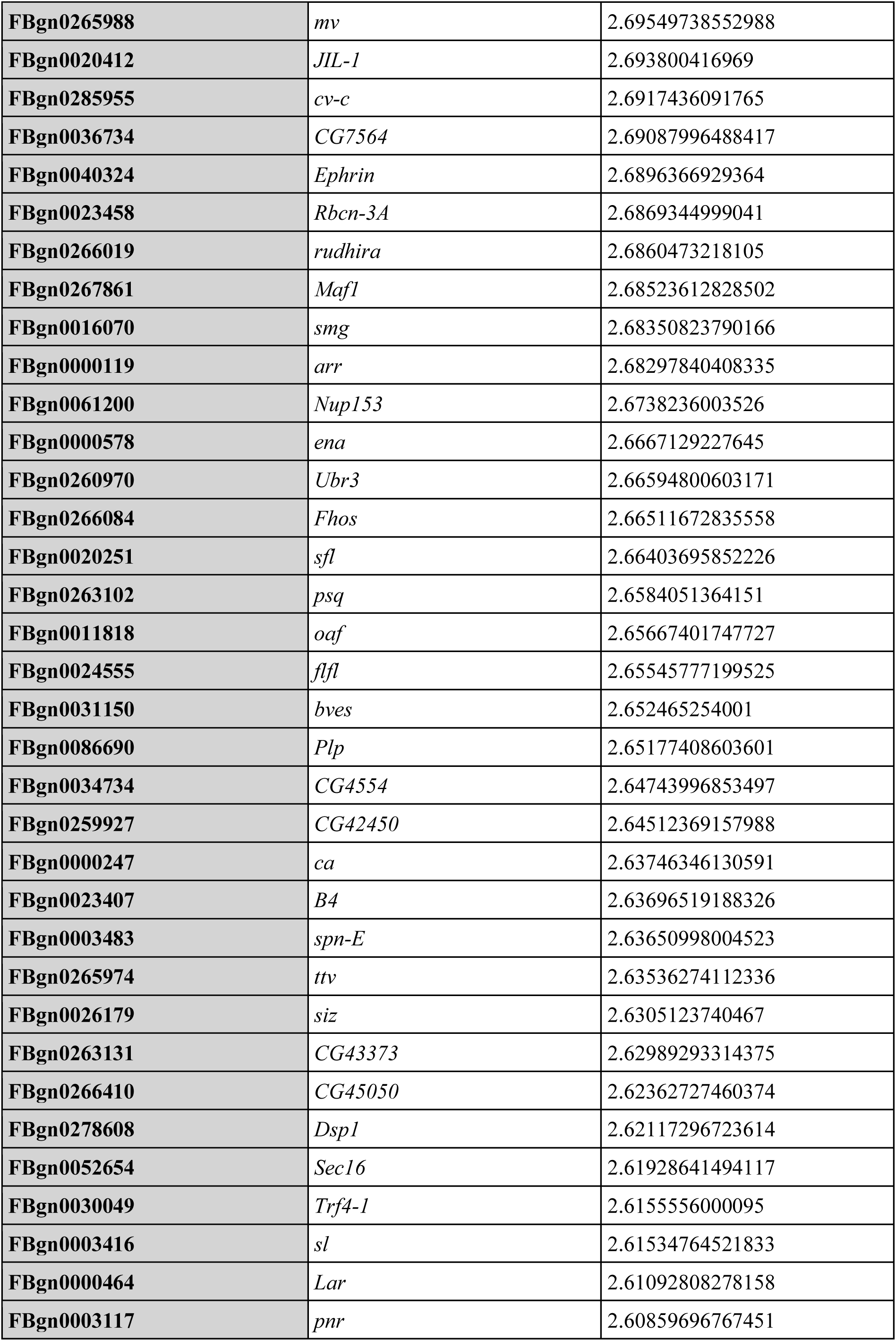
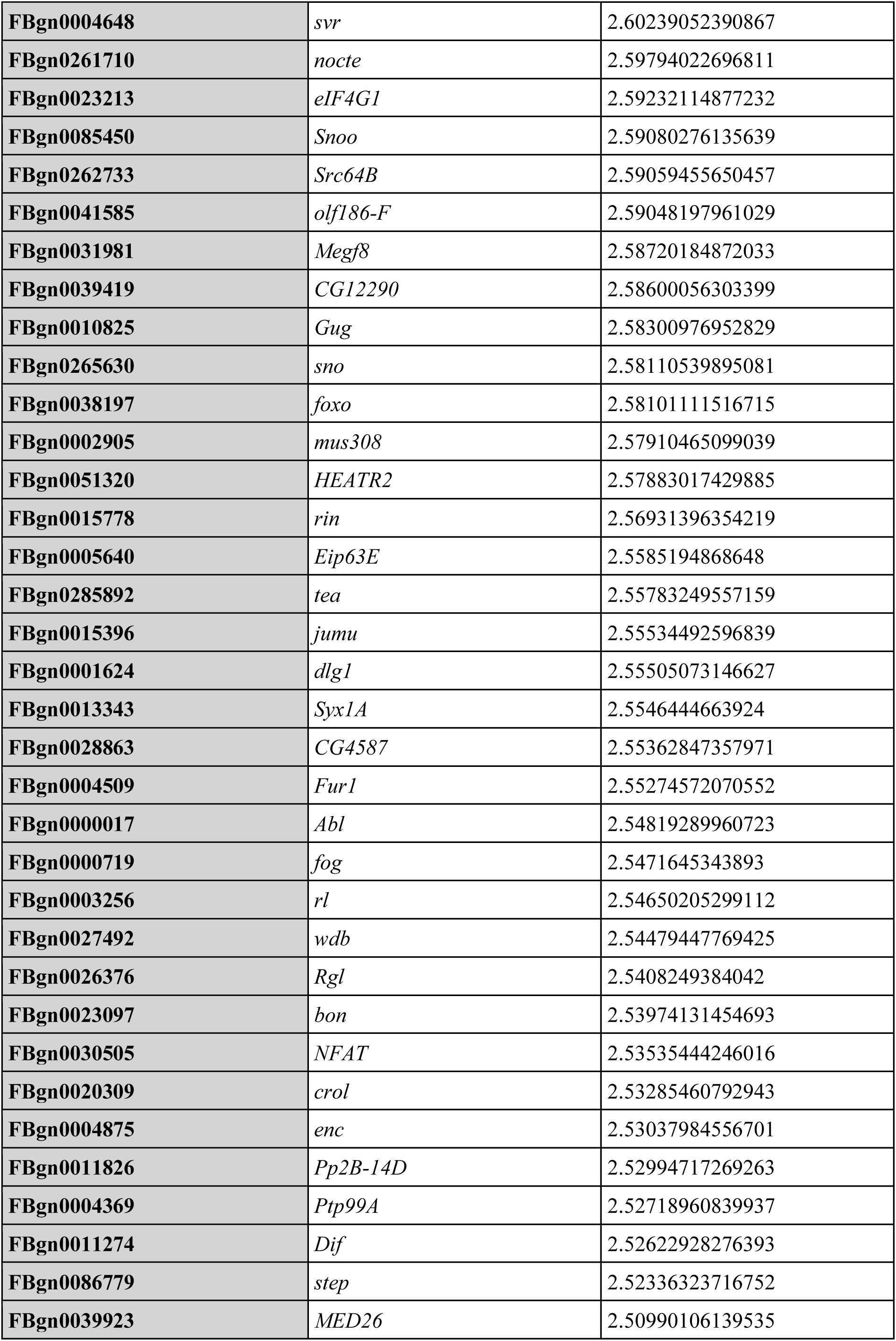
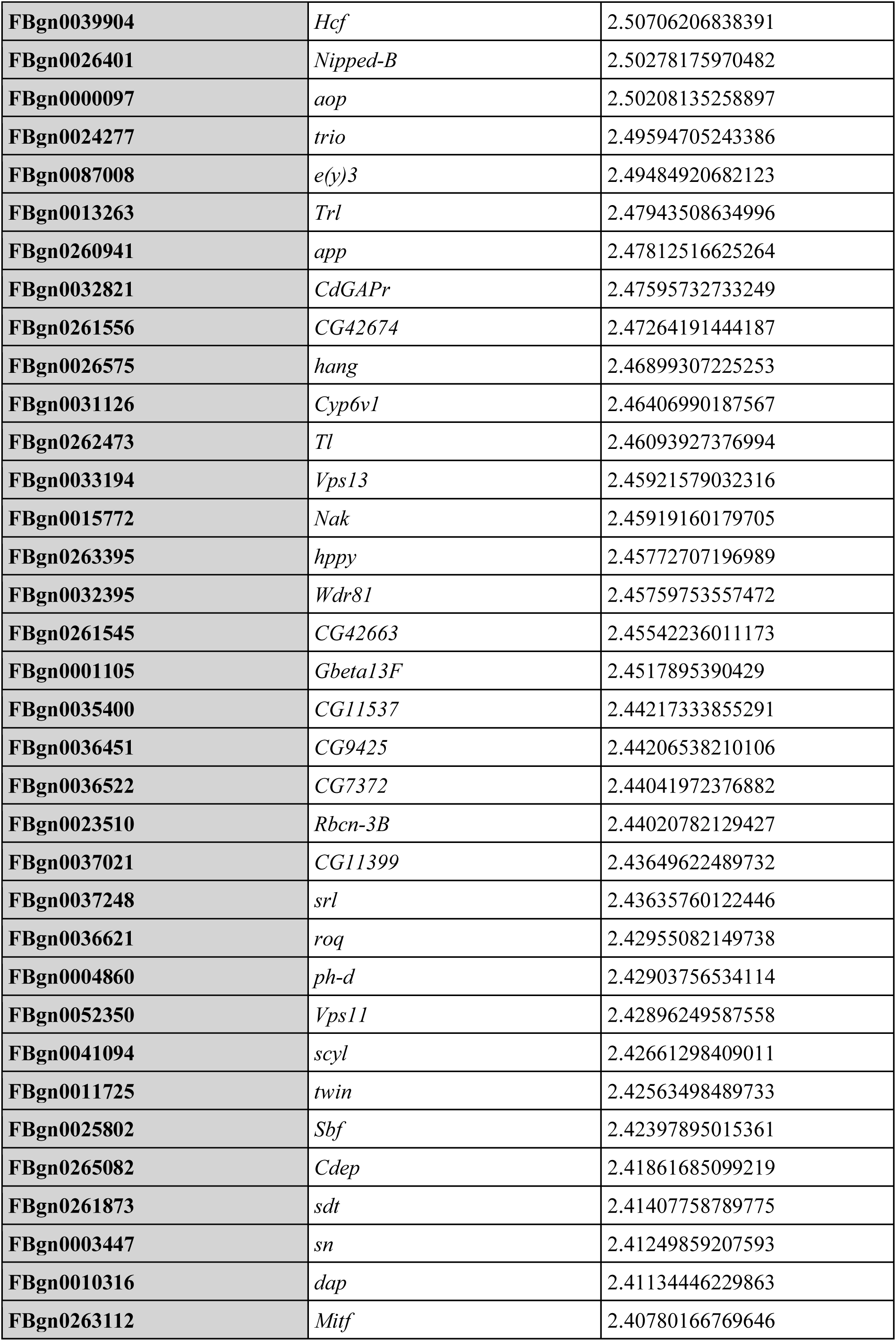
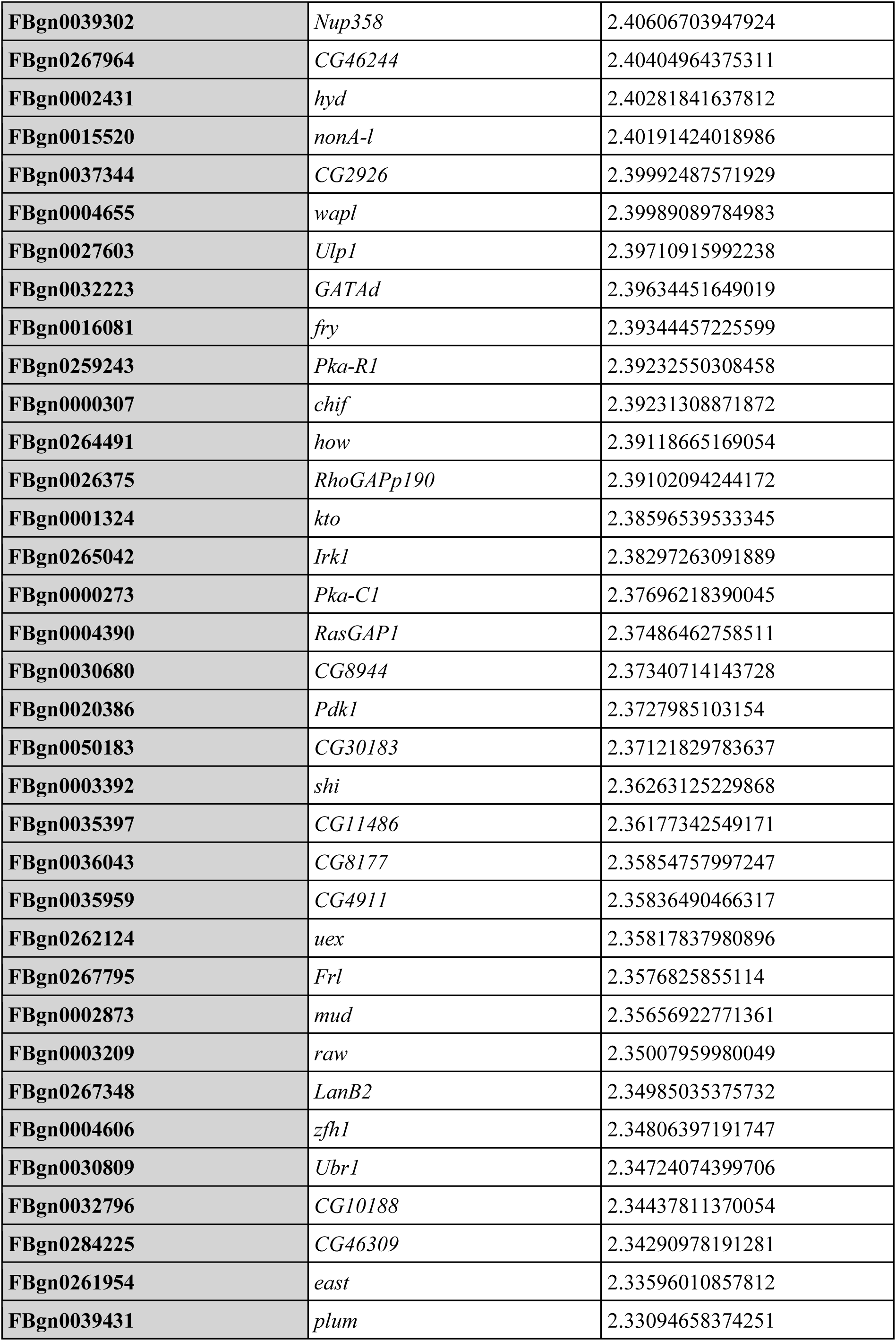
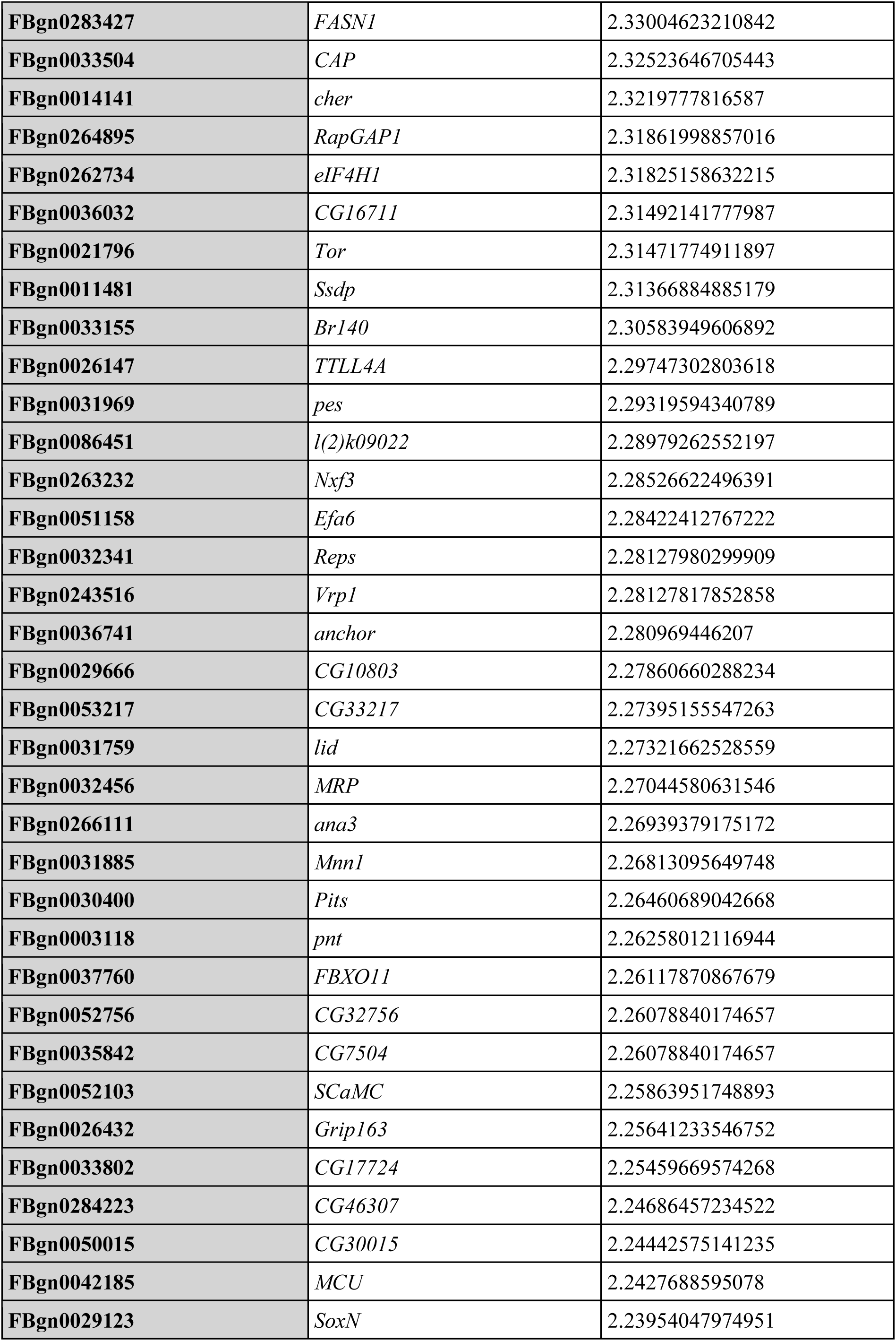
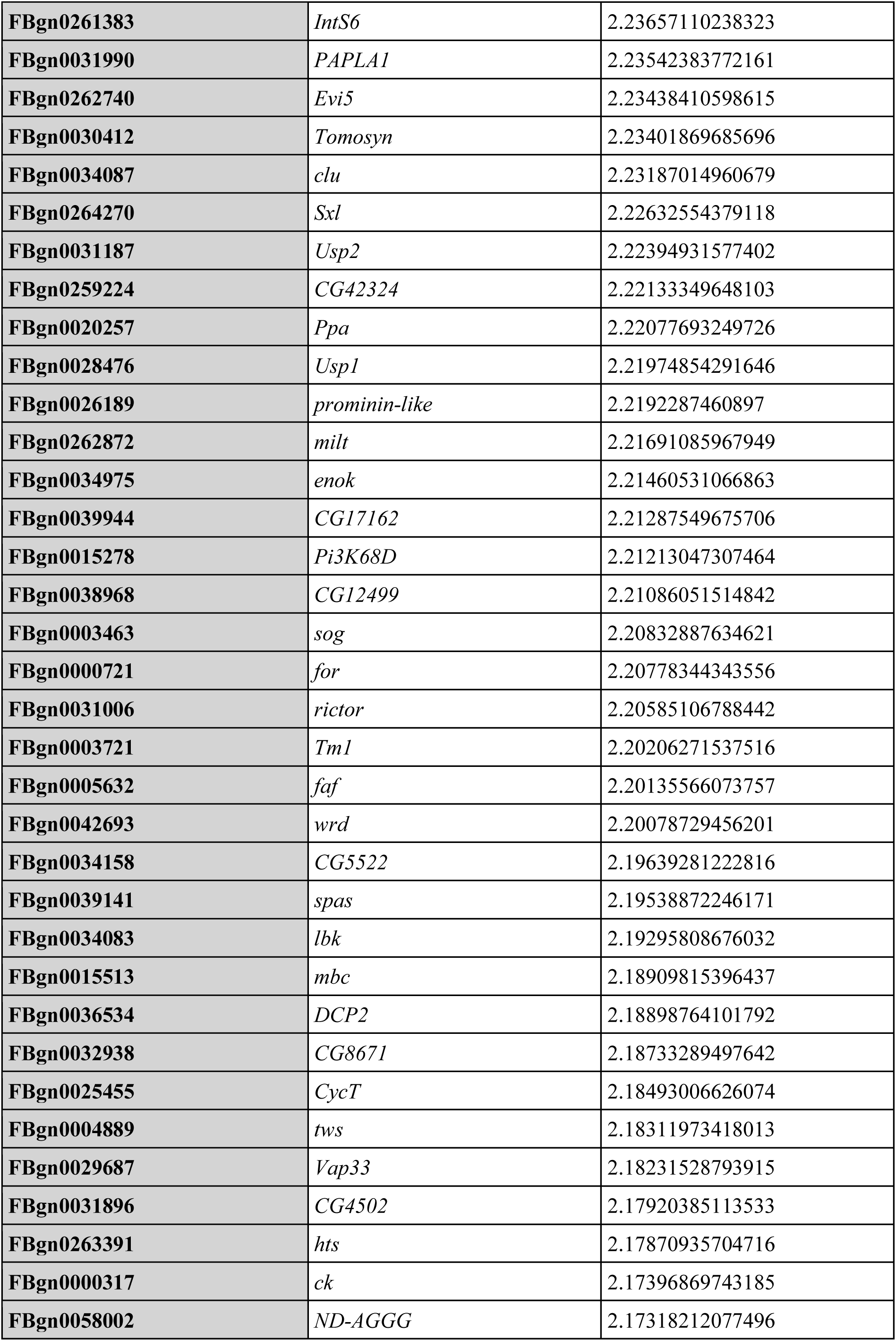
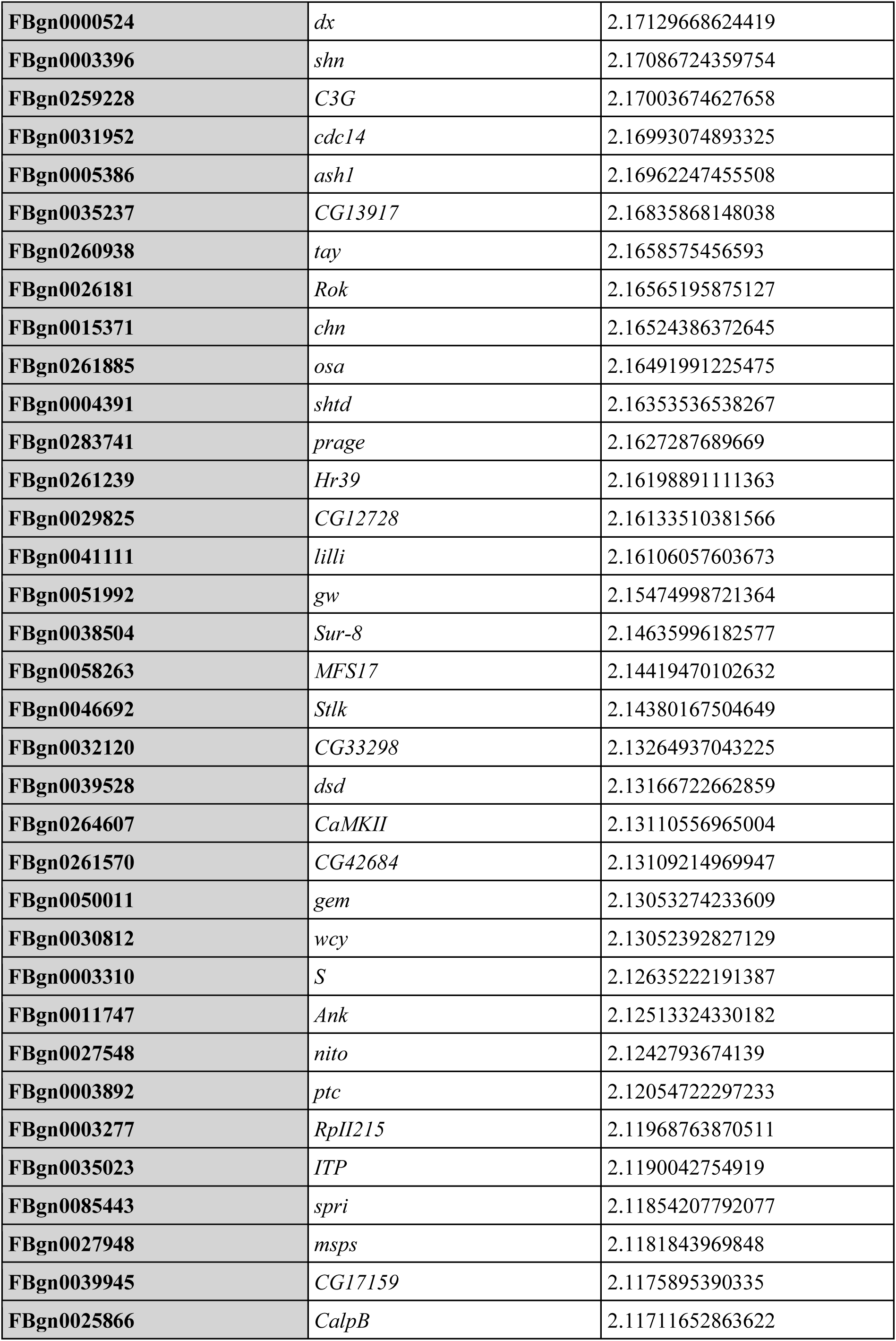
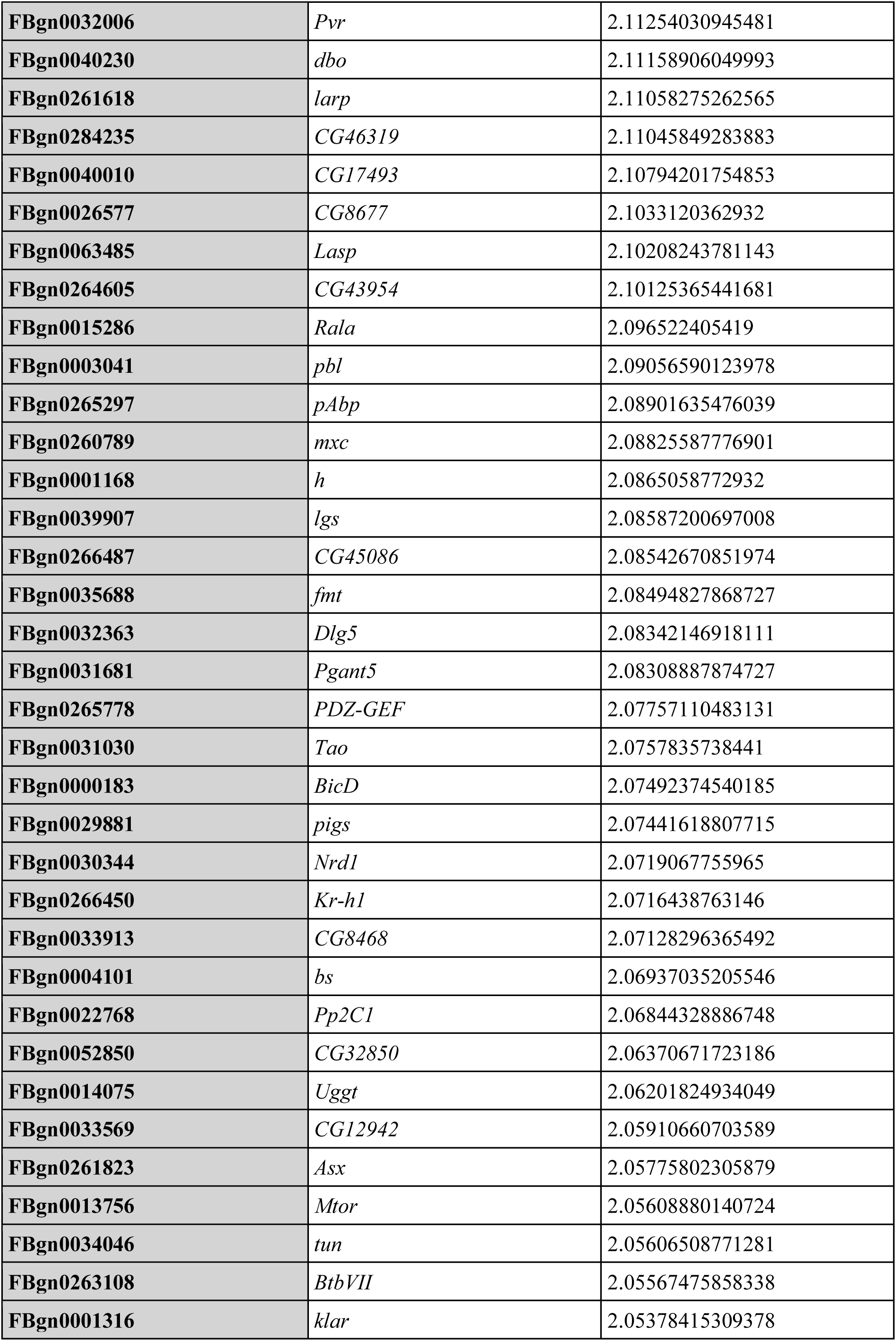
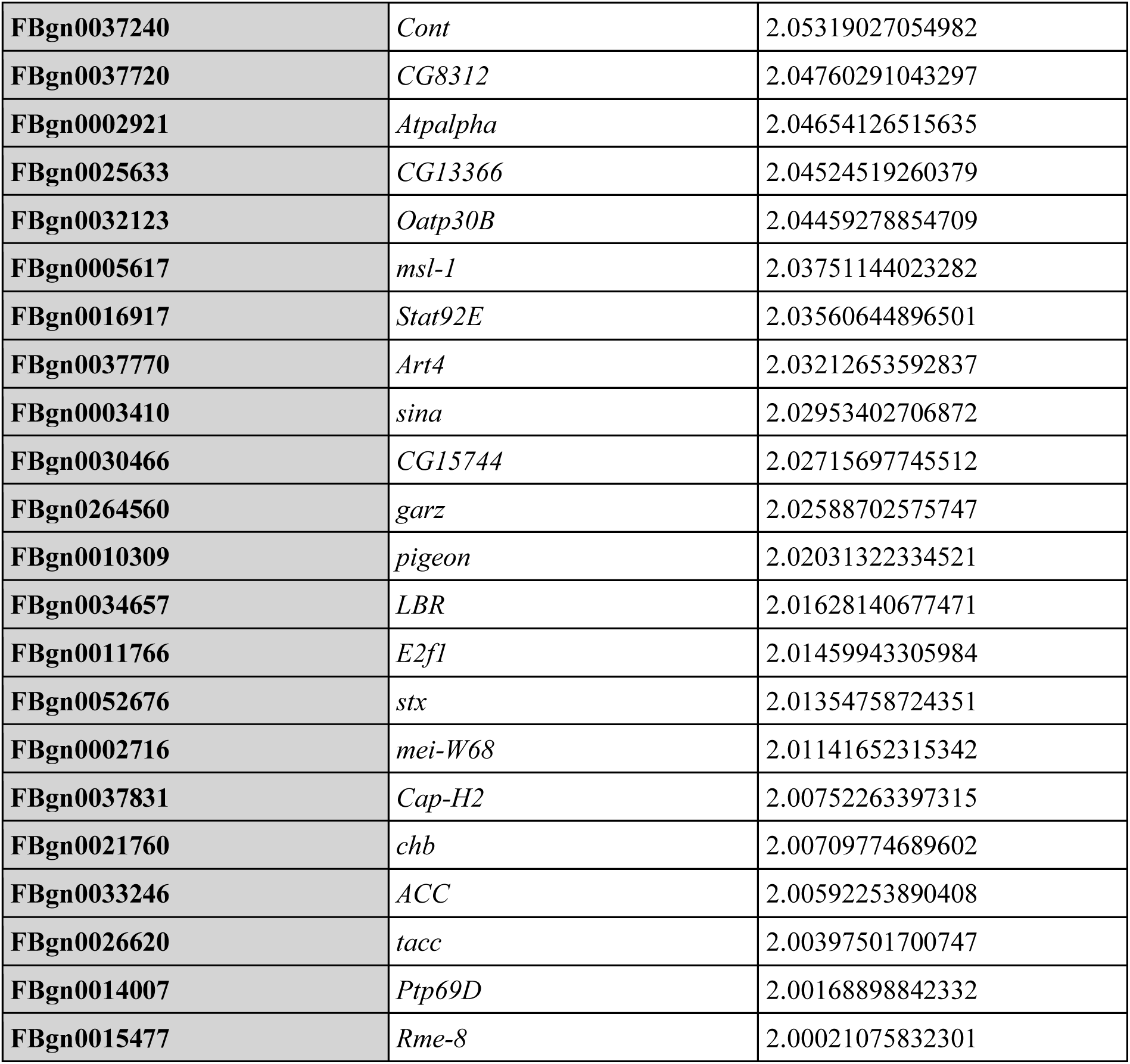
dMi-2-GFP iCLIP-seq list of enriched mRNAs with Fold change > 4 and (*p < 0.05*)

We performed RNA immunoprecipitation (fRIP) to corroborate our iCLIP2 data. After formaldehyde crosslinking, we used GFP-Trap beads to immunoprecipitate extracts of cells expressing dMi-2-GFP, dLint1-GFP and of cells not expressing GFP-tagged protein (control) **(Supplementary Figure 1D)**. Co-precipitated RNAs were then analysed by RT-qPCR. Like dMi-2, dLint1 is a subunit of a chromatin associated repressor complex (LINT complex, (Macinkovic et al., 2019; Meier et al., 2012)). We analysed nine dMi-2-associated transcripts, seven that were preferentially crosslinked to dMi-2 and two that were not (highlighted in **Figure 1C**). None of these transcripts were significantly enriched in dLint1 precipitates compared to control precipitates **(Supplementary Figure 1E)**. In contrast, significant enrichment in dMi-2 precipitates over Lint1 and control precipitates was detected for 7 transcripts. This result corroborates the iCLIP data and strengthens the hypothesis that dMi-2 associates with RNAs *in vivo*.

### RNA binding is unlikely to act in the recruitment of dMi-2 to chromatin

Association of dMi-2/CHD4 with nascent RNA has previously been suggested to assist in the recruitment of dMi2/CHD4/NuRD to chromatin at heat shock and rRNA genes (Mathieu et al., 2012; Zhao et al., 2018). A widespread role for RNA binding in mediating the chromatin association of dMi-2 would predict that dMi-2 ChIP-seq and dMi-2 iCLIP2 patterns across genes are correlated. A metagene analysis of dMi-2 RNA crosslink sites revealed preferential crosslinking towards the 3’ end of transcripts (**Figure 2A**; **Supplementary Figure 2A)**. However, a metagene analysis of dMi-2 chromatin binding sites at these genes as determined by ChIP-seq (Lenz et al., 2021) highlights a clear preference for promoter sequences and the 5’ region of genes **(Figure 2A)**. In fact, dMi-2 chromatin occupancy is lowest towards the 3’ end of genes. Moreover, dMi-2 target genes identified by iCLIP and ChIP-seq, respectively, show only a limited overlap **(Figure 2B)**. Closer inspection of iCLIP crosslink site distribution over the intron/exon junction preceding the transcription termination site (TES; also referred to as the cleavage and polyadenylation site) reveals that crosslinking of dMi-2 to the terminal exon is dramatically more likely than crosslinking to the terminal intron. While this observation does not rule out the possibility that dMi-2 also binds nascent RNA, it demonstrates that dMi-2 is predominantly associated with processed RNA **(Figure 2C)**. Taken together, these results indicate that the binding of dMi-2 to nascent RNA is unlikely to play a dominant role in recruiting dMi-2 to chromatin.

**Figure 2:**
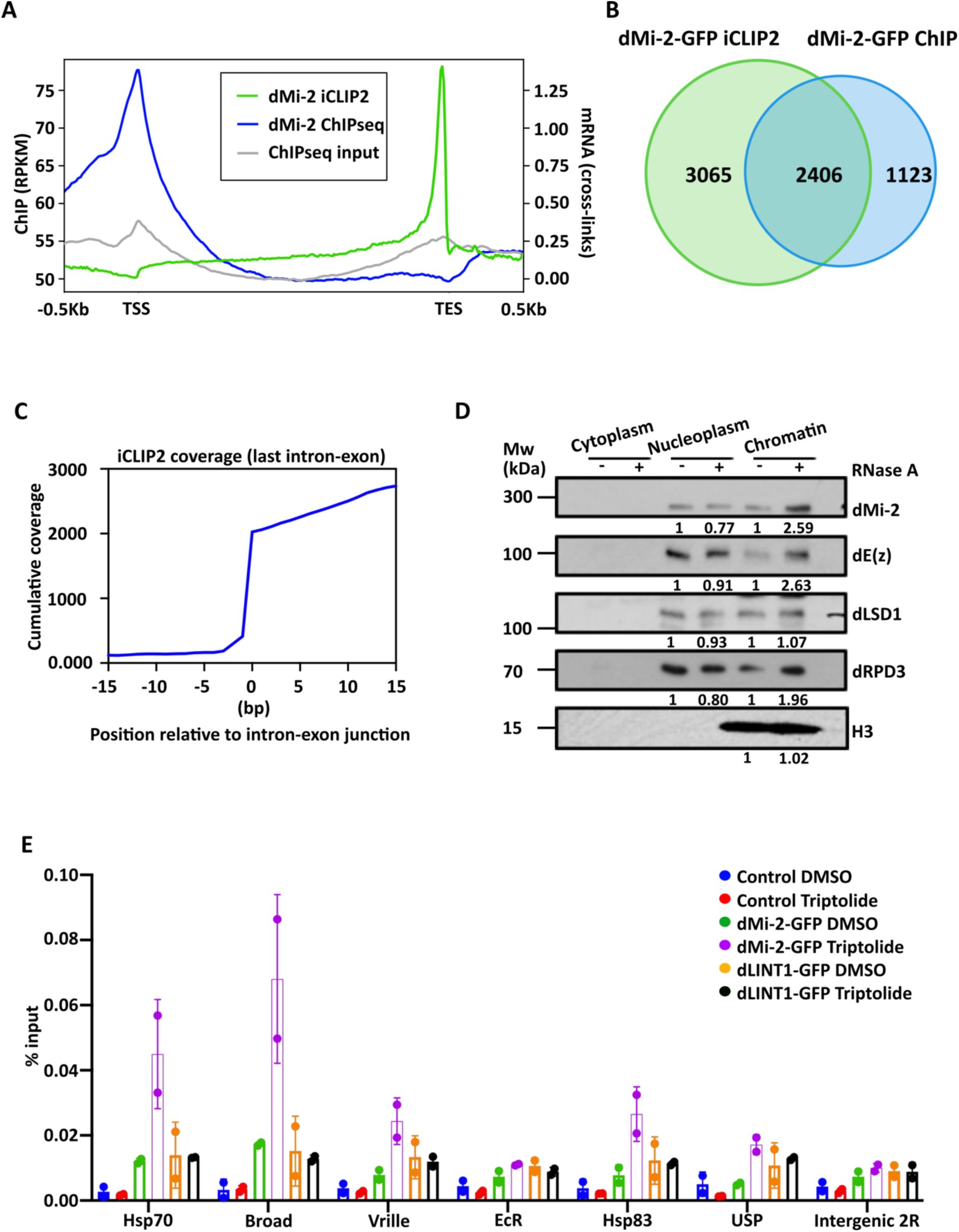
RNA binding does not play a major role in recruiting dMi-2 to chromatin. **(A)** Distribution of dMi-2 iCLIP2 crosslinking sites (green), dMi-2 ChIPseq (blue) and ChIPseq input (grey) across a metagene. The transcription start site (TSS) and transcription end site (TES) are indicated. Left Y-axis shows ChIP-seq signal strength (RPKM), right Y-axis shows iCLIP2 signal strength (number of crosslinks). **(B)** Venn diagram showing the overlap between dMi-2 bound RNAs and genes as determined by iCLIP2 and ChIP-seq, respectively. **(C)** Accumulated iCLIP2 read coverage across the last intron-exon junction. All annotated genes with at least two exons were aligned at the last junction between intron and exon. The first base of the exon is at position zero. **(D)** S2 Cells were permeabilized and treated with RNase A (+) or BSA (-). Cells were fractionated into cytoplasmic, nuclear and chromatin and analysed by Western Blot as indicated. Histone H3 was used as loading control. The western blot signals were quantified using ImageJ software. The BSA treated control signals were set to 1 and the fold change in the RNase A treated samples was calculated and is shown below each panel. **(E)** Control (blue and red), dMi-2-GFP (green and purple), and dLINT1-GFP (orange and black) expressing S2 cells were treated with DMSO or 10 µM Triptolide, respectively, for 3 hours, as indicated on the right. Chromatin was prepared and subjected to ChIP-qPCR analysis. Individual data points of two independent biological replicates were plotted as percent of input. Error bars indicate the standard deviation between two biological replicates.

### RNA antagonizes the interaction of dMi-2 with chromatin

NuRD and PRC2 cooperate to establish repressed chromatin regions characterized by high nucleosome occupancy and nucleosomes containing H3K27me3 (Bracken et al., 2019; Reynolds et al., 2012a). RNA has recently been found to contribute to bringing PRC2 to its genomic targets but also to dissociate the complex from chromatin at actively transcribed genes (Beltran et al., 2019; Beltran et al., 2016; Long et al., 2020). Given the close cooperation between PRC2 and NuRD, we considered the possibility that RNA simultaneously removes both repressive complexes from chromatin. To test this hypothesis, we treated permeabilized S2 cells with RNase A or BSA (control). We then biochemically fractionated cells into cytoplasm, nucleoplasm and chromatin, and subjected these fractions to Western blot analysis **(Figure 2D)**. We found that RNase treatment results in a decrease of the nucleoplasmic fraction of the PRC2 subunit Enhancer of zeste (E(z)) and a corresponding increase of the chromatin-bound fraction of the same protein. This observation agrees with published data that demonstrate a similar behaviour of EZH2 from mouse ES cells in this assay, suggesting that RNA-mediated chromatin displacement of PRC2 is conserved between mammals and insects (Beltran et al., 2016). The chromatin-associated fractions of dMi-2 and the dNuRD subunit dRPD3 were also increased upon RNA digestion. In contrast, the distributions of dLSD1, a histone demethylase, and of histone H3 were not significantly affected by RNase treatment. Thus, RNA-dependent displacement from chromatin is not a general property of chromatin-associated proteins in this assay.

If RNA antagonizes chromatin association of dMi-2 *in vivo*, inhibition of transcription would be expected to facilitate dMi-2 binding to chromatin. We treated S2 cells with triptolide or DMSO (control) to test this hypothesis. Triptolide induces degradation of the largest subunit of RNA polymerase II. As a consequence, transcription at active genes is strongly reduced (Beltran et al., 2016; Riising et al., 2014) **(Supplementary Figure 2B)**. We used ChIP-qPCR to determine the effect of inhibition of transcription on dMi-2 chromatin binding at six established dMi-2 target genes (Kreher et al., 2017) **(Figure 2E)**. Five of these showed increased dMi-2 chromatin binding in triptolide-treated cells. No such increase of dMi-2 chromatin binding was detected at an untranscribed control region (intergenic 2R). Importantly, dLint1, although bound to the same genes, displayed no increased chromatin association when transcription was inhibited. This demonstrates that increased chromatin binding upon triptolide treatment is specific for dMi-2.

Taken together, these results establish that RNA counteracts dMi-2 chromatin association *in vivo*.

### dMi-2 has broad RNA binding activity in vitro

Because dMi-2 exists in two multi-subunit complexes: dNuRD and dMec (Kunert et al., 2009), dMi-2 RNA binding and the effects of RNA on chromatin association could be mediated by other subunits of these complexes. We sought to investigate the inherent RNA binding activity of dMi-2 *in vitro*. We used electrophoretic mobility shift assays (EMSA) to investigate RNA binding by purified, recombinant dMi-2 **(Supplementary Figure 3A; Table 2)**. We tested dMi-2 binding to segments of two RNA molecules: the Hsp70Aa transcript, which we have previously shown to bind dMi-2 during the heat shock response (Murawska et al., 2011), and the unrelated Pgant35a transcript. The two transcripts are similar in length (Hsp70Aa: 2.4 kb, Pgant35a: 2.2 kb). For each RNA we selected four 100 nucleotide regions spanning the transcript **(Figure 3A**, regions #1, #2, #4 and #5; **Supplementary Figure 3B)**. All eight RNA molecules were efficiently bound by dMi-2 **(Figure 3B)**. In contrast to the apparent preference of dMi-2 for binding towards the 3’ ends of transcripts *in vivo* **(Figure 2A)**, we did not detect strong differences in binding to 5’ and 3’ regions by recombinant dMi-2 *in vitro*. These results indicate that dMi-2 has broad RNA binding activity *in vitro*.

**Figure 3:**
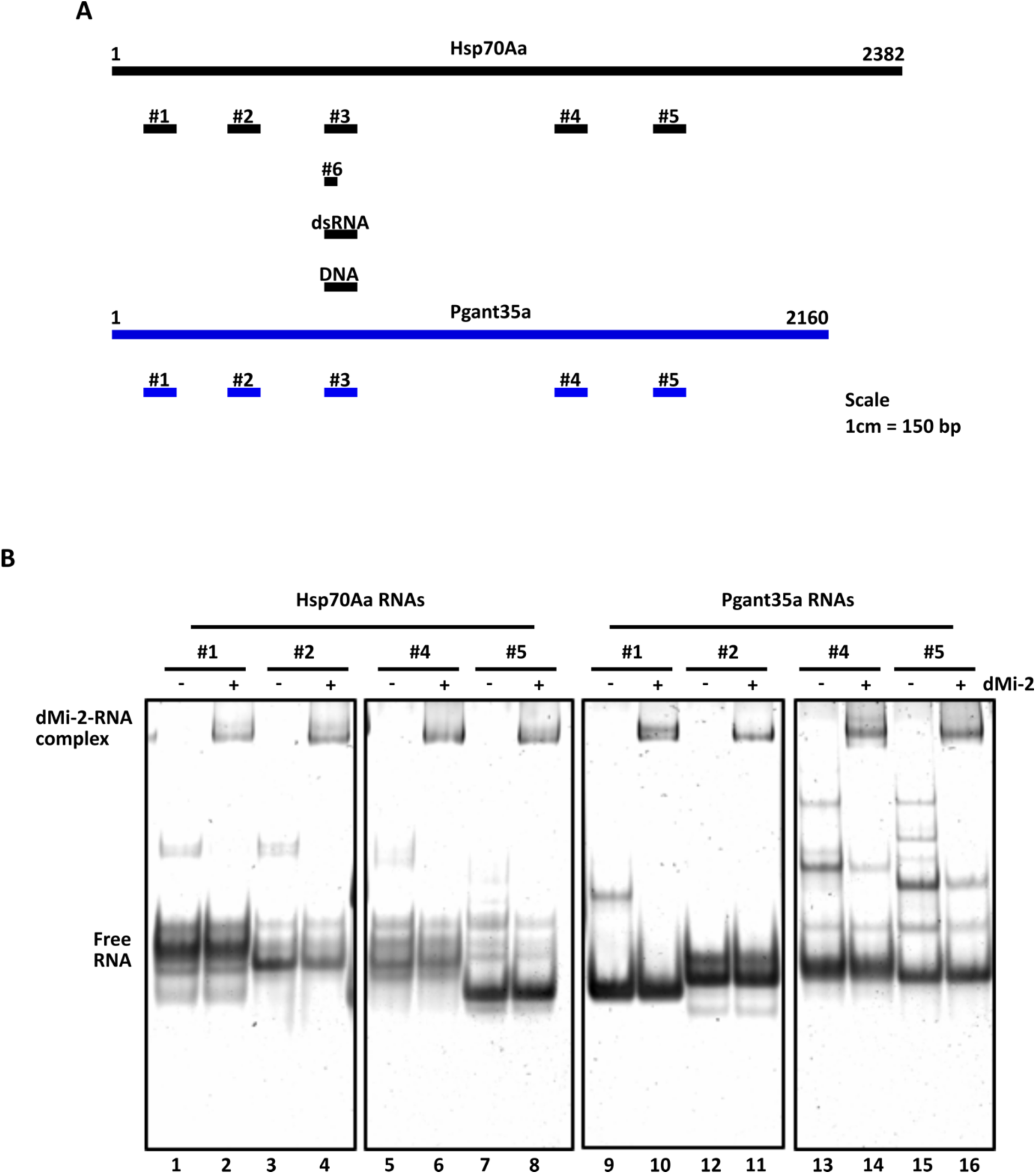
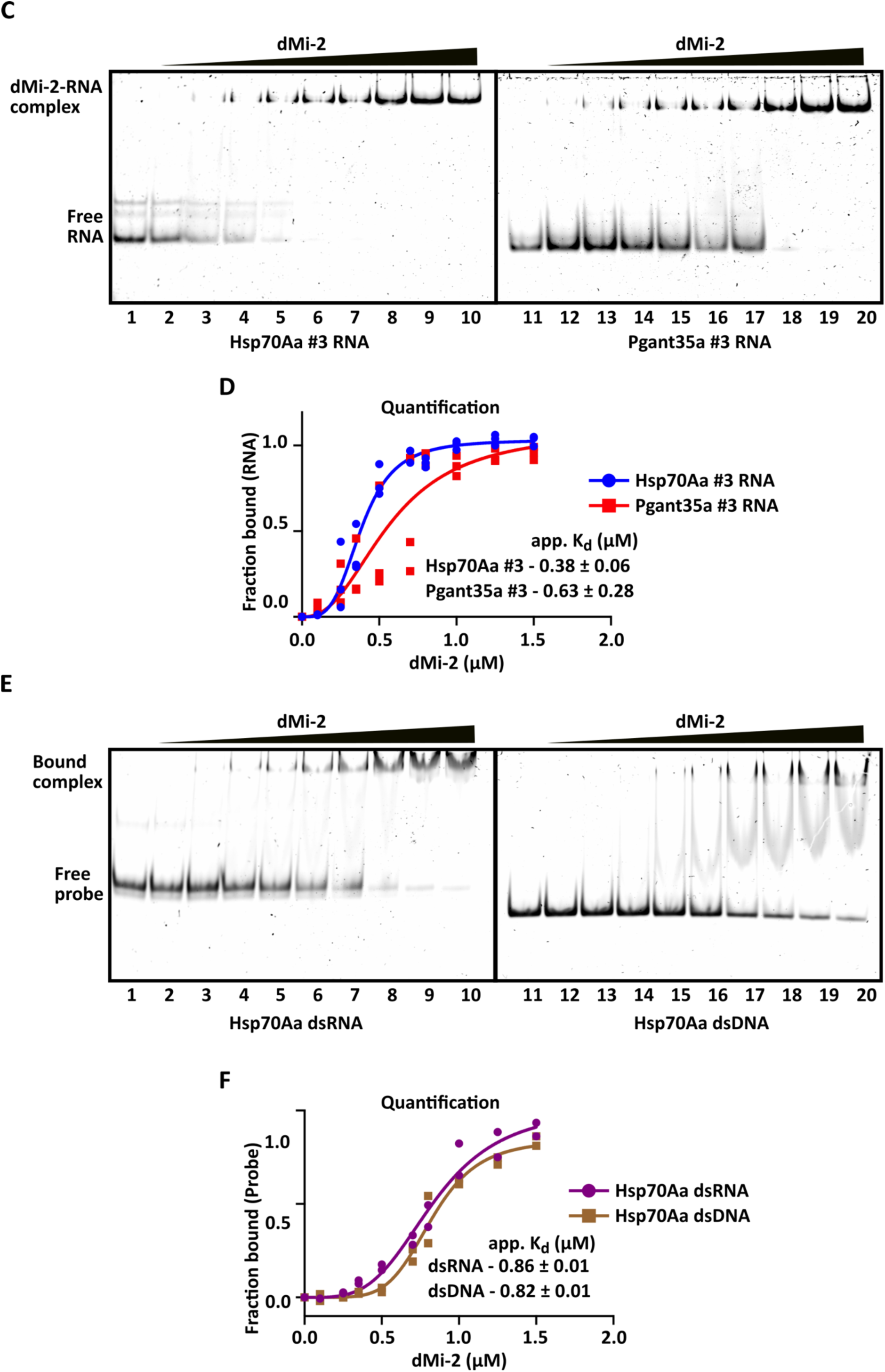
dMi-2 has broad RNA binding activity in vitro. **(A)** Schematics representing the regions of Hsp70Aa (black) and Pgant35a (blue) genes from which corresponding RNAs were generated. The RNAs #1, #2, #3, #4 and #5 are all 100 nucleotides in length. RNA #6 is 40 nucleotides in length. The location of 100 bp double stranded RNA (dsRNA) and DNA fragments from the Hsp70Aa gene, used in Figure 3E are indicated (dsRNA, DNA). Scale is indicated on the right. **(B)** dMi-2 binding to RNA fragments derived from Hsp70Aa and Pgant35a genes. For EMSAs, 220 nM of each RNA and 112 nM of recombinantly purified dMi-2 was used per reaction. RNA without protein was loaded in lanes 1, 3, 5, 7, 9, 11, 13 and 15. Gels were stained with SYBR Gold. **(C)** EMSAs with Hsp70Aa #3 RNA (lanes 1-10) and Pgant35a #3 RNA (lanes 11-20). 250 nM RNA was used in each reaction and recombinantly purified dMi-2 was titrated from 0.1 to 1.5 *μ*M. RNA without protein was loaded in lanes 1 and 11, respectively. Representative image of three independent biological replicates is shown. **(D**) Quantification of the EMSA shown in (C). The fraction of the RNA bound by dMi-2 is plotted against the concentration of dMi-2. Data points from three independent biological replicates are shown. The apparent dissociation constant (app. K_d_) indicated in (D) and (F), were derived with Hill slope equation using GraphPad Prism software. **(E)** EMSA of corresponding 100 bp double stranded RNA (dsRNA) and double stranded DNA (dsDNA) fragments derived from Hsp70Aa gene. 250 nM dsRNA or dsDNA was used for each reaction and dMi-2 was titrated from 0.1 to 1.5 μM. dsRNA and dsDNA controls without protein were loaded in lanes 1 and 11, respectively. Representative image of two independent biological replicates. **(F)** Quantification of the EMSA shown in (E). Data points from two independent biological replicates are shown. The gels were stained with SYBR Gold.

**Table 2:**
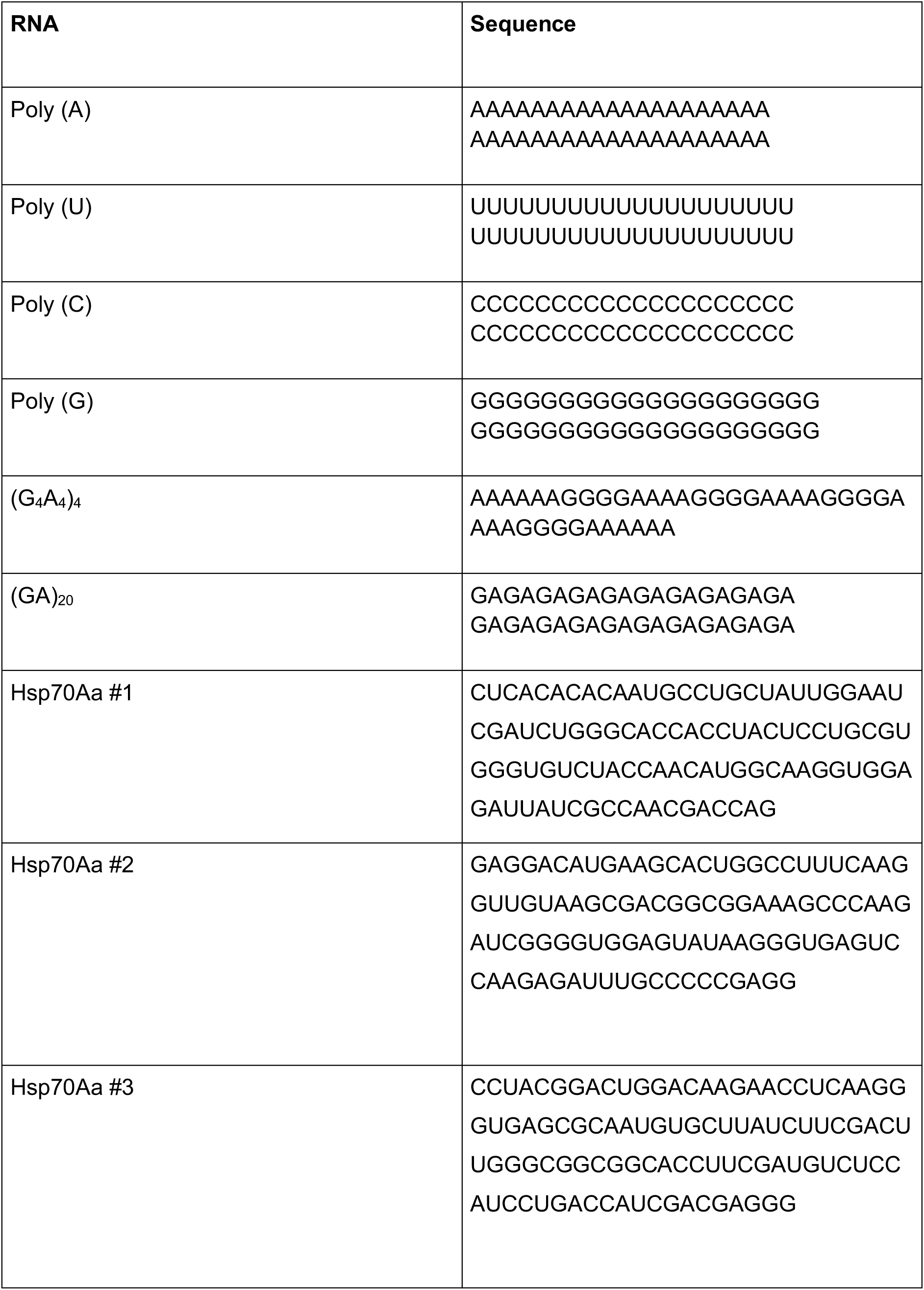
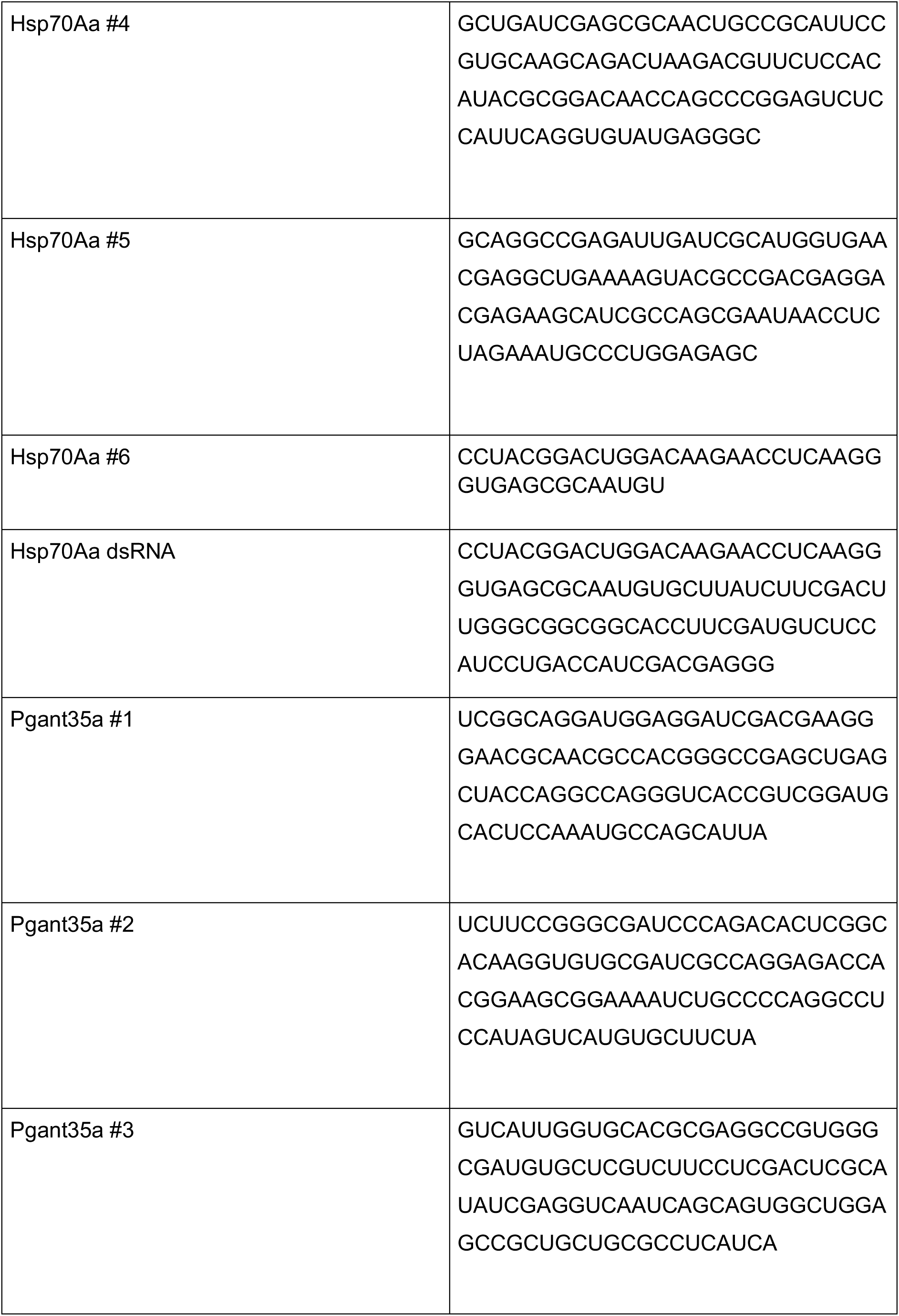
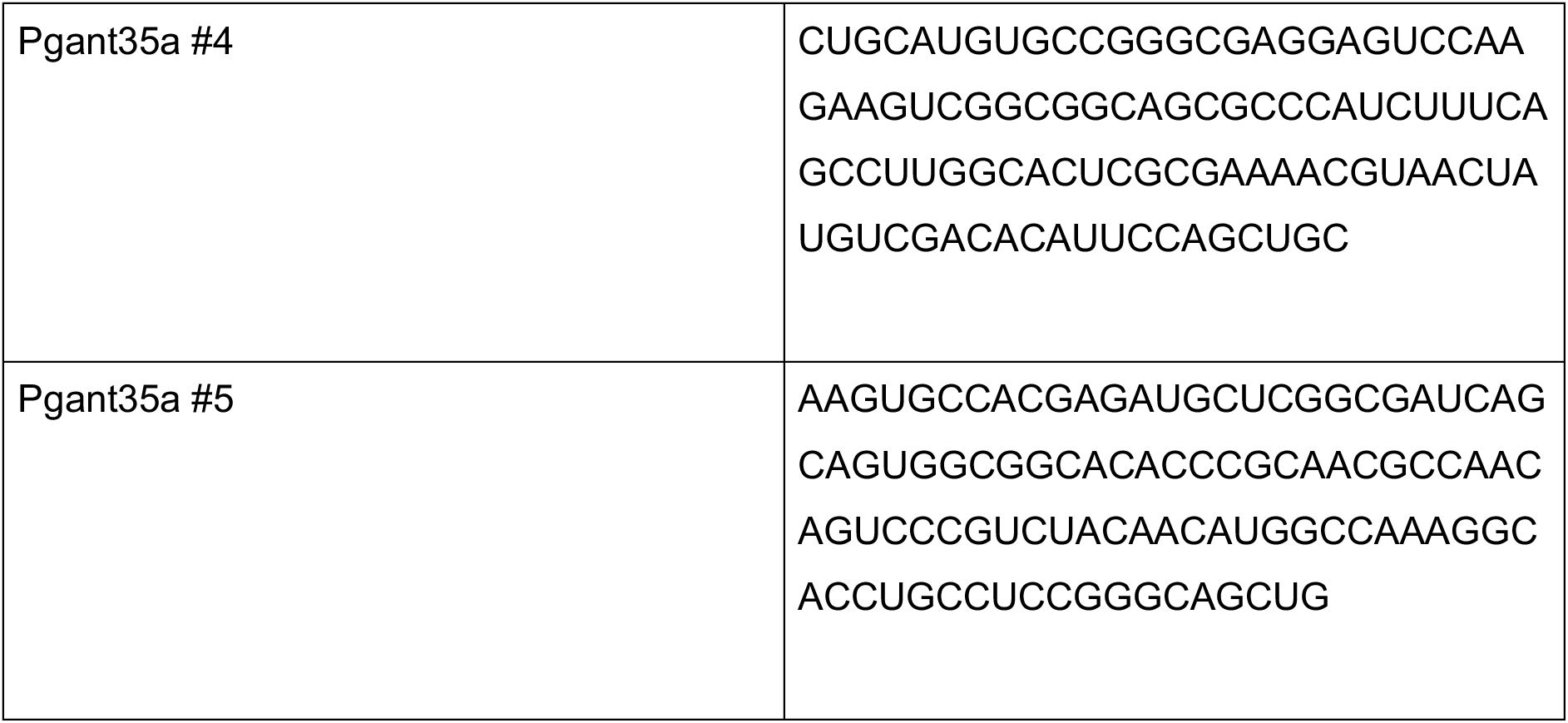
RNAs used in EMSAs.

We next asked if we could detect quantitative differences of dMi-2 binding to different RNA molecules. We performed titration experiments to derive dissociation constants (K_d_) for two 100 nucleotide RNA fragments (Hsp70Aa #3 and Pgant35a #3, **Figure 3A, Supplementary Figure 3C)**. The apparent K_d_ values for dMi-2 binding to Hsp70Aa and Pgant35a RNAs were 0.38 (+/- 0.06) μM and 0.63 (+/- 0.28) μM, respectively **(Figure 3C and 3D)**. This result shows that the dMi-2 binding affinities of RNA molecules with identical length but nonidentical nucleotide sequence (and potentially different secondary structures) are overall quite similar, but can exhibit subtle differences.

We also compared the affinities of dMi-2 binding to double stranded RNAs and to double stranded DNAs of corresponding length and sequence **(Figure 3E and F, Supplementary Figure 3D and E)**. dMi-2 bound to a 100 bp double stranded Hsp70Aa fragment with an apparent K_d_ of 0.86 (+/- 0.1) μM. The corresponding double stranded DNA fragment of the same sequence was bound with a similar apparent K_d_ of 0.82 (+/- 0.1) μM. These K_d_ values are higher by a factor of two than the apparent K_d_ value for binding of dMi-2 to the corresponding single stranded RNA molecule **(Figure 3D)**. This agrees with previous findings and indicates that while dMi-2 can bind to double stranded RNA and DNA it binds with higher affinity to single stranded RNA (Murawska et al., 2011). Furthermore, dMi-2 complexes with single stranded RNAs were remarkably resistant to high salt (800 mM KCl) and high detergent concentrations (8% NP40) **(Supplementary Figure 3F and 3G)**, supporting the conclusion that dMi-2 has robust RNA binding activity in the absence of dMi-2-associated dNuRD or dMec subunits.

### dMi-2 preferentially binds G-rich RNAs

Our analysis revealed that dMi-2 has broad RNA binding activity *in vitro*, in agreement with our finding that dMi-2 is associated with many RNAs *in vivo* **(Figure 1)**. The results also indicate that dMi-2 does not bind with the same affinity to all RNA molecules, suggesting that sequence (and/or secondary structure) might modulate binding. We asked if RNA regions to which dMi-2 was crosslinked *in vivo* had a non-random sequence composition surrounding the crosslinking site that would facilitate dMi-2 binding. The relative distribution of bases surrounding dMi-2 crosslinking sites is shown in **Figure 4A**. Of note, as a consequence of UV-induced crosslinking, U is more likely to be crosslinked to proteins than other bases (Sugimoto et al., 2012). Indeed, U was identified at the crosslinking site much more often than the remaining three bases. Of these, G was the most abundant at both the crosslinking site (position 0) as well as at the neighbouring position +1. Furthermore, this analysis detected an overrepresentation of A and C at positions +4 and +5, respectively, indicating that dMi-2 preferentially associates with a GGNNAC motif *in vivo*. To further probe the effect of base composition on dMi-2 binding we used homopolymeric RNAs in EMSAs. We compared binding of dMi-2 to poly(G), poly(C), poly(U) and poly(A) 40-mers **(Figure 4B, Supplementary Figure 4A to D)**. dMi-2 bound to poly(G) RNA with an apparent K_d_ of 0.03 (+/- 0.01) μM. The affinity for binding to the other three homopolymeric RNA fragments was too low to allow us to determine an apparent K_d_ value. These results indicate that dMi-2 interacts preferentially with G-rich RNA.

**Figure 4:**
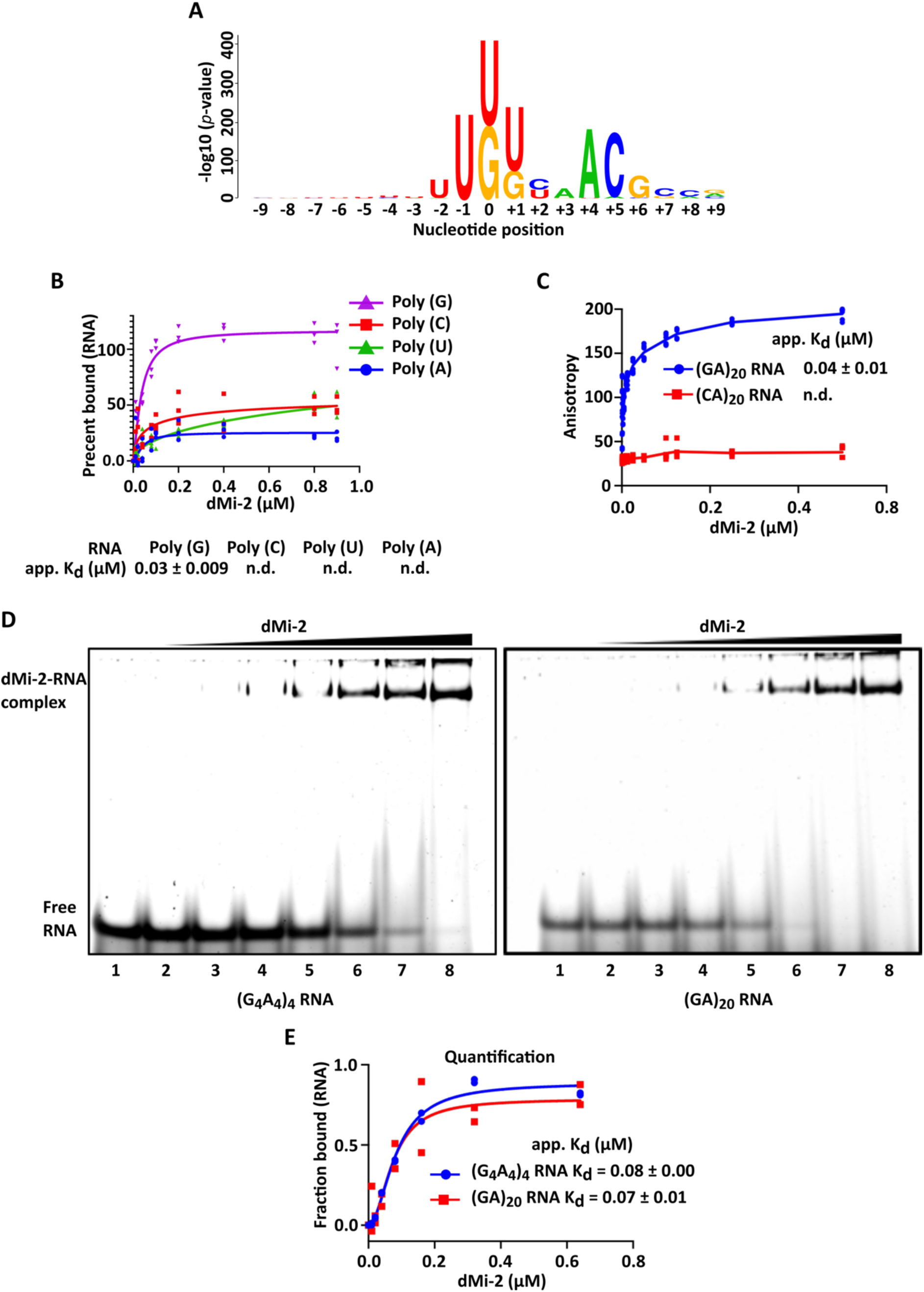
dMi-2 preferentially binds G-rich RNAs. **(A)** KpLogo sequence plot dicating enrichment of G-nucleotides at the crosslinking site and possible preferential binding to 5’- GGNNAC-3’ sequence. Nine nucleotides up- and down- stream (-9, +9) of the crosslinking site (0) are plotted. The size of nucleotide letters represents the enrichment as (p-value) above expected frequency sampled from random mRNA sequences. Crosslinking to uridine nucleotides is favoured due to use of UV-C irradiation. Therefore, the higher frequency of U-nucleotides is disregarded. The second most frequently crosslinked nucleotide is guanine. **(B)** Quantification of EMSAs of homopolymer RNAs Poly (A) (blue), Poly (U) (green), Poly (C) (red) and Poly (G) (purple). Each RNA is 40 nucleotides in length. The apparent dissociation constant (app. K_d_) was calculated using Hill Slope equation with GraphPad Prism. Individual data points of three biological replicates are plotted. K_d_ values for Poly (C), Poly (A) and Poly (U) RNA could not be defined (n.d.). The K_d_ for Poly (G) is indicated. **(C)** Fluorescence polarization assays to determine the binding efficiency of recombinantly purified dMi-2 with FAM-labelled (GA)_20_ RNA (blue) and (CA)_20_ RNA (red). 5 nM RNA was used in each reaction and dMi-2 titrated from 0.0001 µM to 0.5 µM. Fluorescence polarization was measured in triplicates. Triplicate data points from two independent biological replicates are plotted. The apparent K_d_ was calculated using Hill Slope equation with GraphPad Prism. **(D)** EMSAs showing the binding of dMi-2 to G-quadruplex forming (G_4_A_4_)_4_ RNA (left panel) and to non-G-quadruplex forming (GA)_20_ RNA (right panel). 250 nM RNA was used for each reaction and dMi-2 was titrated from 0.01 μM (lanes 2) to 0.64 μM (lanes 8). RNA without protein was loaded in lanes 1. Each EMSA is a representative figure of at least two independent biological replicates. The gels were stained with SYBR gold. **(F)** Quantification of EMSAs in (D). (G_4_A_4_)_4_ RNA: blue and (GA)_20_ RNA: red. App. K_d_ was determined by Hill Slope equation using GraphPad Prism. Individual data points of two independent biological replicates are plotted.

To confirm this finding using a different method, we compared binding of dMi-2 to (CA)_20_ RNA with binding to (GA)_20_ RNA in a fluorescence polarization assay. In this experiment dMi-2 bound to (GA)_20_ RNA with an apparent K_d_ of 42 (+/-17) μM **(Figure 4C**). In contrast, binding of dMi-2 to (CA)_20_ RNA was so weak that we were unable to calculate a K_d_ value. We conclude that dMi-2 has a strong preference for binding to G-rich RNA sequences *in vitro*.

Some G-rich RNA molecules can form G-quadruplex structures and some chromatin regulators such as PRC2 display preferential binding to G-quadruplex-forming RNAs (Wang et al., 2017). We confirmed that the poly(G) 40mer used in Figure 4B does indeed form a G-quadruplex structure (**Supplementary Figure 4E**). We next compared binding of dMi-2 to a G-quadruplex-forming RNA ((G_4_A_4_)_4_ 40mer) with binding to an RNA fragment with very similar sequence composition that cannot form G-quadruplex structures ((GA)_20_ 40mer) **(Supplementary Figure 4F and 4G)**. dMi-2 bound with indistinguishable apparent affinities to both RNAs ((G_4_A_4_)_4_: K_d_ = 0.08 (+- 0.0) μM; (GA)_20_: K_d_ = 0.07 (+/- 0.01) μM) **(Figures 4D** and **4E)** indicating that dMi-2 binding to RNA is facilitated by G-rich sequences but not by G-quadruplex structures.

### A novel RNA binding domain in the N-terminus of dMi-2

dMi-2/CHD4 has several domains that have previously been demonstrated to interact with nucleic acids. These include the HMG box and adjacent sequences that interact with poly(ADP-ribose) and the chromodomains and the ATPase domain, both of which interact with nucleosomal DNA during the remodelling reaction (Bouazoune et al., 2002; Kovac et al., 2018; Murawska et al., 2011; Silva et al., 2016). We used a series of dMi-2 truncation mutants to assess if these domains also contribute to RNA binding in EMSAs. **Figure 5A** and **5B** show that neither of the regions with an established role in contacting nucleosomal DNA during the remodelling reaction exhibit significant RNA binding activity (compare lanes 2, 5 and 8). In contrast, fragments containing the N-terminal 376 amino acid residues, which harbour the HMG box, exhibited substantial RNA binding activity (compare lanes 2, 4 and 6). Moreover, removal of this N-terminal region resulted in a reduction of RNA binding (compare lanes 2, 3 and 7), suggesting that region 1-376 makes a major contribution to RNA binding *in vitro*.

**Figure 5:**
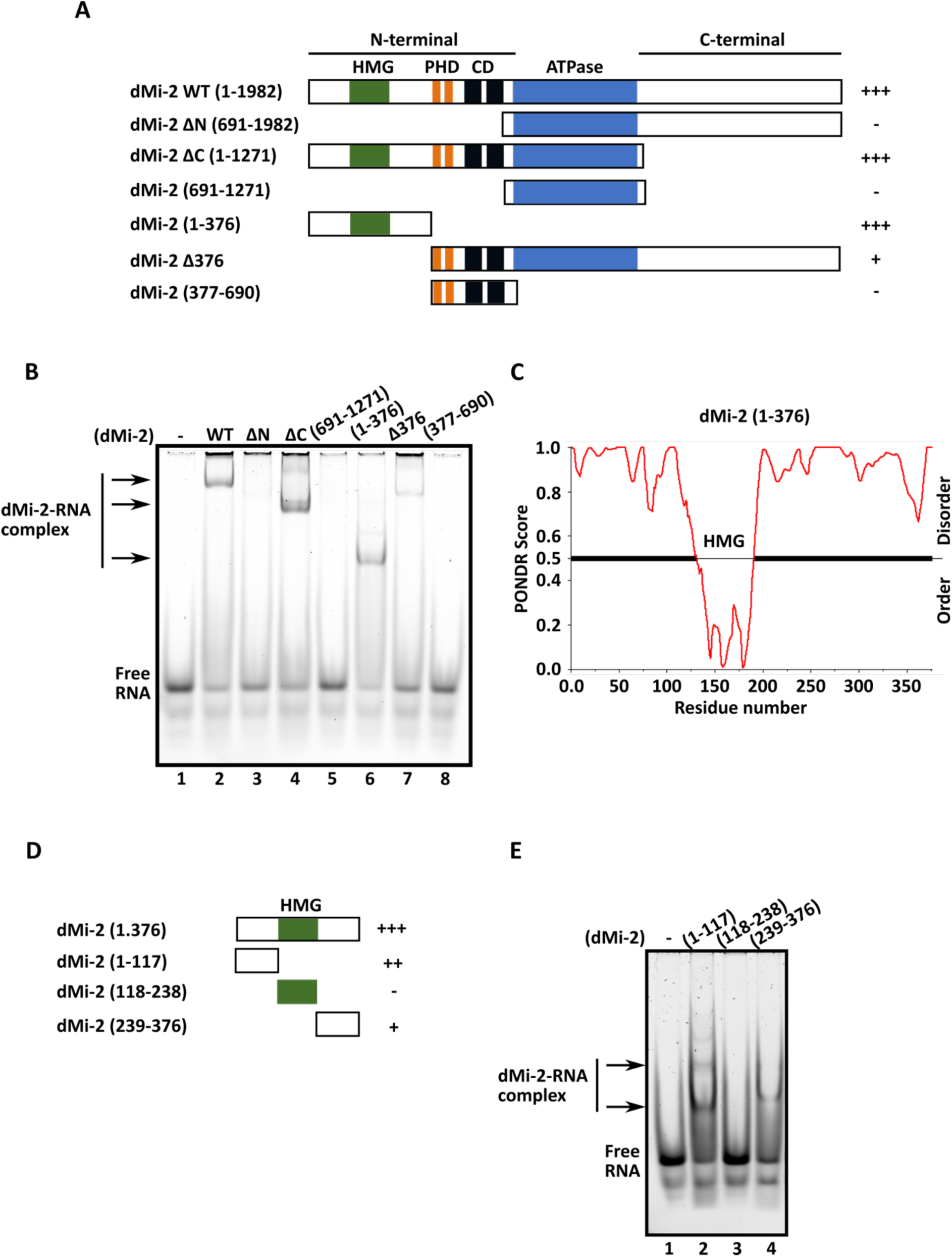
A novel RNA binding domain in the N-terminus of dMi-2. **(A)** Scheme representing wild type (WT) and truncation mutants of dMi-2. High mobility group box (HMG) (green), Plant homeodomains (PHD) (orange), chromodomains (CD) (black) and ATPase domain (blue) of dMi-2 are marked. Plus (+) and minus (-) signs on the right indicate the RNA binding properties of full length and truncation mutants with (+++) representing the best RNA binding. **(B)** EMSA of dMi-2 WT and truncation mutants. 250 nM of Hsp70Aa #6 RNA (see Figure 3A) was used for each reaction against 500 nM recombinantly purified proteins. RNA without protein was loaded as control in lane 1. The dMi-2-RNA complexes are shown by arrows on the left. **(C)** Prediction of intrinsically disordered regions (IDRs) in the N-terminal part of dMi-2. PONDR was used with optimal settings to predict IDRs. PONDR Score 1.0 (y-axis) is the highest score for the prediction of IDR while a PONDR Score of 0.0 indicates a well folded structure. **(D)** Scheme representing the truncation mutants of the dMi- 2 N-terminal region. HMG-box domain is denoted in green. Plus (+) and minus (-) signs on the right indicate the RNA binding properties of the truncation mutants of dMi-2 with (+++) being the highest RNA binding. **(E)** EMSA of dMi-2 (1-117) (lane 2), (118-238) (lane 3) and (239-376) (lane 4). 250 nM Hsp70Aa #6 RNA (see Figure 3A) was used for each reaction with 1.5 μM proteins. RNA without protein was loaded as control in lane 1. Arrows on the left show the dMi-2-RNA complexes.

The central HMG box in the 1-376 fragment is flanked by regions that are predicted to be disordered by PONDR **(Figure 5C)**. We further truncated this region to assess the importance of the HMG box for RNA binding (**Figure 5D**). Surprisingly, the isolated HMG box failed to interact with RNA in our assay **(Figure 5E)**. In contrast both disordered regions flanking the HMG box exhibited robust RNA binding activity.

Collectively, these results suggest that two N-terminal IDRs make a substantial contribution to RNA binding by dMi-2.

### G-rich RNA inhibits dMi-2- and CHD4-mediated nucleosome remodelling in vitro

We next asked if RNA can interfere with dMi-2 remodelling activity *in vitro*. We selected three 40mer RNAs and verified by EMSA that all three bound to dMi-2 *in vitro* **(Supplementary Figure 6 A–D)**. The (G_4_A_4_)_4_ 40mer bound dMi-2 with an apparent K_d_ of 0.08 (+/-0.02) μM) and the (GA)_20_ 40mer with an apparent K_d_ of 0.08 (+/-0.01) μM). The Hsp70Aa fragment, which contains a lower percentage of G nucleotides (30%), bound more weakly with an apparent K_d_ of 0.31 (+/-0.1) μM. To test remodelling activity, we used a DNA fragment with an end-positioned nucleosome. We used limiting amounts of dMi-2 to sensitize the assay for activation or inhibition. In agreement with our previous work, dMi-2 mobilized 25-30% of the histone octamer to a central position, resulting in a nucleosome that migrated more slowly during native gel electrophoresis **(Supplementary Figure 6E-G).** We then challenged the remodelling reaction by addition of increasing amounts of RNA. Addition of either of the two G-rich RNAs resulted in efficient inhibition of remodelling with a half maximal inhibitory concentration (IC50) of 140.10 (+/- 20.93) nM ((G_4_A_4_)_4_) and 152.65 (+/-52.25) nM ((GA)_20_), respectively (**Figure 6A)**. In contrast, the Hsp70Aa 40mer failed to impact the remodelling reaction even when used at high concentrations (1.5 μM). This result indicates that G-rich RNAs, which bind with high affinity to dMi-2, inhibit nucleosome remodelling. In contrast, RNAs that are not G-rich and bind dMi-2 with lower affinity are not capable of interfering with remodelling under our assay conditions.

**Figure 6:**
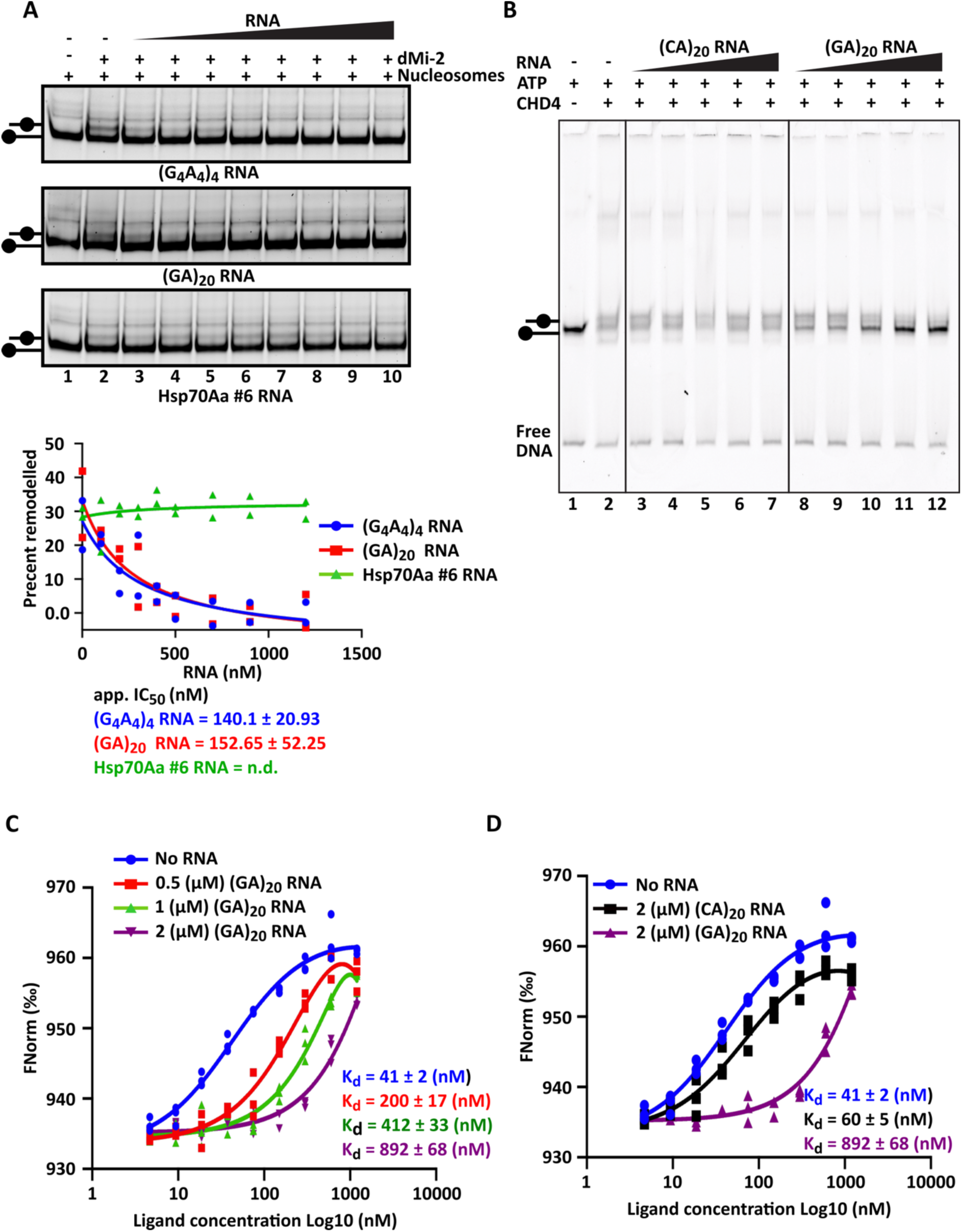
G-rich RNA inhibits dMi-2- and CHD4-mediated nucleosome remodelling *in vitro*. **(A)** Remodelling inhibition of dMi-2 by RNA. Remodelling reactions were set up using 15 nM of the end-positioned mononucleosomes with 80 bp of flanking DNA in each experiment as denoted by plus sign, except in controls (minus signs on the top). End-positioned mononucleosomes, lanes 1 are represented on the left of each panel. 50 nM dMi-2 was added in each reaction (plus sign), except in controls (minus sign on the top). RNAs (G_4_A_4_)_4,_ (GA)_20_ and (Hsp70Aa #6) were titrated from 0.1 to 1.2 μM, as indicated. Remodelled nucleosome, represented on the left, can be observed in lanes 2-6 of (G_4_A_4_)_4_ and (GA)_20_ and lanes 2-10 in (Hsp70Aa #6). Reactions were loaded on native polyacrylamide gels, stained with SYBR gold stain. Representative images of at least three independent biological experiments. Bottom panel, quantification of remodelling assays performed in presence of increasing amounts of (G_4_A_4_)_4_ RNA (blue), (GA)_20_ RNA (red) and Hsp70Aa #6 RNA (green). The IC_50_ values were calculated with non-linear regression using GraphPad Prism. Individual data points of two independent biological replicates are plotted. **(B)** Remodelling reactions of end-positioned mononucleosomes with 60 bp flanking DNA (0W60) with human CHD4 performed in presence of increasing amounts of (CA)_20_ RNA (lanes 3-7) and (GA)_20_ RNA (lanes 8-12). The un-remodelled and remodelled nucleosome positions are indicated on the left. Free DNA is indicated on left. For each reaction 100 nM nucleosomes and 10 nM CHD4, respectively, were used. RNA was titrated from 0.5 to 2 µM. The nucleosomal DNA carried a 5’-Cy3 label, and the gel was imaged with a Typhoon FLA 9000 laser scanner. **(C) and (D)** Microscale thermophoresis (MST) assays showing competition between RNA and nucleosomes for binding to human CHD4. CHD4 was titrated against 25 nM Cy5-labelled nucleosome. Following 10 min incubation on ice, the samples were loaded into Monolith NT.115 instrument for MST measurement. Each binding curve is generated via GraphPad Prism and is a fit to at least three technical replicates (shown as individual dots).

We then turned to CHD4, the human homologue of dMi-2, and asked if RNA-mediated inhibition of remodelling was conserved between *D. melanogaster* and *H. sapiens*. In the same gel-based remodelling assay, CHD4-mediated nucleosome remodelling was robustly inhibited by addition of the G-rich (GA)_20_ RNA **(Figure 6B)**. The (CA)_20_ RNA, on the other hand, failed to inhibit remodelling even at high concentrations (2 μM). We also measured inhibition of CHD4-mediated remodelling by (GA)_20_ RNA using a Real Time Fluorescence Assay (RTFA) ((Zhong et al., 2020); **Supplementary Figure 6H and 6I)**. Here, we used nucleosomes carrying an AF488 fluorophore on histone H2A and a BHQ1 quencher at one end of the DNA fragment **(Supplementary Figure 6H)**. Nucleosome remodelling relieves quenching of the AF488 fluorescence, allowing detection of remodelling in real time. Again, we observed an RNA concentration-dependent reduction of CHD4-mediated nucleosome remodelling **(Supplementary Figure 6I)**. Together, these results establish that inhibition of nucleosome remodelling by RNA is a property that is conserved between dMi-2 and CHD4.

### RNA directly inhibits binding of dMi-2/CHD4 to its nucleosomal substrate

What is the mechanism of inhibition of dMi-2/CHD4-mediated nucleosome remodelling by RNA? One possibility is that RNA engages with the active site of the enzyme or a nearby allosteric site to lower remodelling activity. Such a mechanism has been invoked to explain the inhibition of PRC2 histone methyltransferase activity by RNA (Zhang et al., 2019). However, given that we could not detect an interaction between the dMi-2 ATPase domain and RNA **(Figure 5),** we considered this possibility to be unlikely. Alternatively, the interaction of dMi-2/CHD4 with RNA might be incompatible with binding of the enzyme to the nucleosome. Consistent with this competition hypothesis, the data in Figure 6A show that RNAs that bind dMi-2/CHD4 with high affinity are more potent inhibitors of remodelling. In order to directly test if RNA can compete with CHD4 binding to the nucleosome, we used microscale thermophoresis. We measured the K_d_ of human CHD4 for a fluorescently labelled nucleosome to be 41 (+/-2) nM (**Figure 6C**). We then repeated the titration in the presence of (GA)_20_ RNA and observed a significant and concentration dependent reduction in the capacity of CHD4 to bind the nucleosome: the presence of 2 μM (GA)_20_ RNA reduced the affinity by ∼20-fold. Moreover, the same concentration of (CA)_20_ RNA reduced the K_d_ by less than two-fold (**Figure 6D**), demonstrating a strong sequence dependence of the ability of RNA to inhibit nucleosome binding by CHD4.

Taken together these results suggest that preferential binding of dMi-2/CHD4 to G-rich RNAs as well as inhibition of nucleosome remodelling by G-rich RNAs is shared between fly and man.

## Discussion

### dMi-2 associates with a broad range of mRNAs

Our study demonstrates that mRNA inhibits the function of an ATP-dependent chromatin remodeller of the CHD family. This inhibition involves the direct interaction of RNA with two novel IDRs in dMi-2. We identify two molecular mechanisms by which RNA binding counteracts dMi-2 function: dissociation of dMi-2 from chromatin and competition with the nucleosome substrate during the remodelling reaction. As discussed below, this mRNA-mediated activity allows a coordinated and simultaneous inhibition of several repressor complexes and thereby safeguards transcriptionally active regions from the establishment of repressive chromatin structures.

We reveal that dMi-2 is associated with thousands of mRNAs *in vivo* defining it for the first time as a *bona fide* RNA binding protein (RBP). This conclusion is supported by the efficient photo-crosslinking of dMi-2 to multiple mRNAs *in vivo* identified by iCLIP2, by the co-immunoprecipitation of formaldehyde-crosslinked dMi-2/mRNA complexes and by the robust interaction of dMi-2 with numerous RNA fragments detected in *in vitro* EMSA and FP assays. We have identified a novel RNA binding region in the N-terminus of dMi-2 that comprizes two predicted IDRs (residues 1-117 and 239-376) flanking an HMG box that has previously been shown to bind DNA and poly(ADP-ribose) (Silva et al., 2016). These IDRs contain regions rich in basic amino acids that might interact with the phosphate backbone of RNA. However, given that formation of dMi-2/RNA complexes is not strongly affected by high salt concentrations we expect that non-ionic interactions with RNA bases will make a substantial contribution to RNA binding.

The RNA-binding IDRs that flank the HMG box are also present in human and *C. elegans* homologs of dMi-2 (CHD4 and LET-418, respectively) indicating an RNA-binding activity that is conserved across 550 million years of evolution. These IDRs are not resolved in the recently reported cryo-EM structure of the CHD4-nucleosome complex, suggesting either that they do not adopt a well-defined conformation in presence of the nucleosome substrate or that the interactions that they form are with a higher-order (>1 nucleosome) chromatin substrate (Farnung et al., 2020). We speculate that the IDRs might adopt more ordered conformations when they encounter RNA.

Although the IDR-containing region displays potent RNA binding activity, it is important to note that dMi-2 deletion mutants lacking this region retained weak RNA binding *in vitro* (Figure 5). This result suggests that dMi-2 contains several RNA binding sites that are distributed across a wider region. These could increase binding to one RNA molecule by providing multiple contact points or allow the simultaneous binding of multiple RNA molecules.

### RNA binding preference of dMi-2

dMi-2 displays a preference for binding G-rich RNA fragments *in vitro.* The basis for this selectivity is currently unclear and will require structural analyses of dMi-2/RNA complexes. dMi-2’s preference for G-rich sequences is not entirely recapitulated *in vivo*. Although G residues are overrepresented at the crosslinking site and the +1 position, the surrounding region is not particularly G-rich. In fact, A and C are strongly overrepresented at the +4 and +5 positions, respectively. It is likely that RNA binding *in vivo* is influenced by other dMi-2 complex subunits. Indeed, the *C. elegans* homolog of dMEP-1, which interacts with dMi-2 to form the dMec complex, has previously been shown to bind RNA (Belfiore et al., 2002; Kunert et al., 2009).

In addition, other RBPs might facilitate the interaction of dMi-2 with RNA via protein-protein interactions. RBPs that are preferentially associated with the 3’ ends or the polyA tail of mRNAs would be of particular interest given that dMi-2 preferentially crosslinks to the 3’ ends of mRNA *in vivo*. An analysis of the RNA binding properties of 24 human chromatin regulators by fRIPseq has revealed a strong relationship between CHD4- and PABP-bound RNAs (Hendrickson et al., 2016). The association of both CHD4 and PBAP with mRNA was highest over the 3’ ends of transcripts. Moreover, CHD4 was found to bind almost exclusively to exons. This finding mirrors our results and suggests that the observed mRNA binding preferences are fundamental features of this remodeller. It is tempting to speculate that an interaction with PABP contributes to the dMi-2/RNA interaction *in vivo*. Indeed, PABP was recovered in a screen for CHD4 interactors (Hoffmeister et al., 2017). An involvement of general RBPs in mediating dMi-2 binding would help to explain why dMi-2 associates with thousands of mRNAs that differ in sequence and secondary structure *in vivo*.

### RNA inhibits nucleosome remodelling

The ability of RNAs to inhibit dMi-2 mediated nucleosome remodelling correlates with their affinity for binding to dMi-2. This observation suggests that it is RNA binding to dMi-2, rather than RNA binding to the nucleosome substrate, that directly modulates remodelling. RNA binds to PRC2 at an allosteric site close to the catalytic centre, suggesting that RNA binding results in a change in conformation (Zhang et al., 2019). In contrast the major RNA binding region of dMi-2 (aa 1-376) is far from its catalytic domain in primary sequence. Still, it is possible that the N-terminus of dMi-2 interacts with the ATPase domain in the three-dimensional structure of the enzyme even though this region could not be resolved in the structure of the CHD4-nucleosome complex (Farnung et al., 2020). Nevertheless, our finding that RNA can compete with the nucleosome substrate for binding to CHD4 suggests a different inhibition mechanism. We propose that RNA binding to dMi-2 is incompatible with the simultaneous binding to nucleosomes, at least for RNA molecules of 40 nucleotides or longer. It is conceivable that dMi-2 associated RNA interferes with the interactions of chromodomains and the ATPase domain with nucleosomal DNA; these interactions are essential for the remodelling reaction (Farnung et al., 2020; Kovac et al., 2018).

SWI/SNF nucleosome remodelling complexes have recently been demonstrated to interact with repressive, long non-coding RNAs such as Xist and Evf2 (Cajigas et al., 2015; Jegu et al., 2019). Xist binding results both in eviction of SWI/SNF complexes from the inactive X chromosome and in inhibition of remodelling and ATPase function (Jegu et al., 2019). It is remarkable that both activating SWI/SNF and repressive dMi-2/CHD4 remodelling complexes are inhibited by RNA, and yet seem to respond to different types of RNA: long non-coding RNA that function to repress transcription in the case of SWI/SNF complexes and protein-coding mRNAs that emanate from actively transcribed genes in the case of dMi-2/CHD4.

### RNA antagonizes both dMi-2/CHD4 and PRC2

PRC2 and dMi-2/CHD4-containing NuRD collaborate to establish repressive chromatin structures. NuRD facilitates PRC2-mediated H3K27 methylation by deacetylating H3K27ac (Reynolds et al., 2012b). Moreover, NuRD increases local nucleosome density which might further contribute to PRC1-mediated chromatin compaction (Bornelov et al., 2018). The effects of RNA binding to PRC2 and NuRD are two-fold: (1) counteracting chromatin binding and (2) inhibition of enzymatic activity. Thus, two different types of enzymes - a histone methyltransferase and an ATP-dependent nucleosome remodeller - are both inhibited by RNA. This RNA-mediated mechanism provides an elegant strategy to simultaneously target the two cooperating repressor complexes - PRC2 and NuRD - and in doing so to safeguard actively transcribed genes from the establishment of repressive chromatin structures (**Figure 7**).

**Figure 7:**
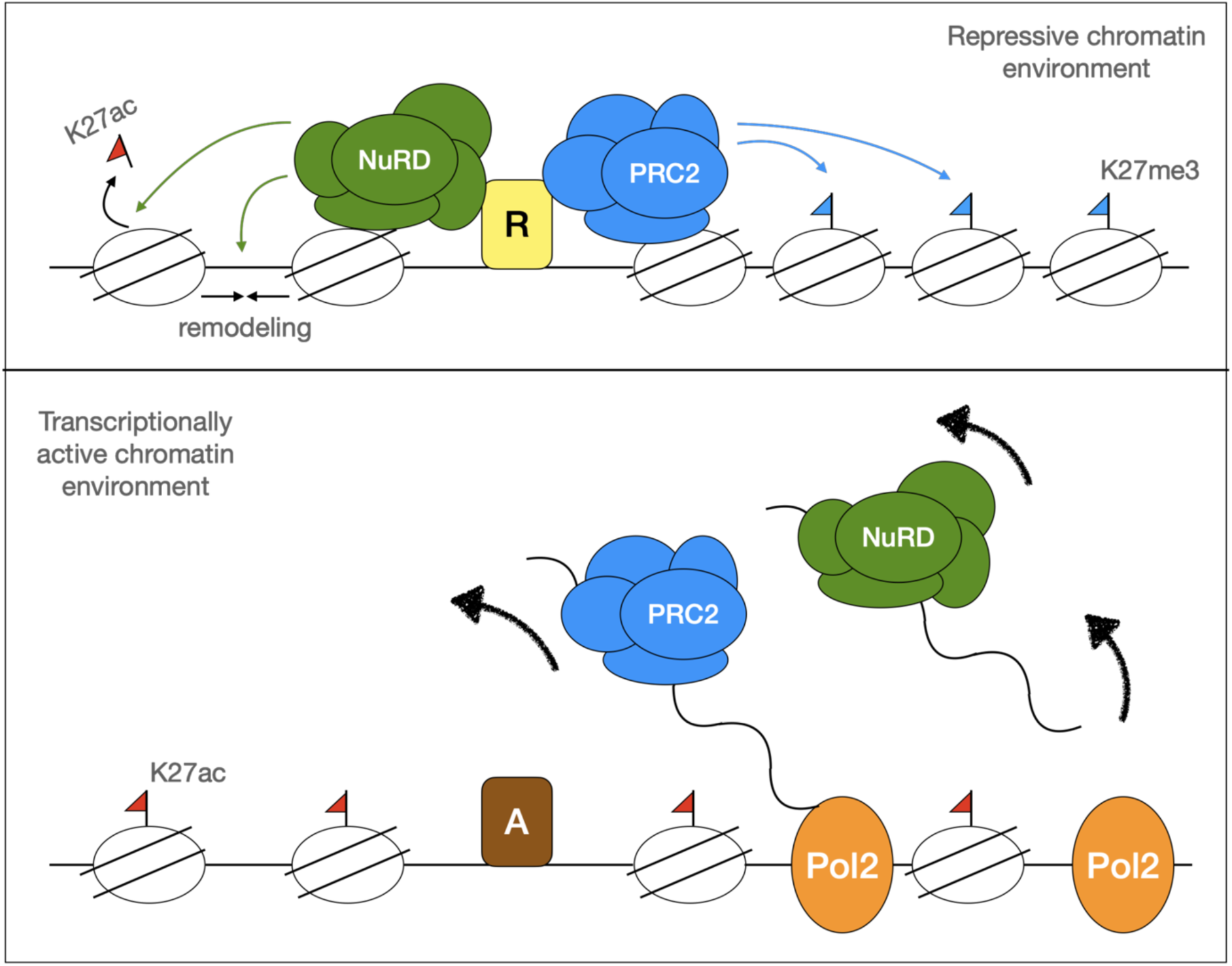
RNA antagonizes NuRD and PRC2 to protect transcriptionally active chromatin environments. Upper panel: In the absence of active transcription, NuRD and PRC2 associate with chromatin by binding to DNA bound repressors (R) or by direct interaction with nucleosomes. NuRD remodells nucleosomes to increase local nucleosome density and deacetylates nucleosomes (e.g. K27ac) to facilitate PRC2 mediated nucleosome methylation. PRC2 generates K27me3 marked nucleosomes and together with PRC1 establishes repressive chromatin structures. Lower panel: During active transcription, NuRD and PRC2 are removed from chromatin by RNA. Nucleosome density and K27me3 can no longer be maintained. Activators (A) recruit histone acetyltransferases and demethylates (not shown) to remove repressive and add active histone modifications. As long as the region is actively transcribed NuRD and PRC2 cannot access chromatin. In addition, their remodelling and histone methylation activities are directly inhibited by RNA.

In summary, our work supports a general and conserved role of mRNA in the coordinated control of multiple chromatin regulating complexes. This activity constitutes a negative feedback loop in which mRNA, the ultimate product of “active chromatin”, directly engages and counteracts two important complexes that generate repressive chromatin structures. We envisage that this mechanism stabilizes an active chromatin region until transcription rates are reduced by the dissociation of activating transcription factors or the binding of transcriptional repressors to enhancers and/or promoters. Reduced mRNA production would then provide a window of opportunity for PRC2 and NuRD to (re)engage with chromatin and to (re)establish repressive chromatin structures.

## Acknowledgements

I.U. and A.B. were supported by DFG TRR81/A01, YZ and JPM were supported by NHMRC APP1126357, OR was supported by the DFG Research Training Group (RTG) 2355 (project 325443116). We would like to thank Dr. Chen Davidovich, Monash University, EMBL-Australia, and Dr. Manuel Beltran, UCL, London, and Dr. Manuel Beltran, UCL, London, for technical advice.

## Author contributions

IU, JPM, AB conceived the study; IU, OR, JPM, AB designed experiments; IU and OR performed iCLIP2; AN and TS curated sequencing data; CT and HRC analysed iCLIP2 data; JL analysed RNA-seq data; IU and MJ performed RIP-qPCR and FLAG IP; IU, MJ and OV performed FP assays; IU and YZ performed nucleosome remodelling assays; YZ performed MST and RTFA assays; IU performed ChIP-qPCR, RT-qPCR, cell fractionation, EMSA and CD assays; MG and RH synthesized RNA; IU, CT, YZ, MJ, OR, JL, HRC, JPM and AB discussed and interpreted results; IU, OR, JPM and AB wrote the manuscript.

## Materials and Methods

### iCLIP2

Individual nucleotide-resolution UV-crosslinking and immunoprecipitation 2 was performed as described in (Buchbender et al., 2020). Briefly, S2 cells expressing GFP-tagged dMi-2 (Lenz et al., 2021) and untagged S2 cells were washed with cold 1× PBS. Protein-RNA cross-linking was performed at 254 nm UV light (300 mJ/cm^2^) using a Stratagene Stratalinker. Cells were harvested by centrifugation for 5 min at 2,000 rpm at 4°C. Cell pellets were resuspended in 4 volumes (vol.) of RIPA buffer (50 mM Tris-HCl pH 7.4, 1% NP-40, 0.1% SDS, 150 mM NaCl, 5 mM EDTA), incubated for 10 min on ice and centrifuged at 13,000 rpm for 20 min at 4°C.

#### DNase and partial RNase digestion

To the whole cell extract, 2 vol. of RQ-1 buffer (40 mM Tris-HCl pH 8.0, 10 mM MgSO_4,_ 1 mM CaCl_2_) was added. Subsequently, TURBO^TM^ DNase (1:500 vol.) and RNaseOUT (1:1,000 vol.) were added to the extract and mixed gently. RNase I (Ambion, AM2294) was added to a final concentration of 10^-3^ (11.764 U/μL), 10^-4^ (1.307 U/μL) and 10^-5^ (0.145 U/μL), respectively. Samples were incubated at 37°C for 3 min with 800 rpm shaking and immediately placed on ice for 3 min. To each sample, 20 μL (1:50 vol.) of 5 M NaCl was added and samples centrifuged at 14,800 rpm for 5 min at 4°C. The extracts were transferred to fresh tubes.

#### Immunoprecipitation (IP)

The GFP-trap beads were washed 2× with 1 mL cold TBS-T buffer (50 mM Tris pH 7.4, 150 mM NaCl, 0.05% Tween-20). To each sample, the washed and equilibrated GFP-trap beads were added, and samples were incubated for 2 h at 4°C while rotating. After IP, the samples were centrifuged at 2,000 rpm for 5 min at 4°C and the flow-through was discarded. The beads were washed 4× with TBS1000-T buffer (50 mM Tris pH 7.4, 1000 mM NaCl, 0.05% Tween-20) and 2× with PNK buffer (70 mM Tris-HCl pH 7.5, 10 mM MgCl_2_, 0.05% NP-40). Beads were transferred to fresh Eppendorf tubes after the third wash.

#### On-bead phosphatase treatment

To the beads, 40 μL of 20× phosphatase mix (80 μL 10× phosphatase buffer (Fermentas), 30 μL SAP (1 U/μL, Fermentas), 10 μL RNaseOUT (40 U/μL, Invitrogen) and 680 μL DMPC treated water) was added and incubated at 37°C for 20 min. Beads were washed 2× with TBS400-T buffer (50 mM Tris pH 7.4, 400 mM NaCl, 0.05% Tween-20). Eppendorf tubes were changed after second wash and beads washed 2× with PNK buffer.

#### On-bead 3’-end RNA linker ligation

To the beads, 25 μL of 1× ligase mix (1.5 μL 3’-linker (100 pmol/μL), 2.5 μL of 10× T4 RNA-ligase buffer (Thermo Scientific), 0.65 μL RNA-ligase (10 U/μL, Thermo Scientific), 0.25 μL RNaseOUT (40 U/μL, Invitrogen), 2.5 μL BSA (1 mg/mL, Thermo Scientific) and 17.60 μL DMPC treated water) was added and incubated at 16°C in a thermomixer overnight. Samples were washed 2× with PNK buffer.

#### On-bead PNK treatment

To the beads, 20 μL of 1× PNK mix (2 μL of 10× PNK buffer (NEB), 2 μL ATP (SCP801; 12.33 pmol/μL), 0.5 μL T4 PNK Enzyme (1000 U/μL, NEB), 0.5 μL RNaseOUT (40 U/μL, Invitrogen) and 15 μL DMPC treated water) was added and incubated at 37°C for 20 min in a thermomixer. Beads were washed 1× with cold TBS-T and 1× with cold PNK buffer. All buffers were freshly supplemented with 1 mM DTT, RNase inhibitor (unless RNaseOUT is supplied) and protease inhibitor cocktail.

#### SDS PAGE and Blotting

The beads were mixed with 20 μL of 2× LDS loading buffer supplemented with 1mM DTT. Samples were denatured for 10 min at 70°C while shaking and loaded on 4-12% Bis-Tris gel along with a pre-stained size marker (Thermo Scientific) and electrophoresed at 200 V for 5 h. Proteins were transferred to a nitrocellulose membrane by blotting overnight at 15 V at 16°C. The protein size marker bands were drawn on a plastic bag using a radioactive pen after placing the plastic bag on top of the membrane. The membrane was exposed, whilst in the plastic bag, to X-ray film for 16 h. The X-ray films were developed and the autoradiograms were used to detect the protein-RNA complex of interest. The corresponding protein-RNA area was marked on the nitrocellulose membrane.

#### RNA isolation

The selected area on the nitrocellulose membrane containing the protein-RNA complex of interest was cut along with the corresponding area in the control lane. To the nitrocellulose membrane pieces, 400 μL of PNK buffer was added and 20 μL of Proteinase K (20 mg/mL). Samples were incubated at 37°C for 20 min with 1,000 rpm shaking. 400 μL of PK buffer (100 mM Tris-HCl pH 7.4, 50 mM NaCl, 10 mM EDTA and 1% SDS) and PK-Urea buffer (PK buffer containing 7 M urea) were added and samples were incubated for a further 20 min at 55°C with shaking at 1,000 rpm.

To each sample, 800 μL of RNA-grade phenol-chloroform-isoamyl alcohol (25:24:1) was added and samples were incubated at 30°C for 5 min while shaking at 1,000 rpm. Samples were centrifuged at 13,000 rpm for 5 min at room temperature using phase-lock gel (5PRIME). The aqueous layer was transferred to a new tube without disturbing the gel. RNA was precipitated by adding 1 μL glycoblue (Ambion, 9510), 80 μL of 3 M sodium acetate (NaOAc) pH 5.5 and 0.7 vol. ice-cold isopropanol, mixed and incubated overnight at -80°C. Samples were centrifuged at 14,800 rpm for 20 min at 4°C, the supernatant was discarded, the RNA pellet washed with 0.9 mL of 80% ethanol, centrifuged again at 14,800 rpm for 5 min. The RNA pellet was air-dried for 3 min and dissolved in 5 μL of RNase-free water and transferred to a PCR tube.

The RNA was reverse transcribed and iCLIP library was prepared as detailed in (Buchbender et al. 2020) and sequenced.

### Western blot

Samples to be analyzed by Western blot (WB) were mixed with SDS loading buffer, denatured at 95°C for 5 min and loaded on SDS polyacrylamide gel along with a size marker and electrophoresed. The proteins were transferred to activated polyvinylidene difluoride (PVDF) membrane at 400 mA (milliamperes) for 2 h in cold conditions. The membrane was blocked with 5% milk in PBS-T (1× PBS pH 7.4, 0.1% (v/v) Tween-20) for 30 min at room temperature while shaking and then incubated over night with primary antibody in 5% milk in PBS-T at 4°C on a shaker. The membrane was washed 3× with PBS-T for 5 min each at room temperature on a shaker and incubated with the respective secondary antibody in 5% milk in PBS-T at room temperature for 2 h on a shaker. The membrane was again washed 3× with PBS-T for 5 min each at room temperature on a shaker. Western blot signals were detected by chemiluminescence using HSP substrate (Milipore, WBKLS0500). The following primary antibodies were used: Rat monoclonal anti-GFP antibody (Chromotek, 80626001AB, 1:5,000), Mouse monoclonal anti-tubulin anti-body (Merck, 3272293, MAB3408, 1:10,000), Rabbit polyclonal anti-dMi-2 anti-body (custom made, 0.5 mg ml^-1^, 1:10,000), Rabbit anti-E(z) antibody (1:3,000), anti-dLSD1, Rabbit polyclonal anti-dRPD3 (Abcam 1767, 1:10,000) and Rabbit polyclonal anti-H3 antibody (Abcam 1791, 1:10,000). The following secondary anti-bodies were used: Sheep polyclonal anti-mouse HRP (Amersham, NA931, 1:30,000), Goat polyclonal anti-Rat HRP (Invitrogen, 31470, 1:1,000).

### RNA-IP-qPCR

80 million cells were cross-linked with formaldehyde at a final concentration of 1% (v/v) and slowly shaken on a plate rocker for 10 min at room temperature. To quench this reaction, glycine was added to a final concentration of 240 mM and incubated for 5 min at room temperature while shaking. Cells were collected and spun down at 1,000 rpm for 10 min at 4°C. The supernatant was removed and the pellet was washed with 10 mL cold 1× PBS, pH 7.4. The supernatant was removed and the pellet was resuspended in 800 µl FA buffer (50 mM HEPES-KOH pH 7.6; 140 mM NaCl; 1% (v/v) Triton X-100; 0.1% Sodium deoxycholate) with freshly added protease inhibitors, 1 mM DTT and RNAsin (Thermo Scientific, 100 Units (U)/mL). The cells were lysed on ice for 15 min. After the lysis, the cells were sonicated for 10 min at the high intensity setting (30 sec on/off) in a Bioruptor (Diagenode). The whole cell extract was adjusted to 25 mM MgCl_2_ and 5 mM CaCl_2_ and 1 µL DNAse I (30 U) was added. The solution was incubated for 10 min at room temperature and stopped by adding EDTA to a final concentration of 10 mM. To remove cell debris, the extract was centrifuged at 13,000 rpm for 10 min at 4°C. 100 µL of the supernatant was removed and stored at -20°C (input sample). 500 µL of the supernatant was used for precipitation. The GFP Trap resins (ChromoTek, gta10) were equilibrated 3× with 1 mL FA buffer supplemented with protease inhibitors, 1 mM DTT and RNAsin (100 U/mL). 30 µL of GFP Trap resins were used for precipitation. The IP was performed overnight at 4°C on a spinning wheel.

The resins were spun down at 2,000 rpm for 4 min at 4°C and the supernatant was removed. The resins were washed 5× with 1 mL FA buffer and 2× with 1 mL TE buffer pH 8.0 (10 mM Tris-HCl, 1 mM EDTA) with freshly added protease inhibitors, 1 mM DTT and RNAsin (100 U/mL). Before the last wash with TE buffer, resins were transferred into fresh Eppendorf tubes. After the last washing step, the supernatant was removed, 100 µL of RIP elution buffer (100 mM Tris-HCl pH 8.0, 10 mM EDTA; 1% (w/v) SDS) with RNAsin (40 U/mL) was added and samples incubated for 10 min at room temperature on a rotating wheel. Samples were spun down at 2,000 rpm for 4 min at room temperature and the supernatant was transferred into new Eppendorf tubes. Again, 100 µL of RIP elution buffer with RNAsin (40 U/mL) was added and samples incubated in a thermomixer at 1,000 rpm for 10 min at 65°C. Samples were spun down with the same settings as before and the supernatant was transferred into the same microfuge tube as the previous elution. 100 µL of RIP elution buffer with RNAsin (40 U/mL) was added to the ‘input’ sample to adjust the volume to 200 µL. Eluates and ‘input’ samples were adjusted to 200 mM NaCl and 20 µg of proteinase K (10 mg/ml) was added. Samples were incubated at 42°C for 1 h and then at 65°C for 2 h for decrosslinking. The RNA was purified with the PeqGOLD Total RNA Kit (PeqLab) or RNeasy Mini Kit (Qiagen) following the manufacturer’s instruction and an additional DNAse I digestion was performed on the column provided in the kit. The purified RNA was then reverse transcribed into complementary DNA (cDNA) with the SensiFAST™ complementary cDNA synthesis Kit (Bioline) and used in qPCR. After cDNA synthesis, the cDNA was diluted 1:20 in nuclease-free water.

cDNA synthesis program:

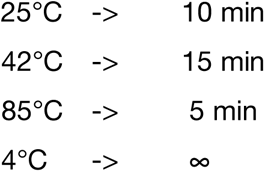

qPCRs were performed using SensiFast^TM^ SYBR® Lo-ROX kit (Bioline). A master mix for each primer pair was prepared following the manufacturer’s instructions. 15 µL of master mix was used per well and 5 µL of cDNA or nuclease-free water control were added to the appropriate wells. The 96-well plate was closed and centrifuged shortly. The RT-qPCR was performed under the following conditions-

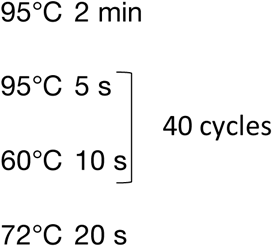

For the RIP experiments, the percentage of the input was calculated as-

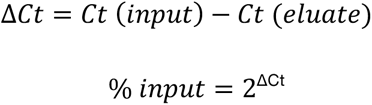

For plotting the values on a graph, the standard deviation was calculated as:

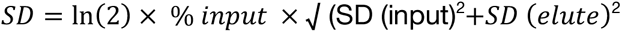

### Chromatin immunoprecipitation-qPCR

100 million cells were seeded in 15 cm dishes and the protein-DNA contacts were cross-linked using 1% (v/v) formaldehyde (Polysciences, 04018-1) for 10 min at room temperature with shaking. The formaldehyde was quenched with 240 mM glycine (final concentration) for 10 min at room temperature with shaking. The cells were harvested and washed 2× with cold PBS pH 7.4, the supernatant was discarded. The cell pellet was lysed with 1 mL lysis buffer (50 mM Tris-HCl pH 8.0, 10 mM EDTA and 1% SDS, supplemented with protease inhibitors cocktail and 1 mM DTT) and incubated 10 min on ice. The cell lysate was sheared with Biorupter (Diagenode) with 3 cycles of 30 seconds ON/OFF for 10 min each at high power. The lysate was centrifuged at 14,800 rpm for 15 min at 4°C, the supernatant (chromatin) transferred to fresh Eppendorf tubes and the pellets discarded. Protein A beads (Invitrogen) were washed 3× with low salt buffer (20 mM Tis-HCl pH 8.0, 2 mM EDTA, 150 mM NaCl, 1% Triton X-100, 0.1% SDS) and blocked overnight in blocking buffer (2 mG/mL BSA, 2% fish skin gelatin in low salt buffer).

The chromatin (130 μL per ChIP) was diluted 10-fold with ChIP-IP buffer (16.7 mM Tris pH 8.0, 167 mM NaCl, 1.2 mM EDTA, 0.01% (w/v) SDS, 1.1% (w/v) Triton X-100) and pre-cleared with pre-equilibrated and blocked protein A beads (80 μL slurry per ChIP) for 1 h on a rotating wheel at 4°C. Pre-cleared chromatin was centrifuged at 14,800 rpm for 10 min at 4°C. The supernatant was transferred to fresh Eppendorf tubes, 13 μL were removed and stored at - 20°C (input). To the pre-cleared chromatin, 50 μL (1:1) pre-equilibrated and blocked (just as protein A beads), GFP-Trap beads (chromoTek) were added and samples incubated overnight on a rotating wheel at 4°C. The samples were centrifuged at 2,000 rpm for 5 min at 4°C, supernatant was discarded and beads were washed 3× with low salt buffer, 3× with high salt buffer (20 mM Tis-HCl pH 8.0, 2 mM EDTA, 500 mM NaCl, 1% Triton X-100, 0.1% SDS), 1× with LiCl buffer (10 mM Tris-HCl pH 8.0, 1 mM EDTA, 250 mM LiCl and 0.1% NP-40) and 1× with TE buffer (10 mM Tris pH 8.0, 1 mM EDTA). For each washing step, samples was incubated on a rotating wheel at 4°C for 10 min. After the last wash, beads were resuspended in 700 μL TE buffer, transferred to fresh Eppendorf tubes and centrifuged at 2,000 rpm for 5 min at 4°C. The supernatant was discarded. 250 μL of elution buffer (100 mM NaHCO_3_ and 1% (v/v) SDS) was added to each sample and incubated for 20 min on a rotating wheel at room temperature, centrifuged at 2,000 rpm for 5 min at room temperature. The supernatant was transferred to fresh Eppendorf tubes. To the beads, 250 μL of elution buffer was added followed by incubation for 20 min at room temperature on a rotating wheel and then at 95°C for 10 min. Samples were centrifuged at 14,800 rpm at room temperature for 5 min. The supernatant was pooled with the first eluate. To the input samples, 486 μL of elution buffer was added. To decrosslink, 20 μL of 5 M NaCl was added and samples were incubated at 65°C overnight with 800 rpm shaking. Protein was digested by adding 10 μL of 0.5 M EDTA, 20 μL of 1 M Tris-HCl pH 6.8 and 2 μL of Proteinase K (10 mg/mL) and samples were incubated at 45°C for 1 h with 800 rpm shaking. DNA was purified using Qiaquick PCR purification Kit (Qiagen) following the manufacturer’s protocol. The DNA was eluted with elution buffer provided in the kit. The DNA was diluted 1:4 with nuclease free water and 5 μL were used for each qPCR reaction. qPCR reactions were set up and analyzed as described above (RIP-qPCR).

### Triptolide treatment

S2 cells were seeded in 15 cm dishes (for ChIP) or 6-well plates (for RT-qPCR) and 10 μM triptolide (dissolved in DMSO) or an equal volume of DMSO added to the media. Cells were incubated for 3 h and were processed for ChIP-qPCR as described above or for RT-qPCR as described below.

### RT-qPCR

Triptolide treated and DMSO treated cells were harvested and RNA extracted using PeqGold Total RNA isolation kit (PeqLab). RNA was reverse transcribed to cDNA using SensiFAST cDNA synthesis kit (Bioline). The cDNA was diluted 1:5 with nuclease free water and 5 μL were used per qPCR reaction. qPCR reactions were set up using sensiFAST SYBR Lo-ROX kit (Bioline). The qPCR was performed as described above for RIP-qPCR.

### RNase A treatment and cell fractionation

RNase A treatment and cell fractionation was performed as explained in (Beltran et. al., 2016). Briefly, 2-4 × 10^7^ cells per condition were aliquoted into 15 mL Falcon tubes. The cells were washed with 1× PBS pH 7.4 at room temperature. Cells were permeabilized with 10 mL of Buffer P (0.05% Tween-20 in 1× PBS pH 7.4) and incubated at 4°C for 10 min with intermittent agitation. The cell suspension was centrifuged at 1,200 rpm for 2 min at 4°C and the supernatant was discarded. The cell pellet was resuspended in 950 μL of 1× PBS pH 7.4 (room temperature). The sample was split into two vials, 450 μL each. One was labelled as ‘sample’ and another ‘mock’. The ‘sample’ vial was taken to a different room while the ‘mock’ vial was processed on an RNase-free bench. To the ‘sample’ vial, 50 μL of 10 mg/mL RNase A (Applichem) was added while to the ‘mock’ vial, 50 μL of 10 mg/mL BSA was added. Each vial was incubated on separate rotating wheels at room temperature for 30 min. Both the vials were centrifuged at 1,200 rpm for 2 min at 4°C. The supernatant was discarded. The cell pellet was washed by pipetting up and down with 10 mL cold 1× PBS.

The cell suspension was centrifuged as earlier. The pellet was resuspended in 1 mL ice-cold buffer A (10 mM HEPES pH 7.9, 10 mM KCl, 1.5 mM MgCl_2_, 0.34 M sucrose, 10% glycerol and supplemented with protease inhibitor cocktail and 1 mM DTT). From this step, 2.5 μL/mL RNase inhibitor – RNaseOUT (40U/μL, Ambion), was added to each buffer for ‘mock’ sample.

Triton X-100 was added to a final concentration of 0.1% (v/v), the tubes were flicked to mix and incubated on ice for 5 min. To isolate the cytoplasmic fraction, the samples were centrifuged at 3,600 rpm for 4 min at 4°C. The supernatant was separated, cleared by centrifugation at 14,100 rpm for 15 min at 4°C and stored at -20°C. The nuclei were washed 2× with 1 mL buffer A (resuspended gently by flicking). The nuclei were lysed with 250 μL of buffer B (3 mM EDTA, 0.2 mM EGTA, supplemented with protease inhibitor cocktail and 1 mM DTT) and mixed by gently flicking the tube. For the ‘mock’ sample, the buffer B was additionally supplemented with RNaseOUT. Both samples were incubated on ice for 30 min. The nucleoplasm was separated by centrifugation at 4,100 rpm for 4 min at 4°C. The nucleoplasm was stored at -20°C. The chromatin pellet was washed twice with 1 mL of buffer B and centrifuged at 4,100 rpm for 4 min at 4°C (mixed gently by flicking the tube). The final chromatin pellet was resuspended in 200 μL of 1× Laemmli buffer supplemented with *β*−mercaptoethanol (*β* - Me). The chromatin samples were sonicated for 7 cycles (30 s ON, 30 sec OFF, at High output) in a Bioruptor (Diagenode). The samples were heated to 95°C for 5 min and stored at -20°C. The cytoplasm, nucleoplasm and chromatin fractions were loaded on 8-12% SDS-gel and electrophoresed. For the ‘mock’ sample, care was taken to clean the bench, the pipettes and all equipment with RNaseZap.

### Electrophoretic mobility shift assay (EMSA)

In a 10 μL standard reaction, DNA/RNA was incubated with the protein of interest in EMSA buffer (40 mM KCl, 20 mM Tris pH 7.6, 1.5 mM MgCl_2_, 0.5 mM EGTA, 10% glycerol, 200 ng/uL BSA). EMSA buffer was freshly supplemented with protease inhibitor cocktail, 1 mM DTT and 40U/mL of RiboLock RNase inhibitor (40U/μL, Thermo Scientific), for RNA EMSA only, for 15 min at room temperature. The samples were loaded on a pre-run 5% native polyacrylamide gel and electrophoresed at 100 V for 1 h. The gel was stained with 1× SYBR gold stain (Thermo Scientific) and imaged with Bio-Rad Chemi Doc Touch system. The bands were quantified by Bio-Rad Image Lab software and after subtracting the background intensity, the bound fractions were calculated as-

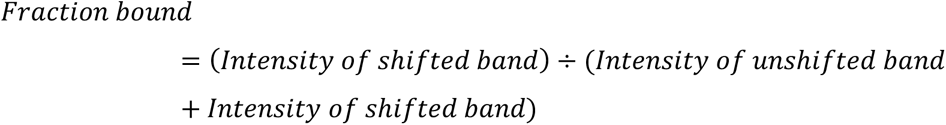

Percent bound was calculated as-

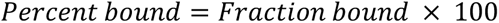

The apparent dissociation constant (app. K_d_) was calculated using the hill slope equation as

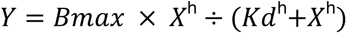

Where,

Bmax = the maximum specific binding in the same unit as Y, Kd = ligand concentration needed to achieve a half-maximal binding at equilibrium in the same unit as X and h = hill slope.

### FLAG immunopurification

The Sf9 cells expressing the FLAG-tagged protein of interest were harvested by centrifugation at 3,200 rpm for 25 min at 4°C. The supernatant was discarded and the cell pellet washed with 30 mL of cold 1× PBS (Phosphate buffer saline), pH 7.4. The centrifugation was repeated as earlier and the supernatant was discarded. The cellular pellet was resuspended and lysed with buffer BC-250 (0.4 mM EDTA pH 8.0, 20 mM HEPES pH 7.9, 10% (v/v) glycerol, 250 mM NaCl and freshly supplemented with protease inhibitors and 1 mM DTT), transferred to 50 mL Falcon tubes and incubated on ice for 15 min. The lysate was snap-frozen in liquid nitrogen and thawed. The freeze-thaw cycle was repeated. The lysate was then cleared by centrifugation at 10,400 rpm for 75 min at 4°C. The supernatant was transferred to a pre-chilled 50 mL Falcon tube and the cell debris was discarded. An aliquot from the whole cell extract was removed as ‘input’, mixed with 5× SDS loading buffer and stored at -20°C. Anti-FLAG M2 agarose affinity gel (Sigma, A2220) were equilibrated by washing 3× with buffer BC-250 (10× bead vol.). To the extract, pre-equilibrated anti-FLAG M2 agarose affinity gel was added and samples were incubated at 4°C for 4 h on a rotator. After immunoprecipitation, the solution was transferred to Bio-Rad Econo-pac gravity-flow columns (Bio-Rad, #732-1010), the flow-through was collected. The beads were washed with 40× bead volume of BC-250, BC-500 (BC buffer with 500 mM NaCl), BC-1000 (BC buffer with 1,000 mM NaCl), BC-500, BC-250 and BC-100 (BC buffer with 100 mM NaCl) buffer in that order. Each buffer was freshly supplemented with protease inhibitor cocktail and 1 mM DTT. The purified protein of interest was eluted with 2× the volume of beads with elution buffer (FLAG peptide (5 mg/mL) diluted 1:20 in BC-100 buffer). The elution was repeated twice more. The elutes were pooled and concentrated in pre-chilled Centricons (Thermo Scientific).

The concentrated elute was dialyzed with buffer BC-50 (BC buffer with 50 mM NaCl and 5% glycerol) and loaded on a 5 mL HiTrap Q Fast Flow anion exchange chromatography column (Sigma) using an ÄKTA^TM^ Pure FPLC (GE Healthcare). The flow through was collected. The bound fraction was washed with BC-50 buffer (20 column volumes) and gradually eluted with BC-2M buffer (BC buffer with 2 M NaCl and 5% glycerol). The flow through and elute sample fractions were loaded on denaturing polyacrylamide (Coomassie stain) and 5% native polyacrylamide (SYBR Gold stain) gels and electrophoresed to verify the purity of the fractions. The respective purified protein fractions were pooled, dialyzed to BC-100 buffer, aliquoted, snap frozen and stored at -80°C.

### Native Polyacrylamide Gel Electrophoresis

EMSA reactions were loaded on native polyacrylamide gels. A 100 mL solution of 5% native polyacrylamide gel was prepared by mixing 12.5 mL of Rotiphorese Gel 40 (19:1, acrylamide:bisacrylamide) (Roth) with 5 mL of 10× Tris-Boric acid-EDTA (TBE) buffer (1 M Tris, 0.9 M boric acid and 0.01 mM EDTA). 400 μL of ammonium persulfate (APS) (Roth) and 80 μL tetramethylethylenediamine (TEMED) (Roth) were added and the solution poured in a glass chamber. The gel was allowed to polymerize at room temperature for 30 min, then pre-run for 1 h at 100 V. After the EMSAs were loaded, the gel was electrophoresed for 1 h at 100 V at RT in 0.5× TBE buffer. The gel was incubated in a 1× SYBR gold (Thermo Scientific) staining solution for 5 min at room temperature with shaking. The gel was washed 2× with deionized water and imaged with a Bio-Rad ChemiDoc Touch system.

### Denaturing PAGE for RNAs

RNA samples were loaded on denaturing polyacrylamide gels to verify the quality of the RNAs. The gels were poured in the same manner as the native gels described above, but were additionally supplemented with 8% urea. The RNA samples were mixed with RNA loading buffer (Thermo Scientific), heated to 65°C for 5 min, instantly cooled for 1 min on ice and loaded on a pre-run denaturing gel and electrophoresed at 100 V for 30 min in 0.5× TBE buffer. The gels were processed as described above.

### Fluorescence Polarization Assay

For FP assays, a protein dilution series was performed, leading to a protein concentration ranging from 0.5 µM to 0.0001 µM in 50 µL BC-100 buffer (0.4 mM EDTA pH 8.0; 20 mM HEPES pH 7.9; 100 mM NaCl; freshly added protease inhibitors and 1 mM DTT). The RNA working stock was diluted with BC-100 buffer to a concentration of 10 nM. 50 µL of the 10 nM RNA working stock was added into each well, leading to a final concentration of 5 nM in a 100 µL reaction. The plate was shortly spun down and fluorescence polarization measured at time point 180 min at 26°C. “Excitation” was set to “Filter” for measuring using a Spark® 20M multimode reader (Tecan). The following formulae were used for calculating the app. K_d_ and the EC50 values:

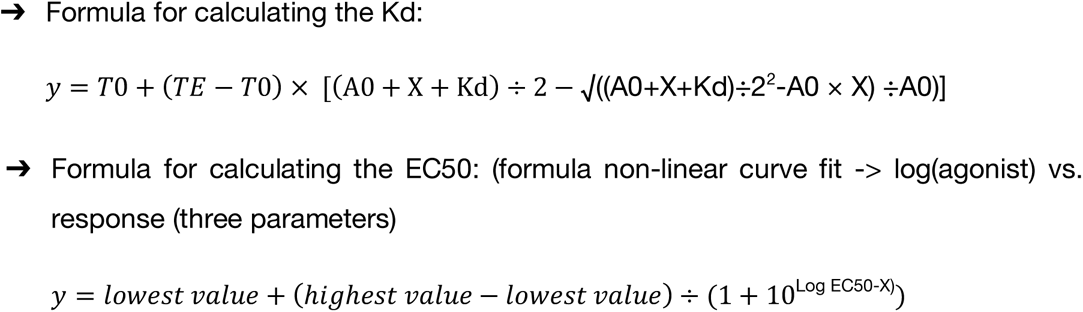

### Circular dichroism (CD) spectroscopy

RNA (10 μM) in 170 μL volume was allowed to fold as described in (Wang et al. 2017). CD spectra were measured in a 0.1 cm path-length quartz cell. The measurements were taken from 220 nm to 350 nm with a data pitch of 0.1 nm and scanning speed of 50 nm/min and a band width of 1 nm. The cell holder temperature was regulated and the cuvette chamber was flushed with dry Nitrogen (N_2_) gas. Spectra were measured at temperatures indicated in the figures. Three replicates were taken for each measurement and the mean data were analyzed by using CAPITO CD analysis and plotting software (Wiedemann et al. 2013). The data were smoothened with Savitzky-Golay filter and mean residue ellipticity (*θ* in grad cm^2^ dmol^-1^) was plotted.

### Nucleosome remodelling assay

A typical nucleosome remodelling reaction was set up in a volume of 10 μL. 50 nM end-positioned mono-nucleosomes with 80 bp flanking DNA were mixed with different concentrations of dMi-2 in BC-100 buffer (0.4 mM EDTA pH 8.0, 20 mM HEPES pH 7.9, 10% (v/v) glycerol, 100 mM NaCl and freshly supplemented with protease inhibitors and 1 mM DTT). Reactions were incubated for 5 min at 26°C, then, 2 mM ATP (Sigma) and 6 mM MgCl_2_ were added and reactions were incubated at 26°C for 45 min. Reactions were stopped by adding competitor plasmid DNA (5 μg/reaction) and samples were incubated on ice for 10 min and loaded on a 5% native polyacrylamide gel and electrophoresed at 100 V for 3 h in 0.5× TBE buffer. Gels were stained with 1× SYBR gold staining solution, imaged with Bio-Rad ChemiDoc Touch system and bands were quantified with Bio-Rad Image Lab software. For effects of RNA on remodelling activity of dMi-2, RNA was titrated in increasing concentrations before the addition of ATP and MgCl_2_. The IC50 was calculated in GraphPad Prism software using the following equation-

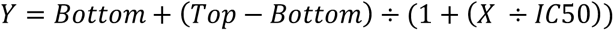

Where IC50 is the concentration of agonist that gives a response half way between Bottom and Top. Top and Bottom are plateaus in the units of the Y axis.

Remodelling reactions with hCHD4 were performed in the following buffer: 50 mM Tris-HCl pH 7.5, 50 mM NaCl, 3 mM MgCl2 and 1 mM ATP. Samples were incubated at 37°C for 1 h, stopped with competitor DNA and loaded onto the gel and processed as described above.

### Microscale thermophoresis (MST) Assay

CHD4 was concentrated to ∼2 µM and buffer exchanged into the dilution buffer (50 mM HEPES pH 7.5, 50 mM NaCl, 3 mM MgCl_2_) using Amicon Ultra-0.5 mL Centrifugal Filters (100 K MWCO, Merck Millipore). For each MST assay, a set of nine titration points was prepared by 1:2 serial dilution using the same dilution buffer. The diluted CHD4 were then mixed with 50 mM HEPES pH 7.5, 100 ng/µL BSA, 0.5% (v/v) glycerol and 25 nM 46^Cy5^w60 nucleosome. Following 10 min incubation on ice, the samples were loaded into premium coated capillary tubes (Nanotemper) before undergoing MST in a Monolith™ NT.115 instrument. Thermophoresis was conducted at 80% LED power, and 20% MST power. Thermophoresis data were then analyzed with MO. Affinity Analysis software v2.3 and plotted using GraphPad Prism software.

### Real time fluorescence assay (RTFA)

Nucleosomes assembled from 0^BHQ1^w60 DNA and AF488-labelled histone octamer were used for RTFAs. The assay was performed at 37°C and the fluorescence intensity was monitored at 526 nm. The reaction mixtures (typically 50 µL) containing 60 nM nucleosome, 10 nM CHD4 in 50 mM Tris-HCl pH 7.5, 50 mM NaCl and 3 mM MgCl_2_ were loaded into the wells of a Corning^®^ black nonbinding surface half-area 96- well plate, which was then subsequently put into a Tecan^®^ Infinite M1000 Pro plate reader pre-warmed to 37°C. Reactions were monitored for 5 min prior to the addition of 1 mM ATP by the injector unit of the plate reader, and the fluorescence changes were continued to be monitored for another 25 min, with data points recorded every 30 s. The data were normalized with a nucleosome-only control.

### Bioinformatic Pipeline

The raw iCLIP2 sequencing reads were assigned to the different libraries by the specific barcode part of the unique molecule identifier (UMI) and deduplicated based on total sequence identity with an in-house script. UMIs were removed from the read sequences and added to the read identifier. Additionally, poly(A) tails with at least eight consecutive A’s were removed from reads and trimmed reads longer than 19 nucleotides were aligned to the *Drosophila melanogaster* genome (dm6) using bwa-mem (0.7.17-r1188, <u>arXiv:1303.3997v2</u>) with minimum score output (-T) set to 19. Alignments were deduplicated based on UMIs and mapping position using UMI-tools (1.0.1, doi:10.1101/gr.209601.116) dedup functionality with default parameters. Only reads mapping uniquely to the genome were considered for further analysis while the rest of the reads were discarded. The uniquely mapped reads were attributed to transcript annotations based on their genomic coordinates using featureCounts from the SubRead (v1.6.4, doi:10.1093/bioinformatics/btt656) software suite for non-overlapping features (-O) and only counting strand specific alignments (-s 1).

Sample Correlation- The read counts from the two biological replicates were checked for correlation by calculating the squared Pearson Correlation Coefficient for paired samples. For visualization a linear model was fitted using the same metric. (there is no analysis at all. There is a simple formula for the Pearson Correlation Coefficient for paired samples and you square that to get R²).

Metagene Analysis- A metagene representation was generated based on read starts from both libraries using bamCoverage, calculateMatrix and plotProfile from the deepTools suite (3.4.3, doi:10.1093/nar/gkw257) with options “-bs 1 --ignoreDuplicates --minMappingQuality 1 --exactScaling --Offset 1”. Additionally, a detail graphic for the TTS of all transcripts with iCLIP reads was generated by manually plotting cross link positions derived from alignment start position minus one within +/-500 nucleotides of the TTS. Transcripts were sorted by mapped number of reads and their proximity to TTS. Relative density of cross links was calculated for the respective region.

Intron-Exon coverage- The cumulative coverage across the last intron-exon junction of all transcripts with at least two exons was determined using a custom script and plotted for +/- 10 nucleotides around the junction.

DE-Seq analysis- We compared iCLIP crosslinking results and mRNA expression level, as determined by RNA-Seq (Lenz *et al*., 2021) to identify mRNAs that preferentially associate with dMi-2. RNA-seq reads were mapped in the same manner as iCLIP reads but without prior trimming. Uniquely mapping reads were then counted using featureCounts as described above. In contrast to iCLIP, RNA-Seq read counts were normalized to FPKM (fragments per kilobase in transcript per million reads in library) to account for the difference in protocols (there is one read per cross-linked transcript molecule in iCLIP, while RNA-Seq can produce multiple reads per molecule especially for longer transcripts). DESeq2 (doi:10.1186/s13059- 014-0550-8) was used to identify mRNAs more frequently captured in iCLIP experiments than expected from their proportional abundance in RNA-Seq data. We chose a threshold for adjusted p-value of < 0.05 and putative log fold difference of iCLIP vs. RNA-Seq reads of > 4 to identify mRNAs that we regard as significantly enriched.

Position relative to cross-link site- Genomic sequences +/-200 nucleotides around all identified cross-linking sites were extracted and absolute as well as relative base content per relative position to the cross-linking site calculated to identify base specific biases at the cross-linking site. Relative base content per position was converted to percent of the median content for each base within the 400-nucleotide window. Additionally, a sequence logo was generated with genomic sequences of +/-20 nucleotides around the crosslinking site using kpLogo (v1.1, doi:10.1093/nar/gkx323) and parameters “-gapped -startPos 21”.

## Supplementary Figure Legends

**Supplementary Figure 1:**
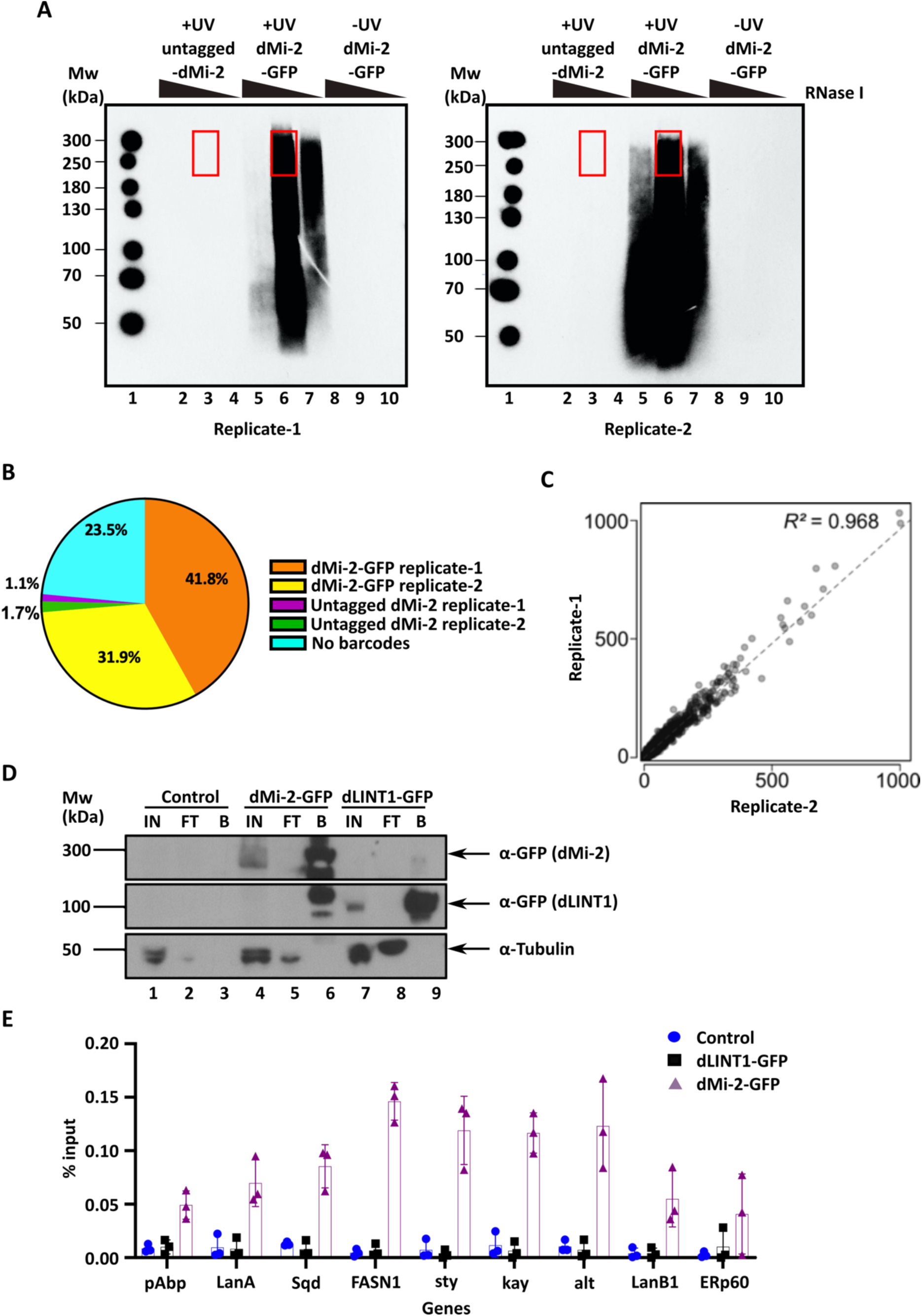
dMi-2 associates with thousands of mRNAs *in vivo*. **(A)** iCLIP2 autoradiographs, replicate-1 (left panel) and replicate-2 (right panel). Same cell lines and treatment conditions as mentioned in Figure 1A. The red boxes indicate the regions of the membranes that were cut out for iCLIP2 sequencing library preparation. Molecular weight is indicated on the left in lane 1 of each replicate. **(B)** Distribution of iCLIP2 reads. dMi-2-GFP replicates-1 and -2 are indicated in orange and yellow, respectively. The control untagged-dMi-2 replicates-1 and -2 are indicated in green and violet, respectively. 23.5% of the reads, indicated in cyan, had no barcode and were not included in our downstream analysis. **(C)** Squared Pearson correlation coefficient (R^2^) of the iCLIP2 data sets from replicate-1, on x-axis, and replicate-2, on y-axis. The R^2^ = 0.968 indicates high correlation between the two replicates. **(D)** Western blot analysis of untagged control and GFP tagged cells lines. Input (IN) from each cell line was loaded in lanes 1, 4 and 7, respectively. Flow through (FT) was loaded in lanes 2, 5 and 8, respectively. GFP-beads (B) were loaded after GFP-immunoprecipitation in lanes 3, 6 and 9, respectively. Western blot membranes were probed with anti-GFP antibody. Anti -Tubulin antibody was used as a loading control. Molecular weights are indicated on the left. **(E)** fRIP-qPCR validation of dMi-2 associated RNAs indicated in Figure 1C. RIP-qPCRs were performed with extracts from control (blue), dLINT1-GFP- (black) and dMi-2-GFP- expressing (purple) cell lines. Percent of input is plotted. Individual data points of three independent biological replicates are shown.

**Supplementary Figure 2:**
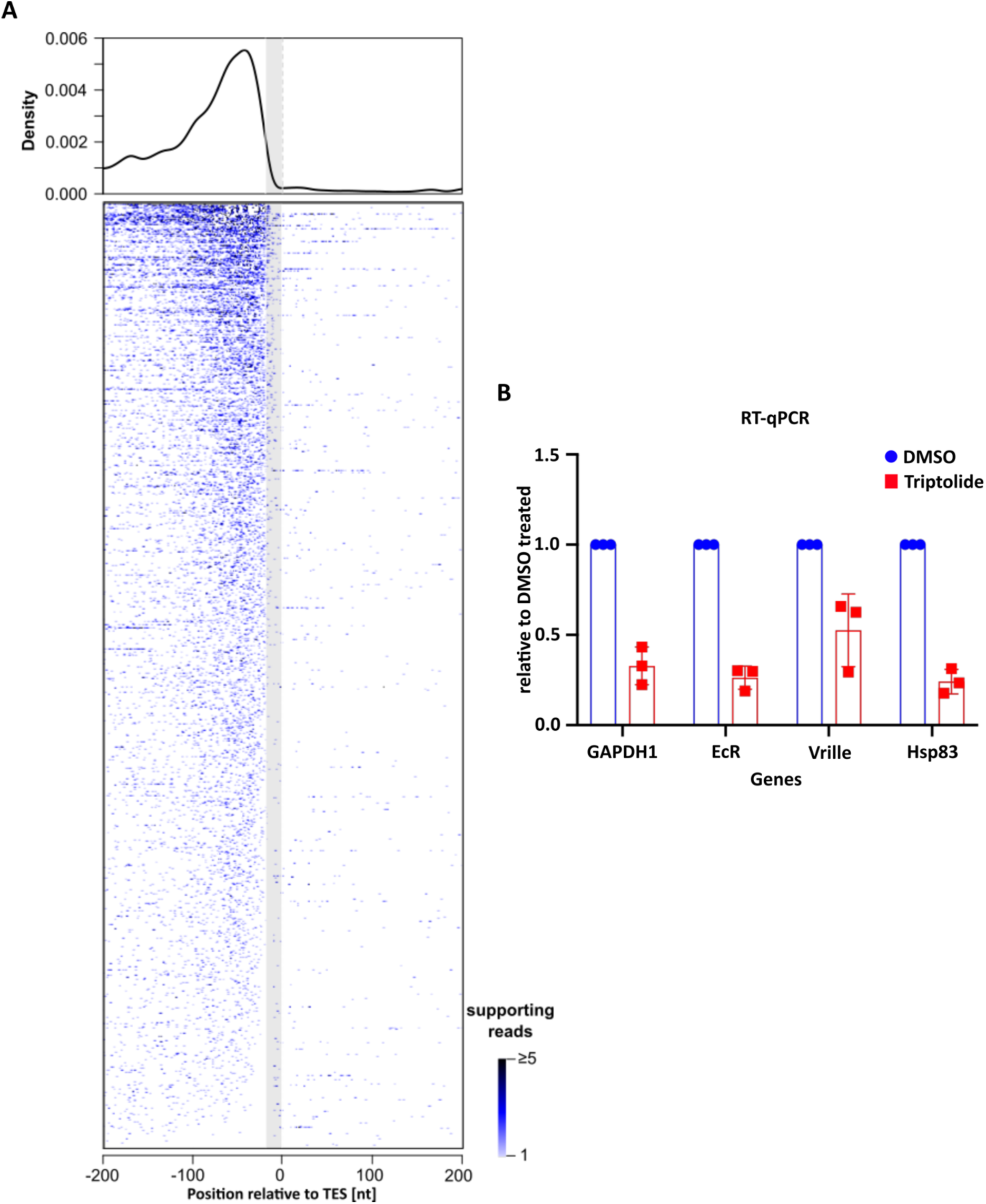
RNA binding does not play a major role in recruiting dMi-2 to chromatin. **(A)** Genes with iCLIP2 reads are aligned at the transcription end site (TES = 0). Crosslinking sites are indicated in blue. The cumulative crosslinking density for all genes is plotted on top. The grey shaded area from position -19 to -1 indicates a part of the transcript where reads are extremely unlikely to be mapped due to the minimum length of trimmed reads of 18 nucleotides. Crosslinks in this zone are mostly due to partial alignments containing a poly(A) signal, very rarely read-through events or false positive mappings. The colour code on the right indicates the number of supporting reads for each crosslinking site from 1 to ≥ 5. **(B)** S2 cells were treated with 10 μM triptolide (red) or an equal volume of DMSO (blue) for 3 hours. RNA was isolated from each sample and cDNA synthesized. qPCRs were performed using primers for the genes indicated. The expression of each gene was measured relative to DMSO control which was set to 1. Individual data points of three independent biological replicates are plotted as fold change. Error bars indicate standard deviation between three biological replicates.

**Supplementary Figure 3:**
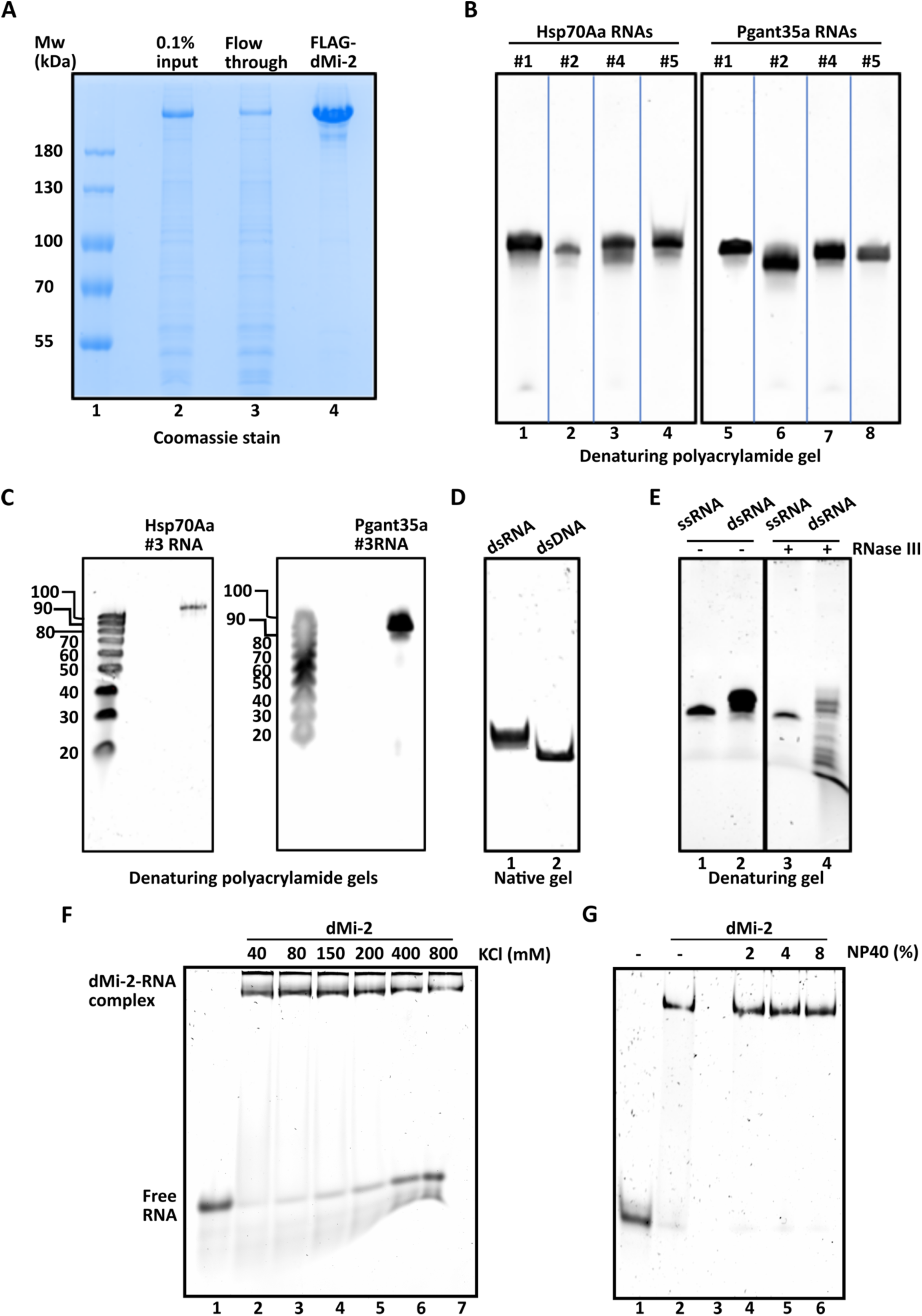
dMi-2 has broad RNA binding activity *in vitro*. **(A)** Immunopurification of FLAG tagged dMi-2 protein. 15 μl of eluted dMi-2 (lane 4) was subjected to SDS PAGE along with 0.1% input (14.4 μg) and 14.4 μg of flow through (lanes 2 and 3, respectively). The gel was stained with Coomassie Brilliant Blue. Molecular weight marker was loaded in lane 1. **(B)** 150 ng of RNAs derived from Hsp70Aa and Pgant35a genes (Figure 3A) were separated on urea denaturing polyacrylamide gels (lanes 1-4 and 5-8, respectively) and visualized with SYBR gold staining. **(C)** #3 RNAs (100-mer ssRNAs) from Hsp70Aa and Pgant35a along with ssRNA size marker are loaded on urea denaturing polyacrylamide gels. **(D)** 100 bp dsRNA and DNA generated from Hsp70Aa gene (lanes 1 and 2, respectively) were loaded on a native polyacrylamide gel and electrophoresed. **(E)** To verify the double stranded nature of dsRNA, it was treated with RNase III (lane 4) which specifically cuts dsRNA. Hsp70Aa #3 RNA, a ssRNA of equal length, was used as a control (lane 3). Untreated controls were loaded in lanes 1 and 2. The specific degradation of dsRNA upon RNase III treatment (lane 4), confirms the double stranded nature of dsRNA. **(F)** EMSA showing binding of dMi-2 to RNA in increasing salt conditions. 250 nM of Hsp70Aa #6 RNA (Figure 3A) and 500 nM of recombinantly purified dMi-2 was used for each reaction. RNA without protein was loaded in lane 1 as control. dMi-2-RNA complex is indicated on the left. **(G)** EMSA showing binding of dMi-2 to RNA in increasing NP40 concentrations. 50 nM of Hsp70Aa #3 RNA and 173 nM of dMi-2 was used for each reaction. No NP40 was added in lanes 1 and 2. Lane 3 was left empty. EMSA samples were loaded on native polyacrylamide gels and stained with SYBR gold.

**Supplementary Figure 4:**
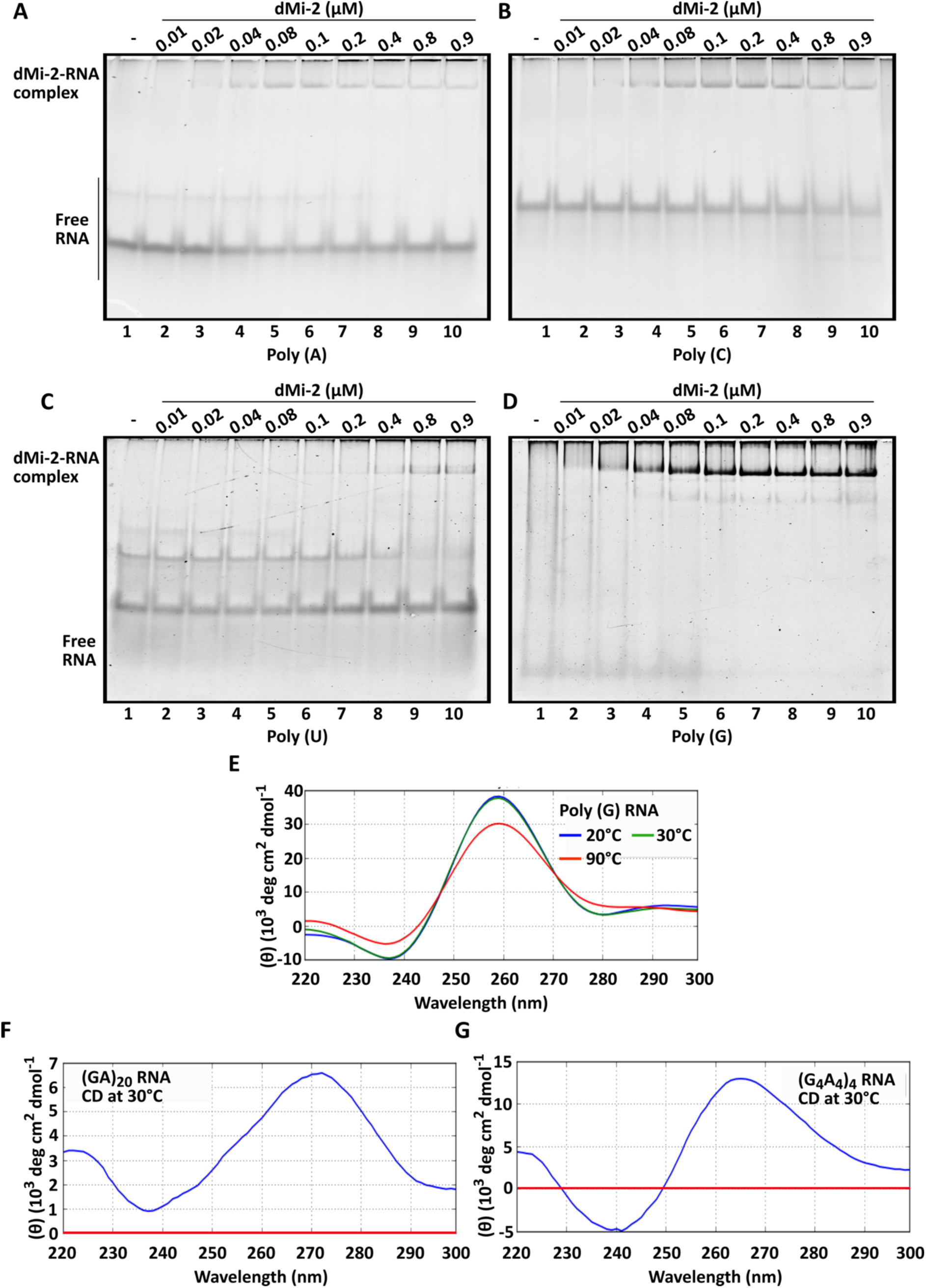
dMi-2 preferentially binds G-rich RNAs. **(A-D)** EMSAs of Poly (A), (C), (U) and (G) RNAs, each 40-mer in length. 250 nM RNA was used in each reaction and dMi-2 was titrated from 0.01 to 0.9 μM. Samples were loaded on native polyacrylamide gels. The gels were stained with SYBR gold. **(E-G)** Circular dichroism (CD) spectroscopy of Poly (G) RNA, (G_4_A_4_)_4_ and (GA)_20_ RNA, each 40-mer in length. CD spectra were measured at temperatures as indicated and between 220 to 300 nm. The CD values were analysed and plotted by CAPITO CD analysis and plotting software and are shown as mean residue ellipticity (θ in grad cm^2^ dmol^-1^). Poly (G) and (G_4_A_4_)_4_ RNAs form characteristic G-quadruplex absorbance pattern with negative peaks between 237-241 nm and positive peaks between 260-265 nm. At least 3 measurements were taken at each temperature and mean values plotted. The red lines in (F) and (G) denote the zero (0) value on Y-axis.

**Supplementary Figure 5:**
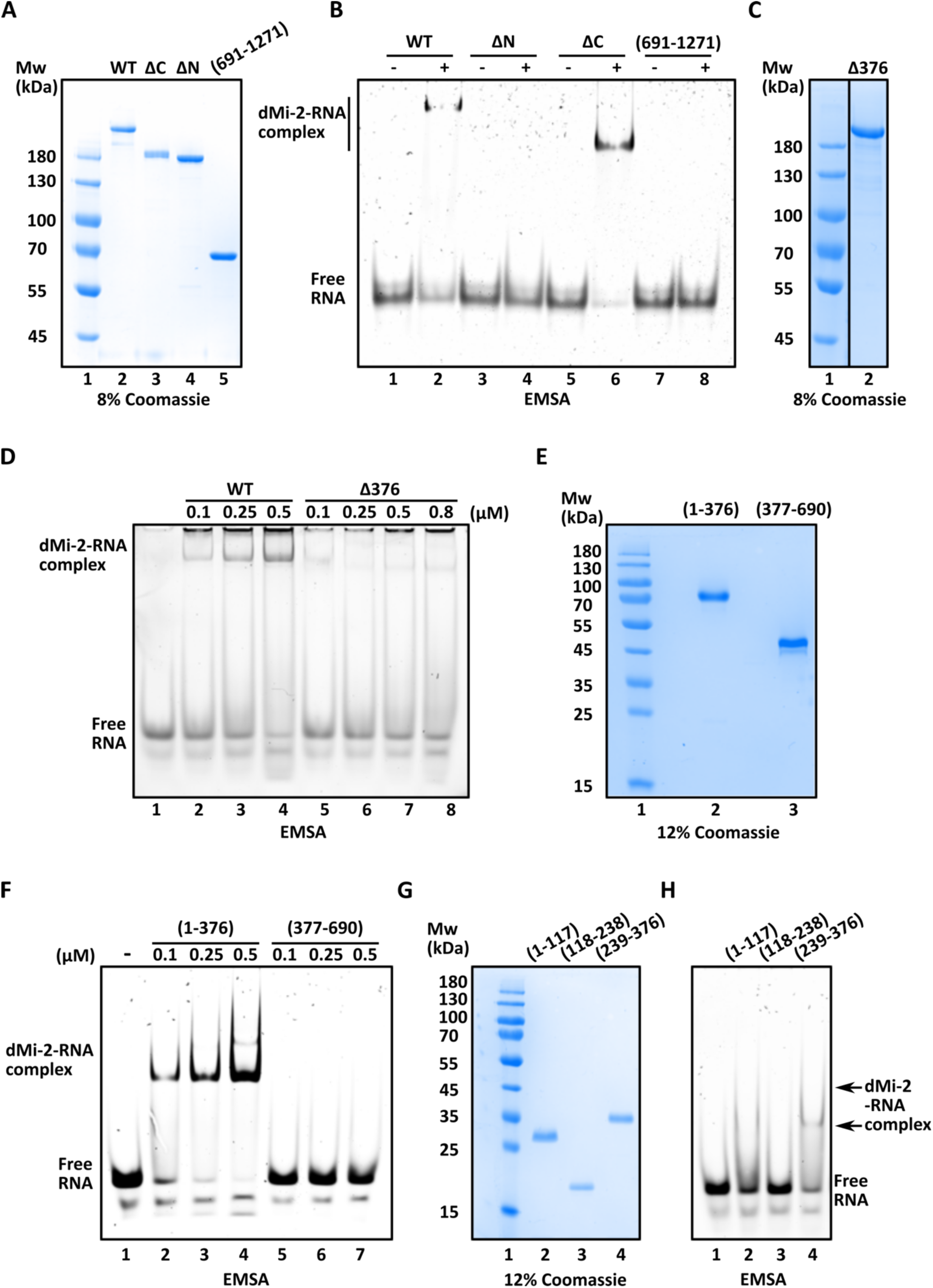
A novel RNA binding domain in the N-terminus of dMi-2. FLAG-tagged dMi-2 Wild type (WT) and its truncation mutants were purified by FLAG immunopurification. The protein samples were loaded on SDS gels in **(A)**, **(C)**, **(E)** and **(G)**. Molecular weight markers were loaded in lane 1 for each gel. The gels were stained with Coomassie Brilliant Blue. Additional EMSAs of dMi-2 WT and its truncation mutants in **(B)**, **(D)**, **(F)** and **(H)**. In **(B)**, 50 ng of Hsp70Aa #3 RNA (Figure 3A) was used for each reaction and 173 ng each of dMi-2 WT (lane 2), dMi-2 ΔN (lane 4), dMi-2 ΔC (lane 6) and dMi-2 (691-1271) (lane 8). In **(D)**, 250 nM Hsp70Aa #6 RNA was used for each reaction with titrations of dMi-2 WT or dMi-2 Δ376, as indicated. In **(F)**, 250 nM Hsp70Aa #6 RNA was used in each reaction and dMi-2 (1-376), lanes 2-4 or dMi-2 (377-690), lanes 5-7, were titrated from 0.5 to 2 μM. In **(H)**, 250 nM Hsp70Aa #6 RNA was used in each reaction and 1.5 μM protein each from dMi-2 (1-117) lane 2, (118-238) lane 3 and (239-376) lane 4. RNA without protein was loaded for each EMSA in lane 1. The gels were stained with SYBR gold.

**Supplementary Figure 6:**
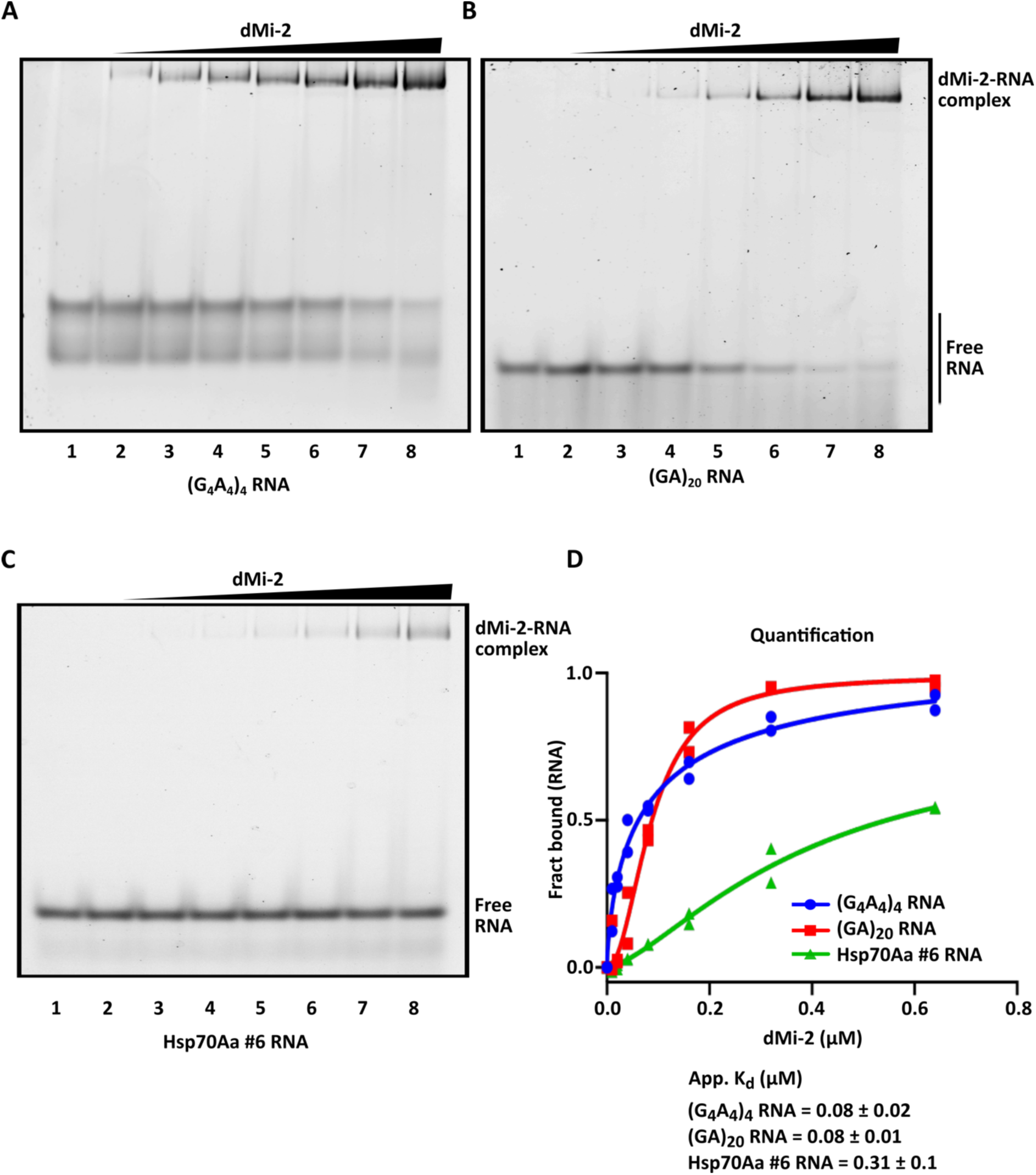
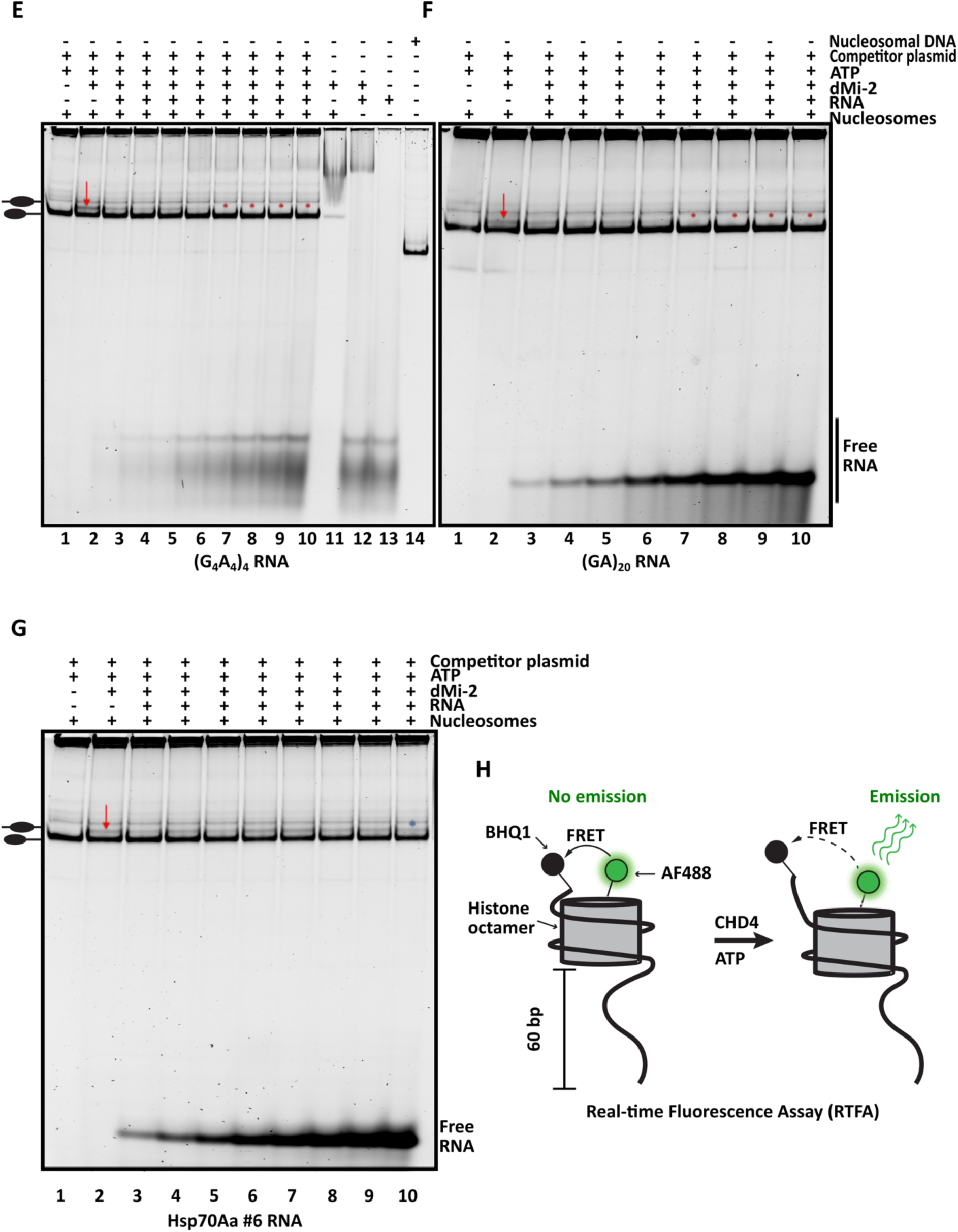
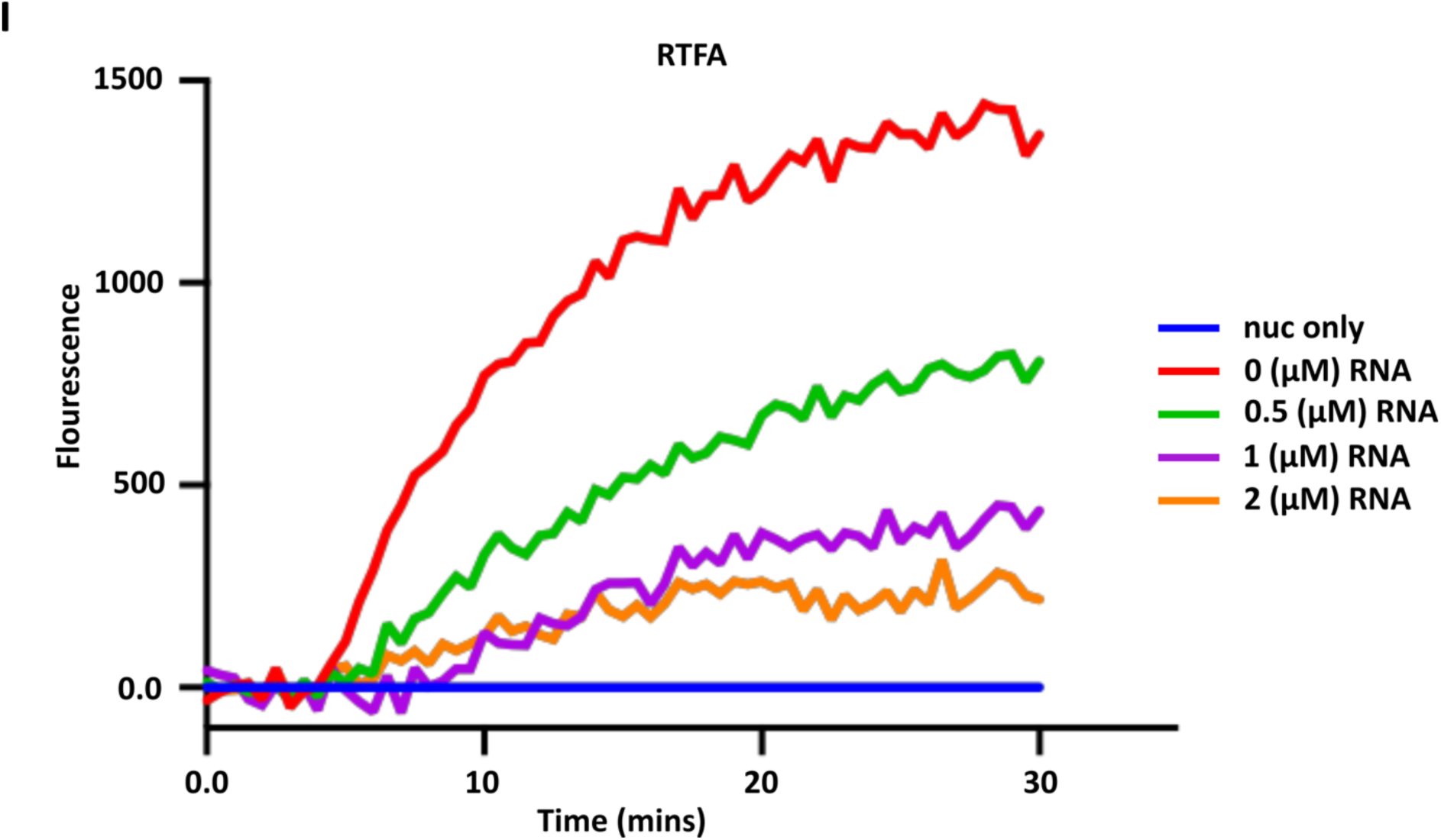
G-rich RNA inhibits dMi-2- and CHD4-mediated nucleosome remodelling *in vitro*. EMSA of **(A)** (G_4_A_4_)_4_ RNA, **(B)** (GA)_20_ RNA and **(C)** Hsp70Aa #6 RNA. 250 nM RNA was used in each reaction and dMi-2 titrated from 10 to 640 nM, lanes 2-8, each. The gels were stained with SYBR gold. RNA without protein was loaded in lanes 1 and denoted on the left. Representative images of at least two independent biological experiments. **(D)** Quantification of the EMSAs shown in (A-C). Apparent dissociation constant (app. K_d_) was calculated using Hill Slope equation (Blue: (G_4_A_4_)_4_ RNA, Red: (GA)_20_ RNA and Green: Hsp70Aa #6 RNA). Individual data points of at least two independent biological replicates are plotted. Nucleosome remodelling assays with titrations of different RNAs **(E)** (G_4_A_4_)_4_ RNA, **(F)** (GA)_20_ RNA and **(G)** Hsp70Aa #6 RNA. 15 nM of the end-positioned mononucleosomes with 80 bp of flanking DNA were used in each experiment as denoted by plus sign, except in controls (minus signs on the top). End-positioned mononucleosomes, lanes 1, represented on the left. 50 nM dMi-2 was added in each reaction (plus sign), except in controls (minus sign on the top). Remodelled nucleosome (represented on the left, can be observed in lanes 2-6 of (E) and (F) and lanes 2-10 in (G). Free RNA in each gel can be observed at the bottom. Reactions were loaded on native polyacrylamide gels, stained with SYBR gold stain. Red arrows point out the remodelled nucleosome band and red asterisk marks in lanes 7-10 of (E) and (F) point out the inhibition of remodelling. Blue asterisk mark in lane 10 of (G) points out remodelled nucleosome even at the highest concentration of Hsp70Aa #6 RNA. Representative images of at least three independent biological experiments. **(H)** Schematic representation of the end-positioned mononucleosomes (0w60) used in Real Time Fluorescence Assay (RTFA). The histone octamer is labelled with a fluorophore AF488 (in green). The nucleosomal DNA is end-labelled with a fluorescence quencher BHQ1 (in black). When the nucleosome is end-positioned, the quencher remains in close proximity to the fluorophore and no FRET signal is detected (left panel). When CHD4, in presence of ATP, remodells the nucleosome, the octamer is moved away from the quencher and the fluorescence is detected (right panel). **(I)** RTFA, as detailed in (H), was used to measure the inhibition of remodelling function of CHD4 in presence of different amounts of (GA)_20_ RNA, 0.5 µM (green), 1.0 µM (purple) and 2.0 µM (orange). RTFA signal in nucleosome only (blue) and in the absence of RNA (red) was measured as controls. RTFA signals were measured at 37 °C for 30 minutes (x-axis), with ATP being injected at 5 min time point. Averaged data from three experiments was plotted.

